# Dual E3 ligase recruitment by monovalent degraders enables redundant and tuneable degradation of SMARCA2/4

**DOI:** 10.1101/2025.08.04.668513

**Authors:** Valentina A. Spiteri, Dmitri Segal, Alejandro Correa-Sáez, Kentaro Iso, Ryan Casement, Miquel Muñoz i Ordoño, Mark A. Nakasone, Gajanan Sathe, Caroline Schätz, Hannah E. Peters, Mark Doward, Lisa Kainacher, Angus D. Cowan, Alessio Ciulli, Georg E. Winter

**Author notes:** These authors contributed equally: Valentina A. Spiteri, Dmitri Segal, Alejandro Correa-Sáez.

## Abstract

Proteolysis-Targeting Chimeras (PROTACs) and Molecular Glue Degraders (MGDs) canonically target proteins for degradation by recruiting them to a single E3 ligase complex. While heterotrivalent PROTACs that can co-opt multiple E3 ligase complexes have been described, to our knowledge all MGDs reported to date are dependent on a single E3. Here, using orthogonal genetic screening, biophysical and structural analyses, we show that a monovalent MGD can covalently recruit CUL4^DCAF16^ and CRL1^FBXO22^ in a parallel and redundant manner to degrade SMARCA2/4. Deep mutational scanning identifies a single cysteine (Cys173) in DCAF16 essential for degrader activity, and intact protein MS confirms covalent adduct at this site. The cryo-EM structure of the DCAF16:SMARCA2:degrader ternary complex reveals a unique binding mode and a distinct interface of neo-interactions, providing insights into degrader specificity. We demonstrate that E3 ligase dependency can be tuned both chemically and genetically. Minimal alterations to the compound’s “degradation tail” switches ligase preference from DCAF16 to FBXO22, while a single L59W mutation on DCAF16 is sufficient to drive DCAF16 engagement for otherwise FBXO22-dependent compounds. These results establish a molecular and structural framework for the design of tuneable dual glue degraders that could mitigate challenges from resistance mechanisms in degrader therapies.

## Main

Targeted protein degradation (TPD) is a therapeutic modality that can induce potent and selective depletion of protein levels via a catalytic mode of action (1–4). It relies on small-molecule degraders that function by recruiting a target protein to an E3 ubiquitin ligase, triggering protein ubiquitination and subsequent proteasomal degradation. Degraders are typically categorised into either bifunctional molecules known as proteolysis targeting chimeras (PROTACs), or monovalent compounds known as molecular glue degraders (MGDs). PROTACs consist of two ligands that can individually bind the E3 ligase and the target, which are connected by a linker. In contrast, MGDs typically bind either the E3 ligase or the target protein to create a composite protein-ligand interface that mediates recruitment of the other protein into a ternary complex (5). Most PROTACs, including all clinically evaluated compounds, co-opt the E3 ligases substrate receptors cereblon (CRBN) or von Hippel-Lindau (VHL) (2, 6). By leveraging new E3 ligase substrate receptor ligands by chemoproteomics approaches and high-throughput phenotypic screening additional E3 ligases have been unlocked for PROTAC design (7–12).

In contrast to the rational design of PROTACs, MGD discovery frequently relies on target-agnostic, phenotypic approaches followed by in-depth mechanistic validation studies (13–19). Recently, successful target-focused MGD discovery campaigns have been reported, which are based on attaching reactive handles, termed degradation tails, to the solvent-exposed region of the target ligand. This strategy was successfully applied to identify and characterize degraders that recruit additional E3 ligases substrate receptors including DCAF16 (11, 20–24), DCAF11 (23, 25), and FBXO22 (26, 27). To date, all drug-like degraders function via a single E3 ligase. In theory, degraders that depend on more than a single ligase would enable more resilient on-target activity and delay or potentially overcome E3 ligase driven resistance mechanisms (28–32). In a proof of concept, a dual ligase strategy has been described by us and others using heterotrivalent PROTACs designed to recruit both VHL and CRBN with a single compound to facilitate the degradation of a target protein (33). Whether drug-like MGDs could achieve a similar dual E3 ligase activity remains to be demonstrated.

Here, we mechanistically characterise a suite of monovalent SMARCA2/4 ligands that work as degraders using genetic screens, biophysical assays, and structural deconvolution. We uncover the first example of a monovalent glue that recruits two distinct E3 ligase complexes, a primary driver, CRL4^DCAF16^, and a secondary contributor, CRL1^FBXO22^, using the same degradation tail in a functionally redundant manner. Deep mutational scanning, cryo-electron microscopy (cryo-EM) and intact-mass spectrometry (MS) reveal covalent engagement of DCAF16 via cysteine 173. Our cryo-EM structure illuminates the modification in detail, further revealing a previously unresolved loop in DCAF16 that engages the degrader, leading to a distinct orientation of the SMARCA2 bromodomain relative to DCAF16-dependent BRD4 glues (34, 35). Finally, we demonstrate that DCAF16 dependency can be chemically and genetically fine-tuned, enabling a framework for the rational design of MGD with tunable E3 ligase profiles.

## Results

### Compound 1 degrades SMARCA2/4 via FBXO22 and DCAF16

A recent patent filing (WO2023018648) reported a SMARCA2/4 degrader, herein referred to as Compound 1 (**1**) (Figure 1a), which comprises a SMARCA2/4 bromodomain ligand bearing a propargyl-azepane tail group (36). **1** was found to induce degradation of SMARCA2/4 in HEK293T and HCT-116 cells (Extended Data Figure 1a, b, Figure 1f) and quantitative mass spectrometry proteomics confirmed degradation selectivity towards SMARCA2/4 and PBRM1 (Figure 1b, Supplementary Table 1) (37). In contrast, **1\***, a close analogue of **1** lacking the propargyl-azepane tail, had no degradation activity (Extended Data Figure 1c), indicating that this moiety is essential for degradation. SMARCA2 degradation was prevented by inhibiting the ubiquitin-activating enzyme UBA1 (TAK243), neddylation (MLN4924), and the proteasome (Carfilzomib, Figure 1c), consistent with Cullin Ring Ligase (CRL) mediated degradation. In support of this, NanoBRET ubiquitination assays in live cells (Extended Data Figure 1d) showed dose- and time-dependent increases in SMARCA2/4 ubiquitination after treatment with **1**, which was abolished by MLN4924 treatment (Extended Data Figure 1 e-f).

**Figure 1.**
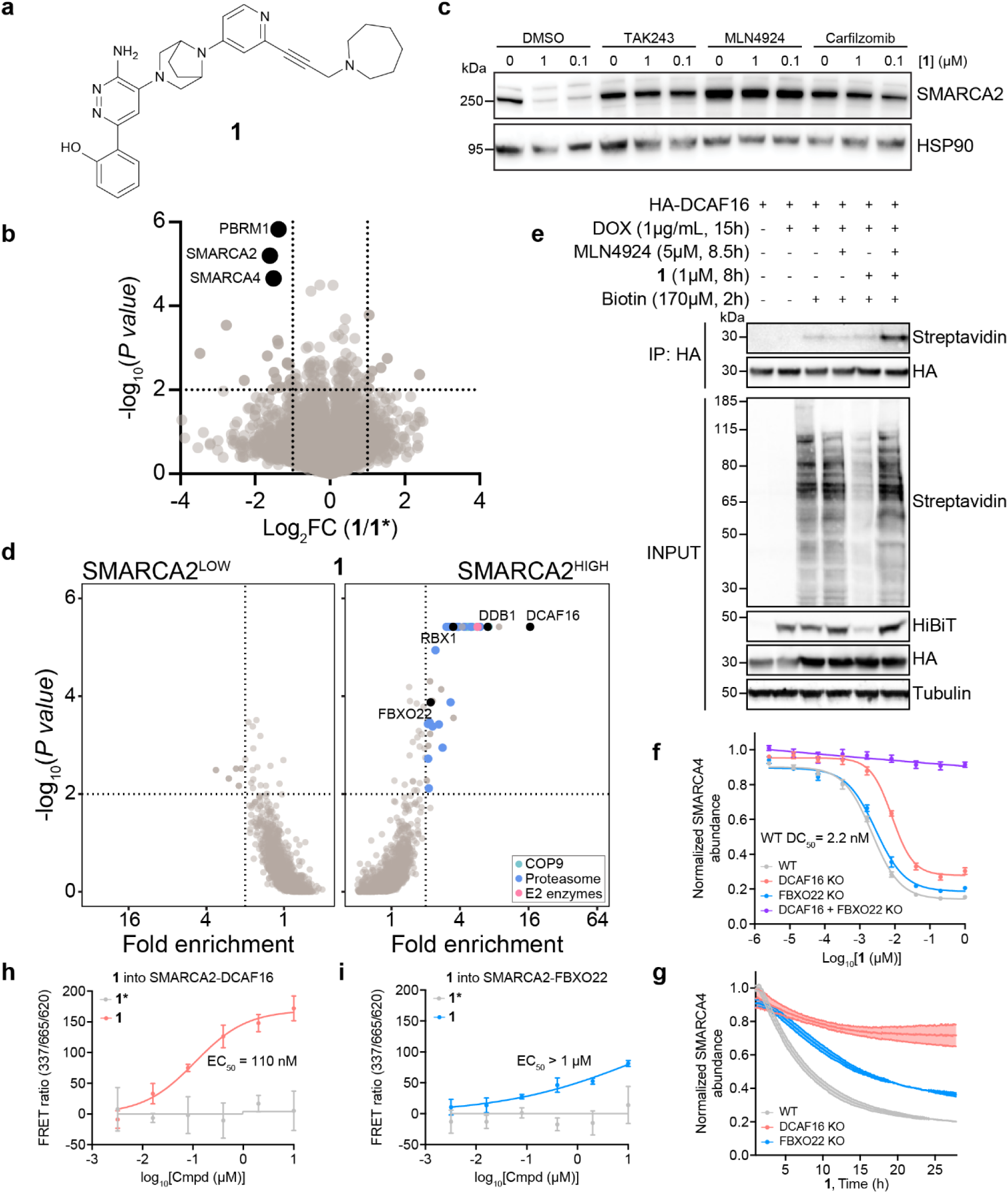
Compound 1 recruits both DCAF16 and FBXO22 to degrade SMARCA2/4. **a**, Struture of Compound 1 (1). **b**, Whole-proteome changes after degrader treatment. Quantitative proteomics in HEK293 cells was performed after 12 hours of treatment with **1** (1 μM) or the inactive control **1\*** (1 μM). Log2-transformed fold change and −log10-transformed Benjamini–Hochberg adjusted one-way analysis of variance (ANOVA) P value of **1** compared with **1\***. n = 3 biological replicates. **c**, Degradation rescue by proteasome and neddylation inhibitors. HEK293T cells were co-treated with **1** (0.1 or 1 μM) and DMSO, TAK243, MLN4924, or Carfilzomib (0.5 μM) for 18 hours. SMARCA2 was quantified by immunoblot. **d**, FACS-based CRISPR screen for SMARCA2 stability. KBM7 iCas9 SMARCA2 bromodomain stability reporter cells were transduced with a ubiquitin-proteasome-system-focused sgRNA library, treated with **1** (0.1 μM) for 24 hours, and sorted based on SMARCA-BFP levels. Fold changes and P values of the SMARCA2^HIGH^ and SMARCA2^LOW^ populations were calculated by comparison with the SMARCA2^MID^ population using robust rank aggregation algorithm (MAGeCK). Significant hits: fold enrichment ≥ 2 and -log_10_P values ≥ 2 (dotted lines). n = 2 biological replicates. **e**, Biotin proximity labelling of DCAF16. Flp-In T-REx HEK293 cells transiently expressing HA-tagged DCAF16 and doxycycline-inducible miniTurboID-SMARCA2-BD were pretreated with 5 μM MLN4924 for 30 min, followed by treatment with **1** (1 μM) and 170 μM biotin for 8 and 2 hours, respectively. HA-tagged DCAF16 was immunoprecipitated and biotinylated proteins were detected via streptavidin immunobloting. **f**, HiBiT endpoint degradation assay. HEK293T cells expressing endogenous SMARCA4-HiBiT with single and double KOs were treated with a serial dilution series of **1** for 16 hours. SMARCA4 levels were quantified using the HiBiT lytic detection system. Data represent mean ± SEM, n = 6 biological replicates. **g**, HiBiT kinetic degradation assay. HEK293T cells expressing endogenous SMARCA4-HiBiT with single KOs were treated with **1** (1 μM) and luminescence was measured every 5 minutes to monitor real-time degradation. Data represent mean ± SEM, n = 3 biological replicates. **h**, **i**, TR-FRET ternary complex formation assay. Anti-GST-europiym bound to GST-SMARCA2 Bromodomain (BD) was inclubated with Cy5-labelled DCAF16-DDB1 (ΔBPB)-DDA1 (**h**) or Cy5-labelled FBXO22-SKP1 (**i**) and increasing concentrations of **1** or **1\***. Data represent mean ± SEM, n = 3 biological replicates.

To identify the CRL required for the activity of **1**, we set up a time-controlled, fluorescence-activated cell sorting (FACS)-based CRISPR screen leveraging a dual fluorescence SMARCA2-Bromodomain (BD) stability reporter and a ubiquitin proteasomal system (UPS)-focused sgRNA library (Extended Data Figure 2a) (26). Following treatment with **1**, the substrate receptor DCAF16 alongside other CRL4^DCAF16^ core complex members, DDB1 and RBX1, emerged as top hits. As expected, we also identified components of the 20S proteasome and COP9 signalosome (Figure 1d, Supplementary Table 2), again indicative of a CRL- and proteasome-dependent mechanism. To validate the role of DCAF16 in driving **1**-mediated degradation of SMARCA2, we employed a proximity-dependent biotin identification (BioID) approach based on miniTurboID-tagged SMARCA2 BD and HA-tagged DCAF16 (38). Treatment with **1** resulted in enhanced biotinylation of DCAF16, demonstrating induced proximity of both proteins upon compound treatment (Figure 1e). However, to our surprise, knockout of DCAF16 failed to fully abrogate degradation of the SMARCA2 BD stability reporter by **1** (Extended Data Figure 2b). Thus, we wondered whether additional factors are redundantly involved in SMARCA 2/4 degradation induced by **1**. To that end, we conducted SMARCA2 BD proximity labelling via unbiased proteomics, demonstrating a significant (p value = 3.16×10^−4^, Fold-change = 2.23) increase in biotinylation of the CRL1 substrate receptor FBXO22 in the presence of **1** (Extended Data Figure 2c, Supplementary Table 3). Intriguingly, even though genetic loss of function approaches frequently fail to resolve genetic redundancies, FBXO22 was also identified as a weak hit in the FACS-based CRISPR screen (Figure 1d, Supplementary Table 2), suggesting the involvement of two unrelated ligases, CRL4^DCAF16^ and CRL1^FBXO22^ in the SMARCA2/4 degradation mediated by **1**.

To validate this hypothesis, we generated single and double knock out (KO) cell lines for both DCAF16 and FBXO22. By western blotting and NanoBRET ubiquitination, we found that only the double KO was able to completely ablate ubiquitination and degradation of SMARCA2/4 mediated by **1** (Extended Data Figure 2d-e). A similar trend was observed in HiBiT degradation assays for endogenously tagged SMARCA4 in dose– response (WT DC_50_ = 2.2 nM, Figure 1f). Moreover, single DCAF16 KO and FBXO22 KO cell lines showed slowed but not abrogated degradation in time-course experiments (Figure 1g). Together, these experiments suggest that both ligases are functionally required to elicit SMARCA2/4 degradation by **1**. To assess if both ligases also form a ternary complex with SMARCA2 BD in the presence of **1** *in vitro,* we developed a time-resolved fluorescence resonance energy transfer (TR-FRET) assay. Indeed, **1** showed dose-dependent ternary complex formation with both CRLs that was more potent for DCAF16 (EC_50_= 110 nM) compared to FBXO22 (EC50 > 1 μM). No complex formation was observed in the presence of the non-degrading analogue **1\*** (Figure 1h-i). Together, our data is consistent with an unprecedented glue degrader mechanism that co-opts two distinct E3 ligase complexes: CRL4^DCAF16^ as the primary driver mediating degradation, and CRL1^FBXO22^ as a secondary modulator.

### Compound 1 covalently adducts DCAF16 at cysteine 173

To identify functional hotspots on DCAF16 we performed deep mutational scanning (DMS) of the full-length protein. This enabled us to systematically assess the effects of every possible point substitution (4300 variants in total) in DCAF16 on SMARCA2 BD degradation mediated by **1**. We performed the screen in DCAF16 and FBXO22 double KO HEK293T cells to eliminate background interference from endogenous ligases. To distinguish context-specific effects from general defects caused by mutations that disrupt overall DCAF16 stability, we concurrently monitored BRD4 BD1-BD2 degradation mediated by two previously described, DCAF16-dependent BRD4 BD1-BD2 monovalent glue degraders, MMH2 and GNE-0011 (34, 39). Comparative analysis between degrader-specific variants revealed that substitutions in C173 profoundly disrupted SMARCA2 degradation (Figure 2a, Extended Data Figure 3a, b) induced by **1**. This contrasts with mutations in C58, L59, K61 and W181, which affect degradation of BRD4 BD1-BD2 mediated by MMH2 or GNE-0011, as previously shown (34). Additionally, the mutation Y62T was also shown to significantly disrupt the SMARCA2 degradation induced by **1**. The impact of both C173S and Y62T were confirmed by western blot (Figure 2b). The DMS results identify C173 as a hotspot for the mode of action of **1** that is specific over previously described DCAF16-dependent BRD4 degraders, suggesting a differentiated mechanism of ligase recognition and a potentially altered architecture of the induced ternary complex.

**Figure 2.**
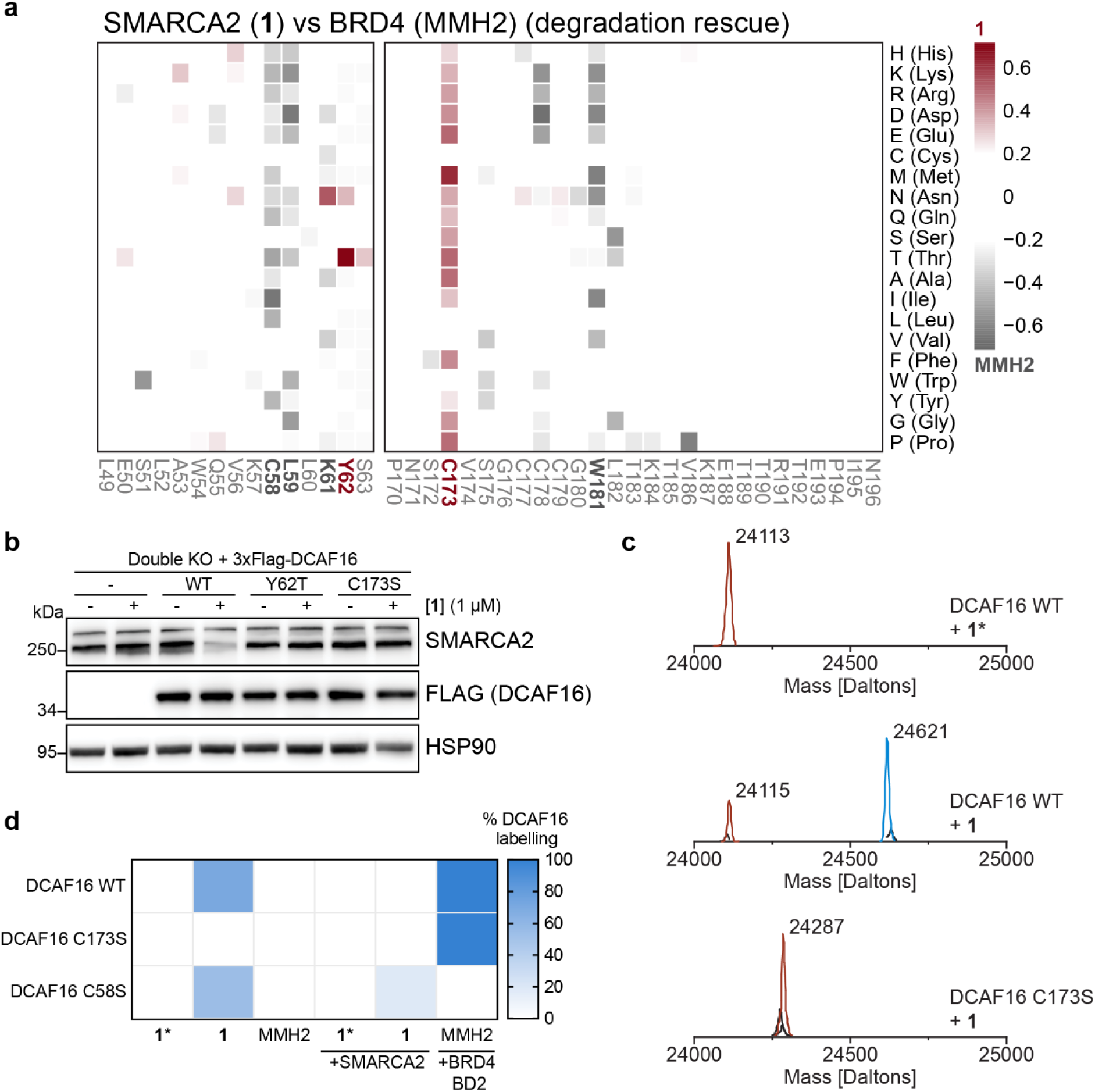
Compound 1 covalently engages with DCAF16 via Cysteine 173. **a**, Deep mutagenesis scanning (DMS) of DCAF16 for degradation rescue. DCAF16 and FBXO22 Double KO HEK293T cells stably expressing stability reporters for SMARCA2 BD or BRD4 tandem-BDs were transduced with a library of EGFP tagged DCAF16 variants containing single amino acid substitutions at every position. SMARCA2 and BRD4 stability reporter cells were treated with **1** (1 µM) and MMH2 (1 µM) for 24 hours, respectively. Heatmap depicting differential log_2_ fold enrichment of DCAF16 mutations normalized to maximum log_2_ fold changes versus unsorted control between SMARCA2 targeting compound **1** and BRD4 targeting MMH2. Data corresponds to SMARCA2^High^ and BRD4^High^ sorted populations, showing mutations that specifically prevent SMARCA2 degradation (red) and BRD4 degradation (gray). n=3 independent measurements. **b**, Validation of DMS. Double KO HEK293T cells stably expressing 3xFlag tagged DCAF16 variants were treated with **1** (1 µM) for 24 hours. SMARCA2 was quantified by immunoblot. **c**, Intact protein mass spectra for wild-type DCAF16 co-incubated with inactive control **1\*** (Top), wild-type DCAF16 co-incubated with 1 (Middle), and DCAF16 C173S mutant co-incubated with 1 (Bottom). All incubations were done with 10 µM DCAF16, 20 µM **1\*** or **1**, and left at room temperature for 18 hours. **d**, Proportion of labelled DCAF16. Heatmap depicting the proportion (%) of labelling from intact-MS for each DCAF16 variant (10 µM) in the presence of **1\***, **1**, or MMH2 (all at 20 µM). Intact-MS experiments for DCAF16 were performed with or without the presence of SMARCA2 BD or BRD4 BD2 (both at 20 µM)

Given the significance of C173, we investigated whether **1** recruits DCAF16 via a covalent mechanism. Leveraging intact protein MS, we demonstrated covalent labelling of recombinant DCAF16 WT and DCAF16 C58A by **1,** while labelling was abrogated for the DCAF16 C173S mutant (Figure 2c-d), indicating that **1** specifically covalently adducts C173. Interestingly, no labelling of DCAF16 was observed upon co-incubation with SMARCA2 BD(Figure 2d and Extended Data Figure 3c). This pattern markedly contrasts with MMH2 where labelling of DCAF16 depends on co-incubation with BRD4 BD2, (Figure 2d and Extended Data Figure 3d). This suggests that, unlike the template-assisted covalent mechanism of MMH2, labelling of DCAF16 is not templated, but rather hindered by SMARCA2 BD, at least at the μM-range of concentrations used in these experiments (34). Collectively, the biophysical and unbiased genetics data show that **1** mediates the formation of a ternary complex between the SMARCA2/4 bromodomains and its driver ligase DCAF16 via covalent modification of C173 on DCAF16.

### Structure of SMARCA2 and Compound 1 glued to DCAF16

To gain a structural understanding of the mechanism underpinning **1**-induced SMARCA2 BD degradation, we solved the structure of the ternary complex formed between SMARCA2 BD, **1** and DCAF16:DDB1(ΔBPB):DDA1 by cryo-EM with a resolution of approximately 3.47 Å (Figure 3a, Extended Data Figures 4-5 and Extended Data Table 2). We observed a clear density for SMARCA2 BD sitting on top of DCAF16 in an orientation tilted slightly relative to the core of the ligase complex. We detected density corresponding to a back loop (residues E164-C177) (Figure 3b) that was unresolved in previously published structures of DCAF16 (PDB IDs: 8OV6 and 8G46) (34, 35). Strikingly, a clear continuous density is seen between C173 that resides on this loop and the compound, consistent with formation of a covalent bond (Figure 3b-c). The position of the SMARCA2 BD is distinct relative to the MMH2-induced position of BRD4 BD2 (PDB: 8G46). SMARCA2 adopts a backward-tilted orientation towards the DCAF16 loop region, in contrast to the forward tilt observed for BRD4 BD2 (Extended Data Figure 6). This difference likely arises from covalent labelling of distinct cysteine residues on DCAF16 and highlights the structural flexibility of DCAF16 in accommodating the same bromodomain fold in distinct orientations, enabling adaptable degrader engagement across diverse targets.

**Figure 3.**
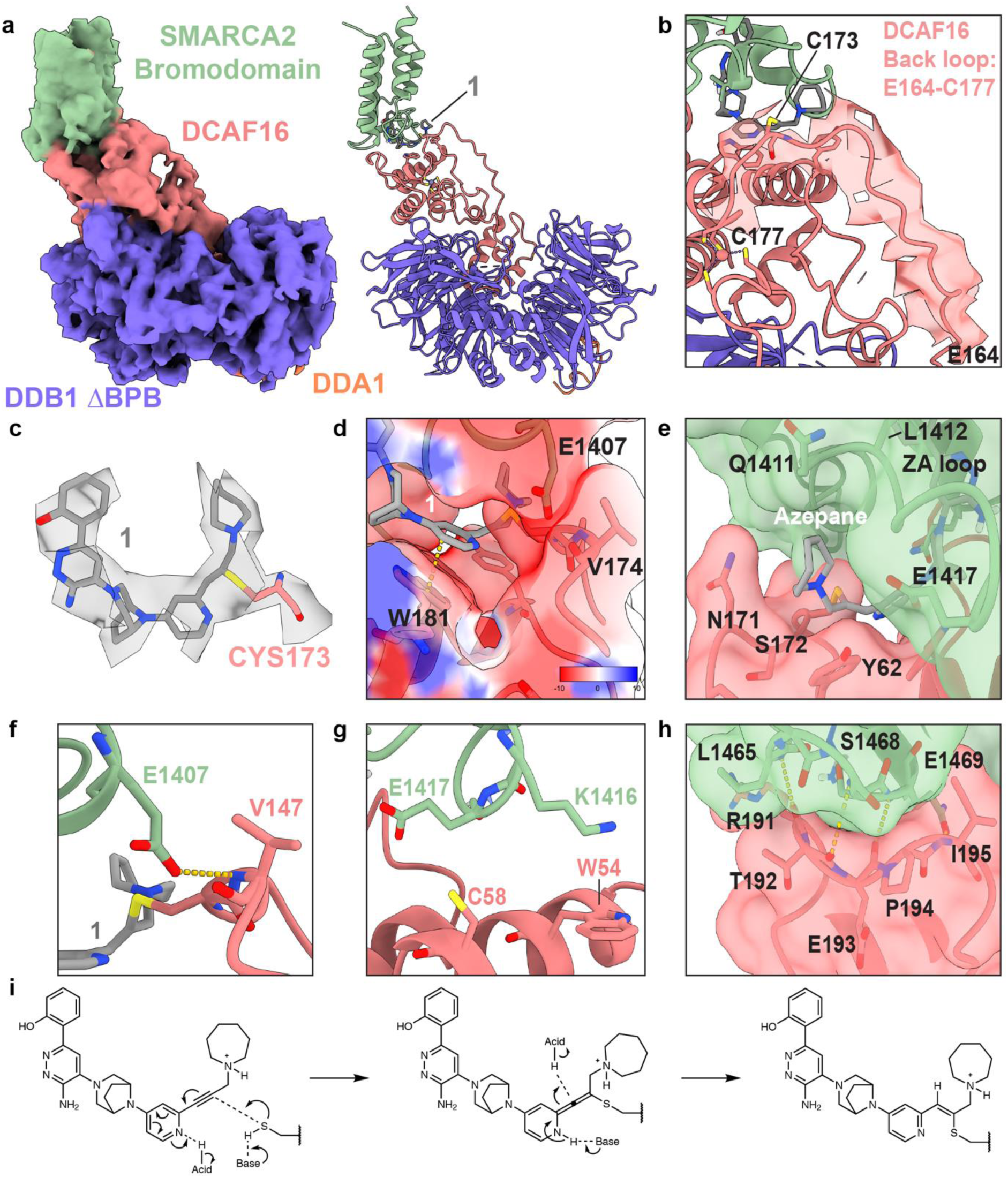
Cryo-EM structure of the ternary complex formed between compound 1, SMARCA2, and DCAF16. **a**, Structure of the complex between DCAF16 (red), DDB1(ΔBPB) (purple), DDA1 (orange), and SMARCA2 BD (green), glued by 1 (gray), solved by cryo-EM at a global resolution of 3.47 Å. Electron density shown on the left and model on the right. **b**, Electron density of the newly resolved back loop (E164-C177) of DCAF16 that harbours C173 and is covalently engaged by **1**. **c**, Electron density around **1** and C173. **d**, Electrostatic surface potential showing area of negative potential (red) around the pyridine ring of **1** due to presence of the V174 backbone carbonyl (DCAF16) and E1407 side chain (SMARCA2). The pyridine of **1** stacks with W181 of DCAF16. **e**, The azepane ring of **1** nestles into a pocket formed by residues Q1411–L1412 of SMARCA2 and P170–S172 of DCAF16, while also stacking with Y62 of DCAF16. **f**, Protein-protein interaction between the side chain carboxylate of E1407 on SMARCA2 and the backbone amide of V174 on DCAF16 **g**, K1416 and E1417 from the ZA loop of SMARCA2 BD point towards W54 and C58 in DCAF16 **h**, SMARCA2 BD BC loop residues, L1465-E1469, hydrogen bond with residues R191-I195 on a DCAF16 loop. **i**, Schematic showing the predicted covalent attachment of **1** to the side chain of C173.

The region of 1 that is solvent-exposed when in a binary complex with SMARCA2 BD forms extensive direct interactions with DCAF16, in addition to the covalent bond with C173. The pyridine group of **1** engages in a π stacking interaction with W181 of DCAF16 (Figure 3d), directing the degradation tail to insert at the interface between the ZA-loop of the SMARCA2 BD and DCAF16 (Figure 3e). Specifically, the terminal azepane ring fits snugly into a pocket defined by Q1411-L1412 on the bromodomain, P170-S172 on the DCAF16 loop, and Y62 on DCAF16 (Figure 3e). DCAF16 Y62 was also identified as functionally relevant via DMS (Figure 2a). In addition to the direct interactions formed between the degrader and DCAF16, we observe extensive induced protein-protein contacts at the ternary complex interface. These contacts bury a total surface area of 1057 Å^2^ comparable to other PROTAC and MGD ternary structures, ranging from 680-2672 Å^2^ (5, 40, 41)). The pyridine of the degradation tail sits in a negatively charged pocket including the E1407 sidechain carboxylate from the ZA loop of SMARCA2 and the backbone carbonyl of V174 on DCAF16 (Figure 3d). This environment suggests the pyridine is likely protonated, further activating the alkyne for nucleophilic attack. A molecular dynamics simulation of the ternary complex revealed that Ser175 on DCAF16 can hydrogen bond with the nitrogen in the pyridine ring of **1** (Extended Data Figure 6b). Moreover, several protein-protein interactions stabilise the ternary complex, including a hydrogen bond between the side chain carboxylate of E1407 on SMARCA2 and the backbone amide of V174 on DCAF16 (Figure 3f). On the other side of the complex, K1416 and E1417 from the ZA loop of SMARCA2 BD point towards a helix in DCAF16 containing C58 that has been covalently engaged by previously reported MGDs (Figure 3g) (34, 42). Finally, at the front end of the interface, SMARCA2 BD residues from L1465 to E1469 in the BC loop, form hydrogen bonds and exhibit close sidechain packing with the R191-I195 loop of DCAF16 (Figure 3h). The direct degrader–ligase contacts and induced protein–protein interactions orient SMARCA2 BD and DCAF16 into a compact ternary complex, stabilized by covalent bonding, hydrogen bonds, π-stacking, and extensive surface burial across several interacting loop regions.

Taken together, the cryo-EM structure unveils in atomic detail the covalent modification of DCAF16 at C173 by **1**, and recruitment of SMARCA2. The cryo-EM structure together with the biophysical and mutagenesis data supports proposal of a mechanism for covalent labelling of DCAF16 by **1,** where C173 undergoes a nucleophilic addition to the electron-deficient acetylene moiety of the compound, which is activated by the electron-poor pyridine ring. This leads to regio- and stereo-specific covalent bond formation and recruitment of the SMARCA2/4 bromodomain to the modified DCAF16 (Figure 3i).

### Chemical tunability of E3 ligase selectivity

To investigate how changes to the degradation tail might impact ligase recruitment, we aimed to characterize Compound 2 (**2**), another SMARCA2/4 degrader from the same patent (Figure 4a, Extended Data Figure 7a) (WO2023018648) (36). In contrast to **1**, with **2** the azepane group was changed to a piperazine and an extra carbon was added to the linker. To determine E3 ligase substrate receptor preference of **2**, we conducted another FACS-based CRISPR/Cas9 screen with the SMARCA2 BD stability reporter, which revealed a complete reversal in driver ligase preference to CRL1^FBXO22^ compared to **1** (Figure 4b, Supplementary Table 2). Moreover, unbiased BioID proximity labelling revealed that FBXO22 was among the few proteins significantly enriched upon treatment with **2** (Figure 4c and Extended Data Table 3). The preferential proximity between SMARCA2 BD and FBXO22 induced by **1** compared to **2** was further validated by BioID labelling coupled to western blotting, which resulted in a higher degree of FBXO22 biotinylation (Extended Data Figure 7b). Western blotting and NanoBRET ubiquitination assays confirmed that single FBXO22 KO is sufficient to abrogate SMARCA2 ubiquitination and degradation induced by **2** (Figure 4d-e). Finally, the driver ligase switch was confirmed by SMARCA4-HiBiT degradation assays in both dose–response (Figure 4f) and time-course experiments (Figure 4g), showing DCAF16 dependence only at higher concentrations. Together, these findings demonstrate that subtle structural modifications to the degradation tail are sufficient to reprogram ligase engagement from CRL4^DCAF16^ to CRL1^FBXO22^, highlighting a chemically tunable mechanism for modulating E3 ligase selectivity in monovalent degraders.

**Figure 4.**
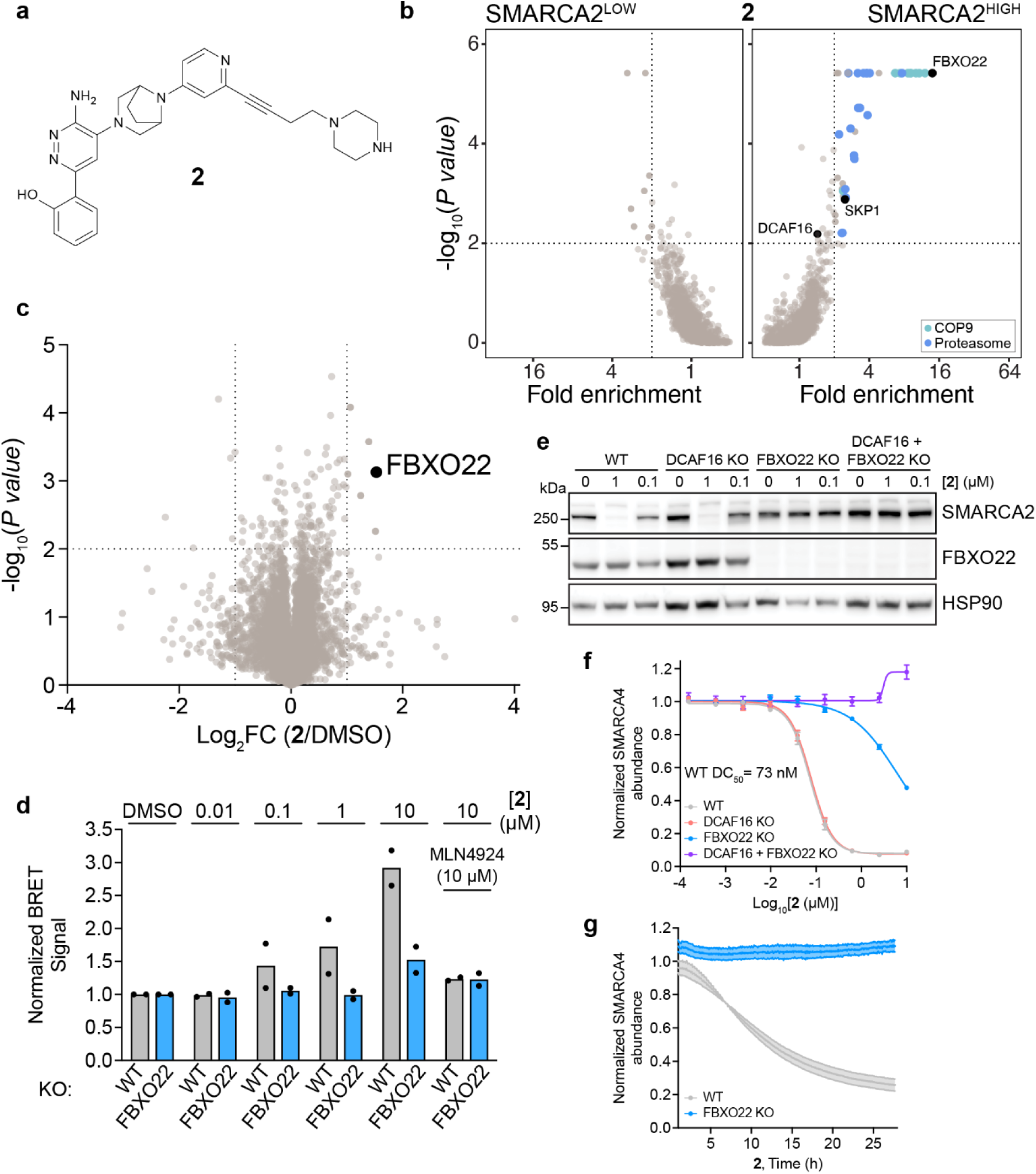
Compound 2 exhibits a driver ligase-switch from DCAF16 to FBXO22. **a**, Structure of Compound **2**. **b**, FACS-based CRISPR screen for SMARCA2 stability. KBM7 iCas9 SMARCA2 bromodomain stability reporter cells were transduced with a ubiquitin-proteasome-system-focused sgRNA library, treated with **2** (1 µM) for 24 hours, and sorted based on SMARCA-BFP levels. Fold changes and p values of the SMARCA2^HIGH^ and SMARCA2^LOW^ populations were calculated by comparison with the SMARCA2^MID^ population using robust rank aggregation algorithm (MAGeCK). Significant hits: fold enrichment ≥ 2 and –log_10_P values ≥ 2 (dotted lines). n = 2 biological replicates. **c**, BioID analysis with **2** treatment. Proximity labelling in Flp-In T-REx HEK293 miniTurboID-SMARCA2-BD inducible cell line. Cells were treated with 1 µg/ml doxycycline for a total of 20.5 hours. 12 hours after induction, cells were pre-treated with 10 µM MLN4924 for 30 mins, followed by addition of **2** (10 µM). Volcano plot of BioID results showing log_2_ fold change (FC) and −log_10_adjusted P values derived from normalized protein spectral counts and compared with DMSO treatment. Scoring window (P value < 0.01, log_2_ FC > 1) is indicated, n = 3 biological replicates. **d**, NanoBRET ubiquitination assay. WT or FBXO22 KO HEK293T SMARCA4-HiBit cells transfected with LgBiT and HaloTag-Ubiquitin were treated with DMSO or increasing concentrations of **2** (± 10 µM MLN4924, 1 hour pre-treatment) for 24 hours. Data represent mean ± SD, n = 2 biological replicates. **e**, Genetic rescue of SMARCA2 degradation. WT, DCAF16 KO, FBXO22 KO, and double KO HEK293T cells were treated with DMSO or **2** (0.1 and 1 µM) for 24 hours. SMARCA2 was quantified by immunoblot. **f**, HiBiT endpoint degradation assay. HEK293T SMARCA4-HiBit cells with single and double KOs were treated with a serial dilution series of **2** for 24 hours. SMARCA4 levels were quantified using the HiBiT lytic detection system. Data represent mean ± SEM, n = 6 biological replicates. **g**, HiBiT kinetic degradation assay. WT and FBXO22 KO SMARCA4-HiBiT cells were treated with **2** (1 µM) and luminescence was measured every 5 minutes to monitor real-time degradation. Data represent mean ± SEM, n = 3 biological replicates.

Given that **1** engages DCAF16 via a covalent mechanism, we asked whether the same might be true for the driver ligase of **2**, FBXO22. Therefore, we mutated surface-exposed cysteine residues of FBXO22 and measured their impact on **2-**induced SMARCA2 degradation. Mutations in C228 and C326 both abolished SMARCA2 degradation induced by **2** (Extended Data Figure 7c), highlighting a set of previously tractable cysteines in FBXO22 used for TPD (26, 43). **1** was also shown to rely on the same set of cysteines (Extended Data Figure 7d), suggesting a shared mechanism of FBXO22 engagement between both compounds. The same patent and a recent publication also described a potent FBXO22-dependent degrader, G-6599, hereafter referred to as compound 3 (**3**) (Extended Data Figure 8a-b) (44). **3** is almost identical to **1** and only features an additional methylene group between the alkyne and azepane groups. Consistent with **1,** a simultaneous knockout of DCAF16 and FBXO22 was needed for full rescue of SMARCA2/4 degradation (Extended Data Figure 8c-d), suggesting that **3** is also dual-ligase dependent. Similar to **2**, we observed that **3** also undergoes a driver ligase switch to become more CRL1^FBXO22^ dependent (Extended Data Figure 8d). To understand if the driver ligase switch observed with **3** alters the molecular recognition of DCAF16, we turned to our DMS screening data. Similar to **1**, C173 and Y62T mutations resulted in the greatest loss of SMARCA2 degradation after treatment with **3** (Extended Data Figure 8e-f, Supplementary table 4). We confirmed C173 is also covalently labelled by **3** using intact protein MS, where DCAF16 WT was labelled in the presence of **3**, in contrast to DCAF16 C173S mutant (Extended Data Figure 8g). These findings underscore how even minimal chemical modifications, such as the addition of a single methylene group, can be sufficient to induce a switch in primary ligase dependency.

To further explore features in the degradation tail critical for ligase engagement and degrader activity, we synthesized and evaluated a set of analogues of **1**. Modification of the azepane ring on **1** to either the smaller and less basic piperazine (**4**) or a cycloheptane (**5**) resulted in a complete loss of SMARCA2/4 degradation in HiBiT assays and Western blotting (Extended Data Figures 9a-e). This highlights that very subtle changes in ring size or basicity profile disrupt degrader function and fail to support recruitment of either DCAF16 or FBXO22 in cells. We used a glutathione reactivity assay to assess whether the loss in activity correlated with changes in electrophilicity. **1** showed the highest reactivity, followed by **3, 4,** and **2,** whereas **5** displayed no detectable reactivity (Extended Data Figure 9f). These findings emphasize the critical role of the azepane moiety in maintaining both the chemical reactivity and structural features necessary for selective engagement of DCAF16, highlighting how small modifications can disrupt covalent binding and degradability.

### Genetic Tunability

After having established that chemical modifications offer a means to fine-tune ligase selectivity, we next asked whether similar control could be achieved through genetic modulation. We again utilized FACS-based DCAF16 DMS in double KO HEK293T cells to genetically adapt the topology of the DCAF16 surface to induce an enhancement/gain-of-function in SMARCA2 BD degradation by **2** and **3**. To distinguish specific mutations from those that simply stabilize DCAF16, we concurrently evaluated how mutations affect BRD4 BD1-BD2 degradation by the DCAF16-dependent degrader GNE-0011. DMS results for **2** and **3** indicated that substitutions at several residues including S51, W54, L59, V174, Y165, G180, E188, and I195 were associated with increased SMARCA2 degradation upon treatment (Figure 5a, Extended Data Fig. 10a-b, Supplementary Table 4). Based on these findings, we expressed the corresponding mutant variants and subsequently examined SMARCA2 degradation using western blot analysis in the presence of **2** (Extended Data Figure 10c). Among the variants tested, L59W emerged as the only mutation capable of unambiguously promoting SMARCA2 degradation in presence of **2** and **3** compared to DCAF16 wildtype (Extended Data Fig. 10c). Consistently, L59W also caused a marked shift in SMARCA4-HiBiT degradation compared to wild-type DCAF16 using compounds **2** and **3** (Figure 5b), illustrating that ligase dependency can be genetically fine-tuned. Mechanistically, the role of the L59W mutation could be explained by its proximity to the degraders in the pocket formed by DCAF16 and SMARCA2 BD (Figure 5c). We noticed that the W181 side chain flips “out of plane” from alternative positions observed in previous DCAF16 structures (PDB IDs: 8G46 and 8OV6), in order to form its π stack with the pyridine of 1 (Figure 5d). From modelling the L59W mutation we hypothesize that W59 sits below W181 stabilising the flipped conformation of this residue and forming an extended “WW-pyridine shelf” stacking with the pyridine rings of **2** and **3**, ultimately allowing for increased DCAF16 engagement with these compounds (Figure 5d).

**Figure 5.**
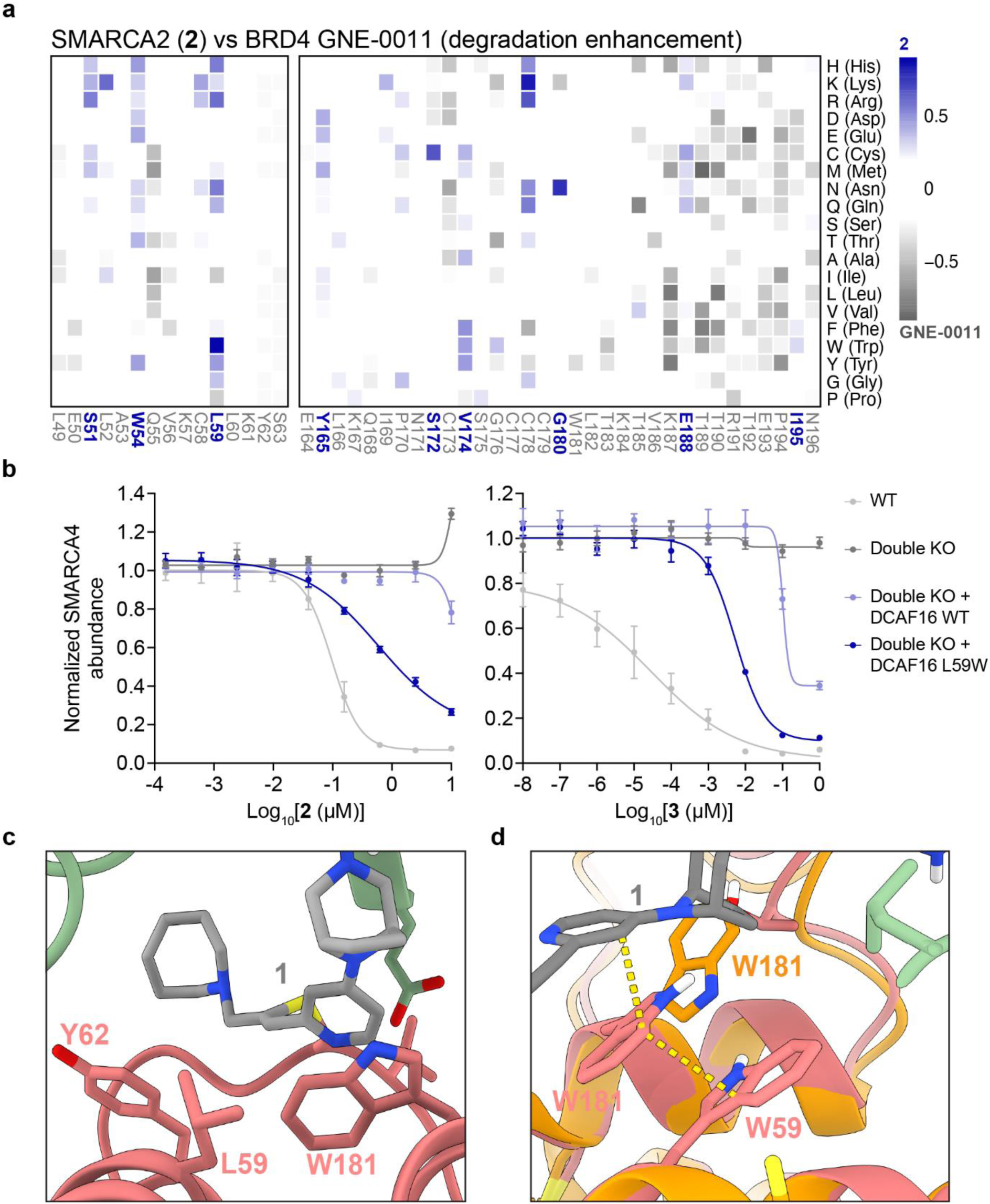
DCAF16 L59W mutation genetically fine-tunes E3 ligase dependancy of compounds 2 and 3. **a**, Deep mutagenesis scanning (DMS) of DCAF16 for degradation enhancement. DCAF16 and FBXO22 double knockout (KO) HEK293T cells stably expressing stability reporters for SMARCA2 BD or BRD4 tandem-BDs were transduced with a library of EGFP tagged DCAF16 variants containing single amino acid substitutions at every position. SMARCA2 and BRD4 stability reporter cells were treated with **2** (1 µM) and GNE-0011 (0.1 µM) for 24 hours, respectively. Heatmap depicting differential log_2_ fold enrichment of DCAF16 mutations normalized to maximum log_2_ fold changes versus unsorted control between SMARCA2 targeting **2** and BRD4 targeting GNE-0011. Data corresponds to SMARCA2^Low^ and BRD4^Low^ sorted populations, showing mutations that specifically enhance SMARCA2 degradation (blue) and BRD4 degradation (gray) n=3 independent measurements. **b**, DMS validation with HiBiT endpoint degradation assay. Wild-type (WT) and double KO HEK293T SMARCA4-HiBiT cell lines stably expressing 3xFlag tagged DCAF16 variants were treated with a serial dilution series of 2 for 24 hours (left) or 3 for 18 hours (right). SMARCA4 levels were quantified using the HiBiT lytic detection system. Data represent mean ± SEM, n = 6 biological replicates. **c**, Position of L59 of DCAF16 in the ternary complex. **d**, Model of the L59W mutation stabilising W181 that π stacks with the pyridine of **1**. The L59W model with **1** is overlaid with another DCAF16 structure (PDB ID: 8G46, orange) where W181 is flipped outward.

In sum, these findings demonstrate that, in addition to chemical tuning, ligase choice can be genetically fine-tuned through specific mutations at the degrader–ligase interface. In particular, the L59W mutation enhances DCAF16 engagement by **2** and **3**, likely by optimizing interactions within the degrader binding pocket, offering a potential rational approach to enhance DCAF16 recruitment.

## Discussion

Target-focused and ligase-agnostic discovery strategies for small-molecule degraders have been instrumental in expanding the scope of E3 ligases that can be pharmacologically co-opted, circumventing the need to develop new binding ligands for the E3 ligase substrate receptors. Applying such approaches to study SMARCA2/4 degraders featuring a unique propargyl-azepane group at the solvent-exposed exit vector, we identified the first monovalent degraders capable of recruiting two E3 ligase substrate receptors, DCAF16 and FBXO22, in a parallel and redundant fashion. The described compounds are hence distinct from monovalent degraders and bivalent PROTACs described to date, which typically recruit a single E3 ligase substrate receptor for target degradation.

A suite of previous studies has identified the structurally unrelated ligases DCAF16, FBXO22 and DCAF11 as particularly susceptible to covalent labelling. It has also been shown that, depending on the exit vector decoration of a small-molecule degrader, target protein degradation can be biased to either go via DCAF16 or DCAF11 (23). A possible explanation for the susceptibility of these ligases is the cysteine rich surfaces that may correlate to their native roles that have thus far largely remained elusive. Previous studies have outlined template-assisted covalent modification on DCAF16 for degraders of BRD4 and BRD9. Our findings indicate that the degraders described in this manuscript do not conform to this model. DCAF16 covalent modification can occur without SMARCA2/4 orienting the degrader for bond formation. This suggests that the propargyl-azepane degradation tail is sufficient to productively orient the degrader on the DCAF16 surface. We also observed that **1** is the most reactive compound in glutathione assay followed by **2** and **3** (Extended Data Figure 9f). The reactivity seems to correlate to DCAF16 dependence *in cellulo,* suggesting a connection between reactivity and ligase dependence. Further work is needed to fully probe such possible connection between ligase reactivity and ligase specificity.

Of note, previous electrophilic compounds have largely targeted cysteines 58 and 178 in more structured or conformationally rigid regions of DCAF16 (11, 20, 34, 42, 45). In contrast, the cryo-EM structure of the DCAF16:DDB1 (**Δ**BPB):DDA1, SMARCA2 BD and **1** reported in this manuscript reveals that covalent adduction can also occur at C173, located within a conformationally flexible loop. Notably, a recent study has also identified C173 as a site for electrophilic engagement, suggesting this loop may represent a broader hotspot for DCAF16-targeted covalent degrader design (24).

Here, we also demonstrate that small changes in the degradation tail can influence E3 ligase preference. Specifically, increasing the linker length by one carbon between the alkyne and azepane (**3**) dialled out contributions from DCAF16, thus resulting in a degrader that relies mostly on FBXO22. Complementing the chemical effort to fine-tune ligase recruitment, we also identify a DCAF16 gain-of-function mutant, L59W. This mutant can synthetically dial in dependency on DCAF16 for **2** and **3**, which otherwise leverage FBXO22 as the driver ligase, by locally fine-tuning key interactions for degrader and neo-substrate recruitment. Collectively, these findings outline how chemical and genetic adaptation of complementary surfaces between E3 ligases and target proteins can inform the design of tuneable degraders.

The uncovered dual ligase mechanism has relevant implications as it may enhance the therapeutic efficacy of degraders and potentially overcome resistance mechanisms that arise via E3 ligase mutations or downregulation in cancer cells (46). We have previously shown how resistance mutations concentrate in substrate receptors CRBN and VHL, particularly in regions crucial for assembling the ternary complex with the degrader and target (29). Given that loss of E3 ligase function is a key resistance mechanism, we previously developed heterotrivalent PROTACs, combining VHL, CRBN, and target ligands (33). The compounds described in this study present an alternative strategy to overcome resistance through monovalent and compact degrader designs. Their ability to chemically toggle between distinct E3 ligase complexes adds a valuable layer of flexibility, enabling modulation of ligase engagement to help bypass or delay resistance mechanisms. As novel chemical modalities continue to emerge, such approaches will be essential for advancing next-generation targeted therapies with greater adaptability and resistance resilience.

## Acknowledgements

The Winter lab is supported by funding from the European Research Council (ERC) under the European Union’s Horizon 2020 research and innovation program (grant agreement 851478), as well as by funding from the Austrian Science Fund (FWF, projects P7909, P36746 and P5918723) and the Vienna Science and Technology Fund (WWTF, project LS21-015). D.S. was supported by EMBO ALTF 283-2023 fellowship. Work in the Ciulli lab on this project was supported by Eisai Co., Ltd. and by the pharmaceutical companies supporting the Division of Signal Transduction and Therapy (Boehringer Ingelheim, GlaxoSmithKline, Merck KaaG) as sponsored research funding to A.C. Funding in the Ciulli Lab is also gratefully acknowledged from the Innovative Medicines Initiative 2 (IMI2) Joint Undertaking under grant agreement no. 875510 (EUbOPEN project). The IMI2 Joint Undertaking receives support from the European Union’s Horizon 2020 research and innovation programme, European Federation of Pharmaceutical Industries and Associations (EFPIA) companies, and associated partners KTH, OICR, Diamond and McGill. A.C.S. was supported by a ‘Fundación Alfonso Martín Escudero’ (Madrid, Spain) postdoctoral fellowship and a UKRI Postdoctoral Fellowships Guarantee Scheme (Council ref. *EP/Z002176/1*) funding the Marie Skłodowska–Curie Actions Individual Fellowship (HORIZON-MSCA-2023-PF-01 101152759 DEGRON). A.D.C. was supported by a Horizion2020 Marie Skłodowska–Curie Actions Individual Fellowship (H2020-MSCA-IF-2020-101024945 DELETER).

We thank the Core Facility for Flow Cytometry of the Medical University of Vienna for access to flow cytometry instruments and assistance with cell sorting and the CeMM Biomedical Sequencing Facility for NGS sample processing, sequencing and data curation. We thank Nathanael Gray for the gift of MMH2. Maria Rodriguez-Rios for the gift of BRD4 BD2 recombinant protein, as well as Maria Rodriguez-Rios and Roberta Ibba for native MS help. We acknowledge the University of Dundee Cryo-EM facility for access to the instrumentation, funded by the Wellcome Trust (223816/Z/21/Z) and MRC (MRC World Class Laboratories PO 4050845509). Finally, Paula da Fonseca,Edward P. Morris, and Kyle Dent for helpful discussion and Diamond for access and support of the cryo-EM facilities at the UK national electron Bio-Imaging Centre (eBIC), proposal BI37630.

## Author contributions

V.A.S., D.S., A.C.S., A.C. and G.E.W. conceived and planned this project. V.A.S., D.S. and A.C.S. designed and conducted experiments with help from K.I., R.C., M.M.O., M.A.N., and G.S. V.A.S., D.S., A.C.S., A.C. and G.E.W. analysed and interpreted original data. K.I. and R.C. designed and synthesized compounds. D.S. and C.S. performed and analysed FACS-based CRISPR screens. D.S. and M.M.O. performed and analysed DMS experiments. A.C.S. and G.S. performed and analysed quantitative expression proteomics and BioID. V.A.S. performed cryo-EM imaging, data processing and 3D reconstruction with help from M.A.N. V.A.S performed and analysed Intact MS and TR-FRET experiments. V.A.S, M.A.N., A.C.S., L.K., H.E.P., M.D., A.D.C and A.T. established critical reagents and methodology. V.A.S., D.S., A.C.S., A.C. and G.E.W. co-wrote the manuscript with input from all co-authors.

## Competing interests

A.C. is a scientific founder and shareholder of Amphista Therapeutics, a company that is developing targeted protein degradation therapeutic platforms. A.C. is on the Scientific Advisory Board of ProtOS Bio. The Ciulli laboratory receives or has received sponsored research support from Almirall, Amgen, Amphista Therapeutics, Boehringer Ingelheim, Eisai, Merck KaaG, Nurix Therapeutics, Ono Pharmaceutical and Tocris-Biotechne. G.E.W. is scientific founder and shareholder of Proxygen and Solgate. The Winter lab received research funding from Pfizer. G.E.W. is a scientific founder and shareholder of Proxygen and Solgate and on the scientific advisory board of Proxygen. He holds equity in Cellgate Therapeutics. He also holds equity in Nexo Therapeutics and serves on their scientific advisory board. The Winter lab has received research funding from Pfizer

## Data availability

Source data for Figure 1b, d Figure 2a, Figure 3 Figure 4b,c, Fig 5a Extended Data Figure 2c, Extended Data Figure 3a,b, Extended Data Figure 8e, and Extended Data Figure 10a, b are included in the supplementary information files of the manuscript (Supplementary Tables 1-4 and Extended data table 1 and 2). Cryo-EM density maps are deposited in the EMDB with the accession code EMD-54549. The atomic model is deposited under Protein Data Bank ID 9S3R, extended PDB ID pdb_00009S3R. Atomic coordinates will be made available upon publication. Quantitative proteomics data have been deposited to the ProteomeXchange Consortium PRIDE repository with the accession codes PXD065445 and PXD065523. All biological materials are available upon reasonable requests under material transfer agreements (MTA) with The Centre for Targeted Protein Degradation, University of Dundee, or CeMM Research Center for Molecular Medicine of the Austrian Academy of Sciences, respectively.

## Extended Data Figures

**Extended Data Figure 1.**
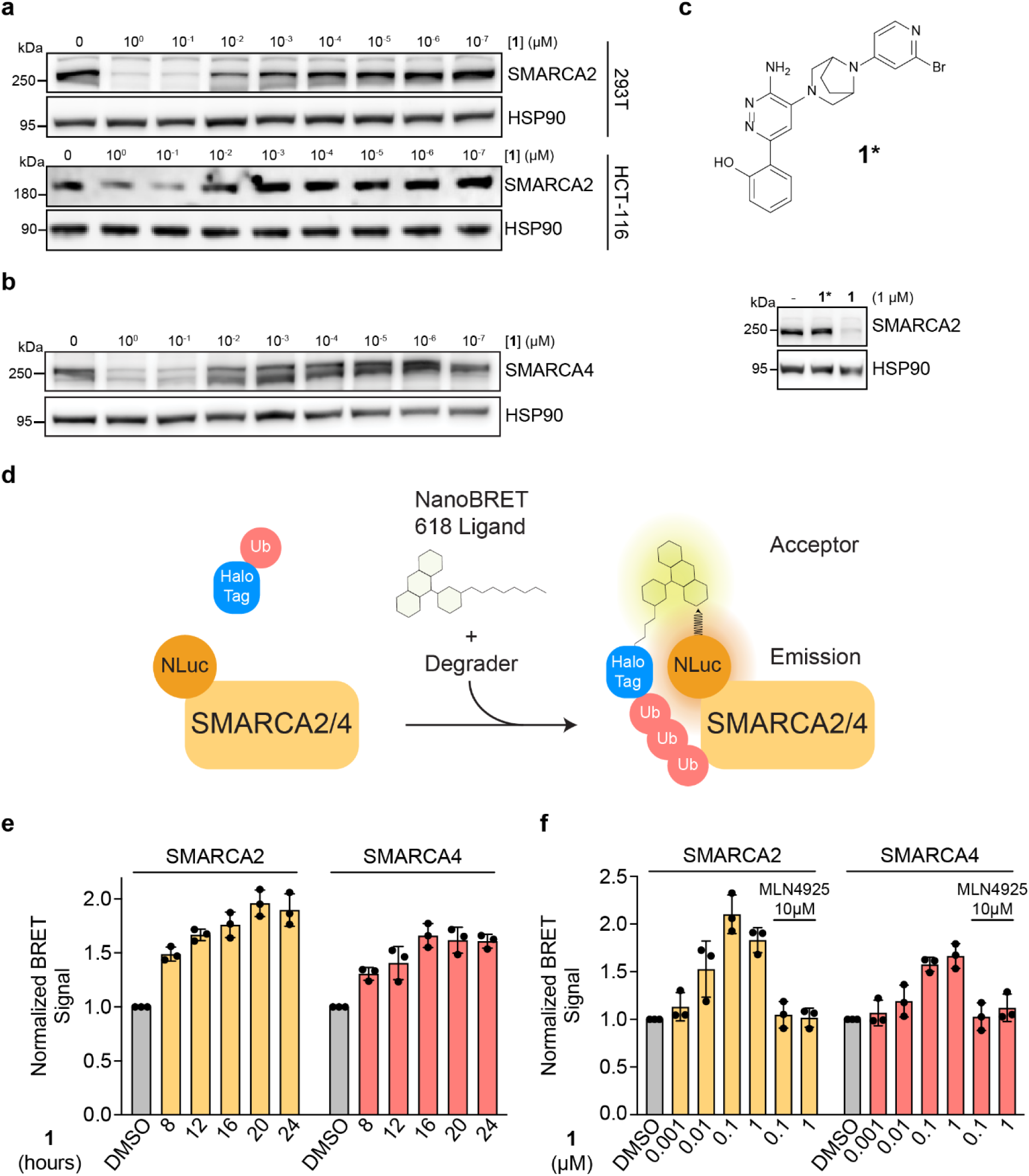
Compound 1 induces ubiquitination and degradation of SMARCA2/4. **a**, **b**, Compound **1** induced degradation of SMARCA2 (**a**) and SMARCA4 (**b**). HEK293T and HCT116 cells were treated with DMSO or a serial dilution series of **1** for 24 h. SMARCA2 and SMARCA4 were quantified by immunoblot. **c**, Structure of inactive control compound **1\*** and immunoblot quantification of SMARCA2 in HEK293T cells after treatment with DMSO, **1\***, or **1** (1 µM) for 24 hours. d, Schematic of NanoBRET ubiquitination assay to evaluate cellular ubiquitination of SMARCA2/4 in the presence of a degrader. **e**, Kinetic NanoBRET ubiquitination assay. HEK293T SMARCA2/4-HiBiT knock-in cells transfected with LgBiT and HaloTag-Ubiquitin were treated with DMSO or **1** (0.1 µM) for various time points. Data represent mean ± SD, n = 3 biological replicates. **f**, Concentration-based NanoBRET ubiquitination assay. HEK293T SMARCA2/4-HiBiT knock-in cells transfected with LgBiT and HaloTag-Ubiquitin were treated with DMSO or increasing concentrations of **1** (+/- 1 hour pre-treatment with 10 µM MLN4924) for 24 hours. Data represent mean ± SD, n = 3 biological replicates.

**Extended Data Figure 2.**
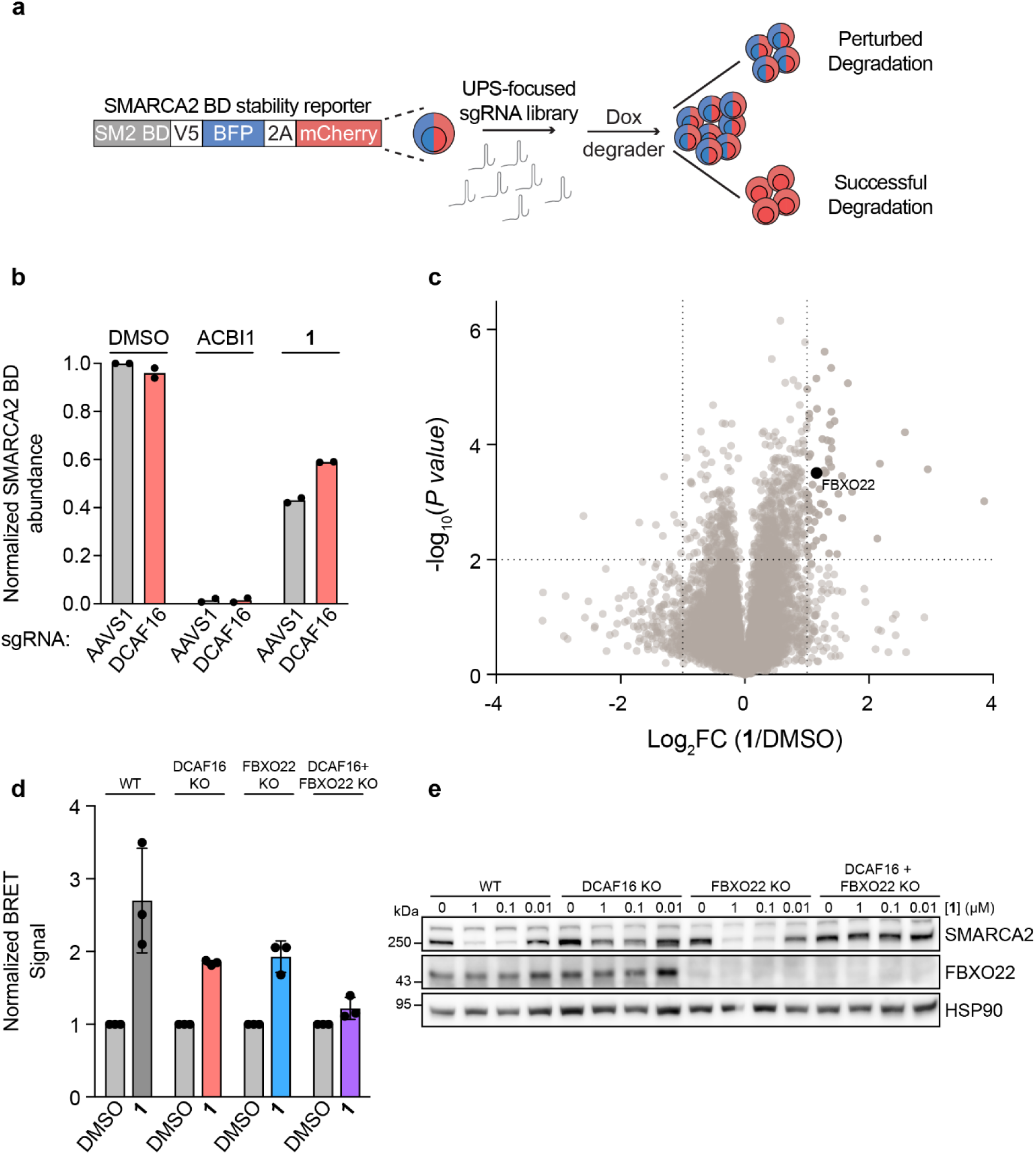
Compound 1 degrades SMARCA2/4 via DCAF16 and FBXO22. **a**, Schematic for FACS-based CRISPR-Cas9 screen with a SMARCA2 stability reporter and **1 b**, CRISPR screen validation. KBM7 iCas9 cells expressing the SMARCA2 BD stability reporter and transduced with AAVS1 or DCAF16-targeting sgRNAs were treated with DMSO, ACBI1 (1 µM), or **1** (1 µM) for 24 hours. SMARCA2-BFP and mCherry levels were quantified by flow cytometry. Data represent mean ± SD, n = 2 biological replicates. **c**, BioID analysis with **1** treatment. Proximity labelling in Flp-In T-REx HEK293 miniTurboID-SMARCA2-BD inducible cell line. Cells were treated with 1 µg/ml doxycycline for a total of 20.5 hours. 12 hours after induction, cells were pre-treated with 10 µM MLN4924 for 30 mins, followed by addition of **1** (10 µM). Volcano plot of BioID results showing log_2_ fold change (FC) and −log_10_ adjusted P values derived from normalized protein spectral counts and compared with DMSO treatment. Scoring window (P value < 0.01, log2 FC > 1) is indicated, n = 3 biological replicates. **d**, NanoBRET ubiquitination assay in KO cell lines. HEK293T SMARCA4-HiBiT knock-in cells with single and double E3 ligase KOs transfected with LgBiT and HaloTag-Ubiquitin were treated with DMSO or **1** (1 µM) for 24 hours. Data represent mean ± SD, n = 3 biological replicates. **e**, Genetic rescue of SMARCA2 degradation. WT, DCAF16 (KO), FBXO22 KO, and double KO HEK293T cells were treated with DMSO or increasing concentrations of **1** for 24 hours. SMARCA2 was quantified by immunoblot.

**Extended Data Figure 3.**
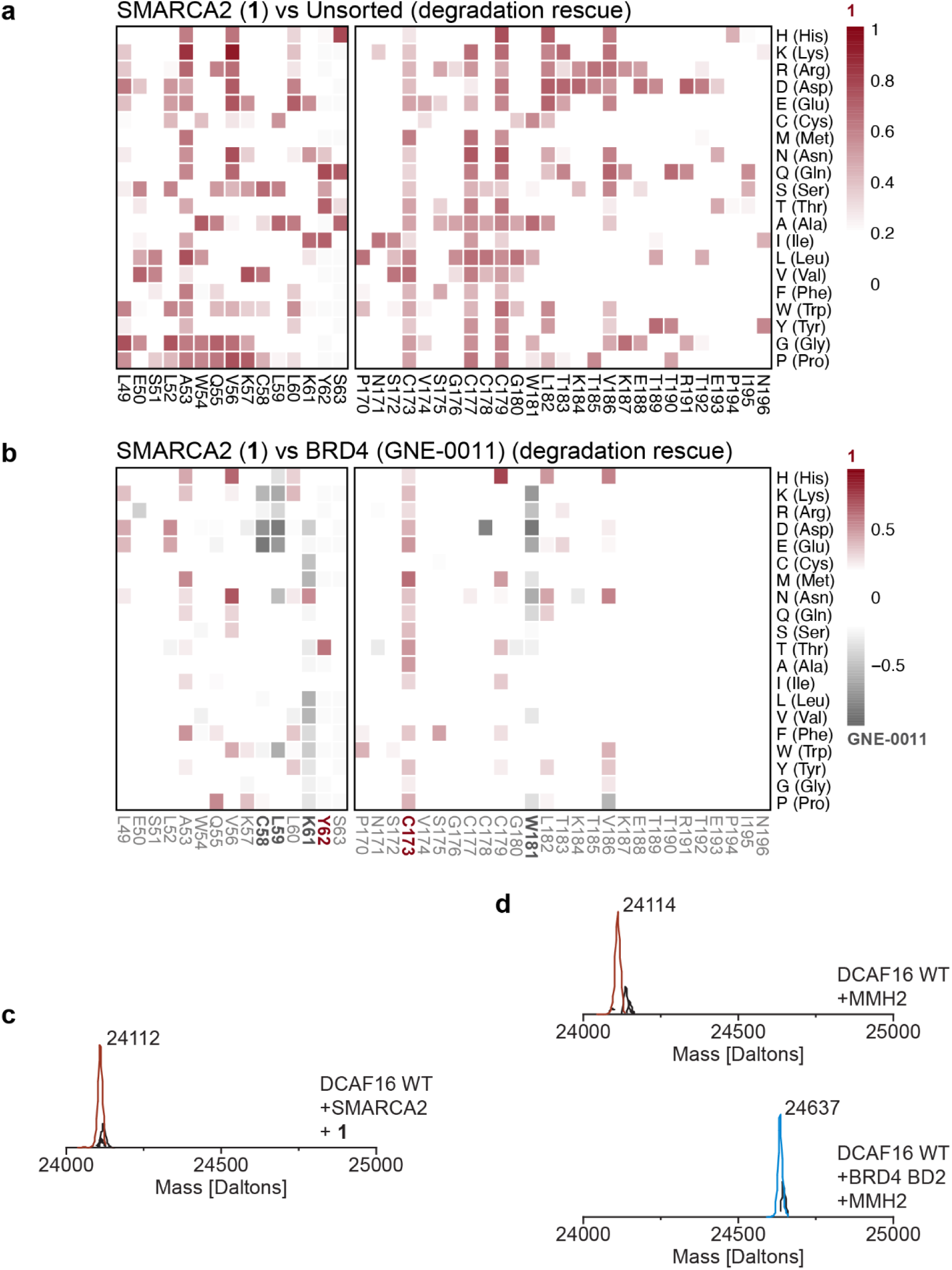
Compound 1 does not engage with DCAF16 in a template-assisted manner. **a**, **b**, DMS of DCAF16 for degradation rescue. DCAF16 and FBXO22 Double knockout HEK293T cells stably expressing stability reporters for SMARCA2 BD or BRD4 tandem-BDs were transduced with a library of EGFP tagged DCAF16 variants containing single amino acid substitutions at every position. SMARCA2 and BRD4 stability reporter cells were treated with **1** (1 µM) and GNE-0011 (0.1 µM) for 24 hours, respectively. Heatmap depicting mean log_2_ fold enrichment of DCAF16 mutations normalized to maximum log_2_ fold changes for **1** versus unsorted control (**a**). Heatmap depicting differential log_2_ fold enrichment of DCAF16 mutations normalized to maximum log_2_ fold changes versus unsorted control between SMARCA2 targeting **1** and BRD4 targeting GNE-0011 (**b**). Data corresponds to SMARCA2^High^ and BRD4^High^ sorted populations, showing mutations that specifically prevent SMARCA2 degradation (red) and BRD4 degradation (gray), n=3 independent measurements. c, Intact protein mass spectra for WT DCAF16 (10 µM) and **1** (20 µM) in the presence of SMARCA2 BD (20 µM). **d**, Intact protein mass spectra for WT DCAF16 (10 µM) and MMH2 (20 µM) in the absence (top) or presence (bottom) of BRD4 BD2 (20 µM)

**Extended Data Figure 4.**
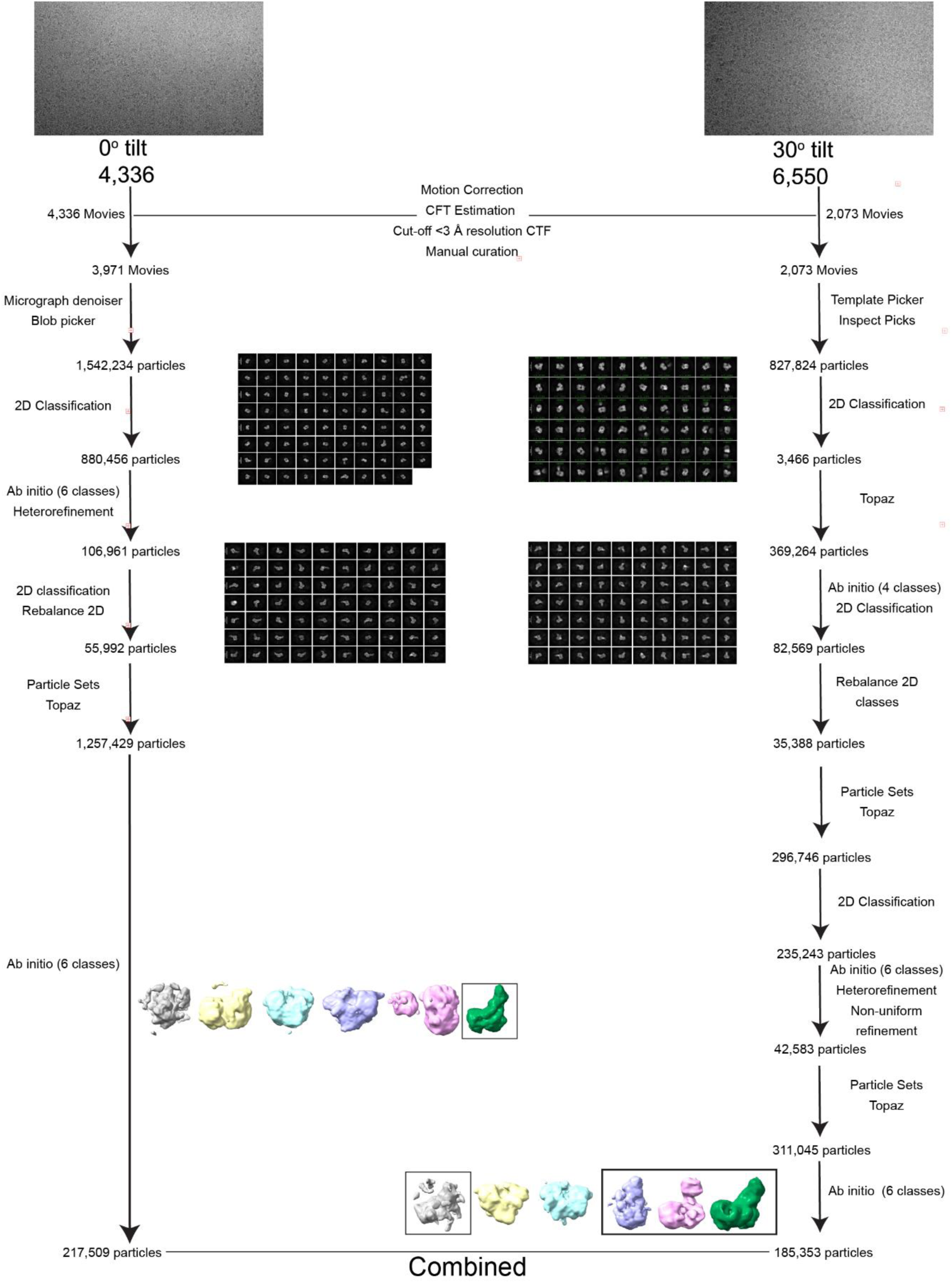
cryo-EM data processing Part 1. Workflow for Cryo-EM data processing for 0° and 30° datasets.

**Extended Data Figure 5.**
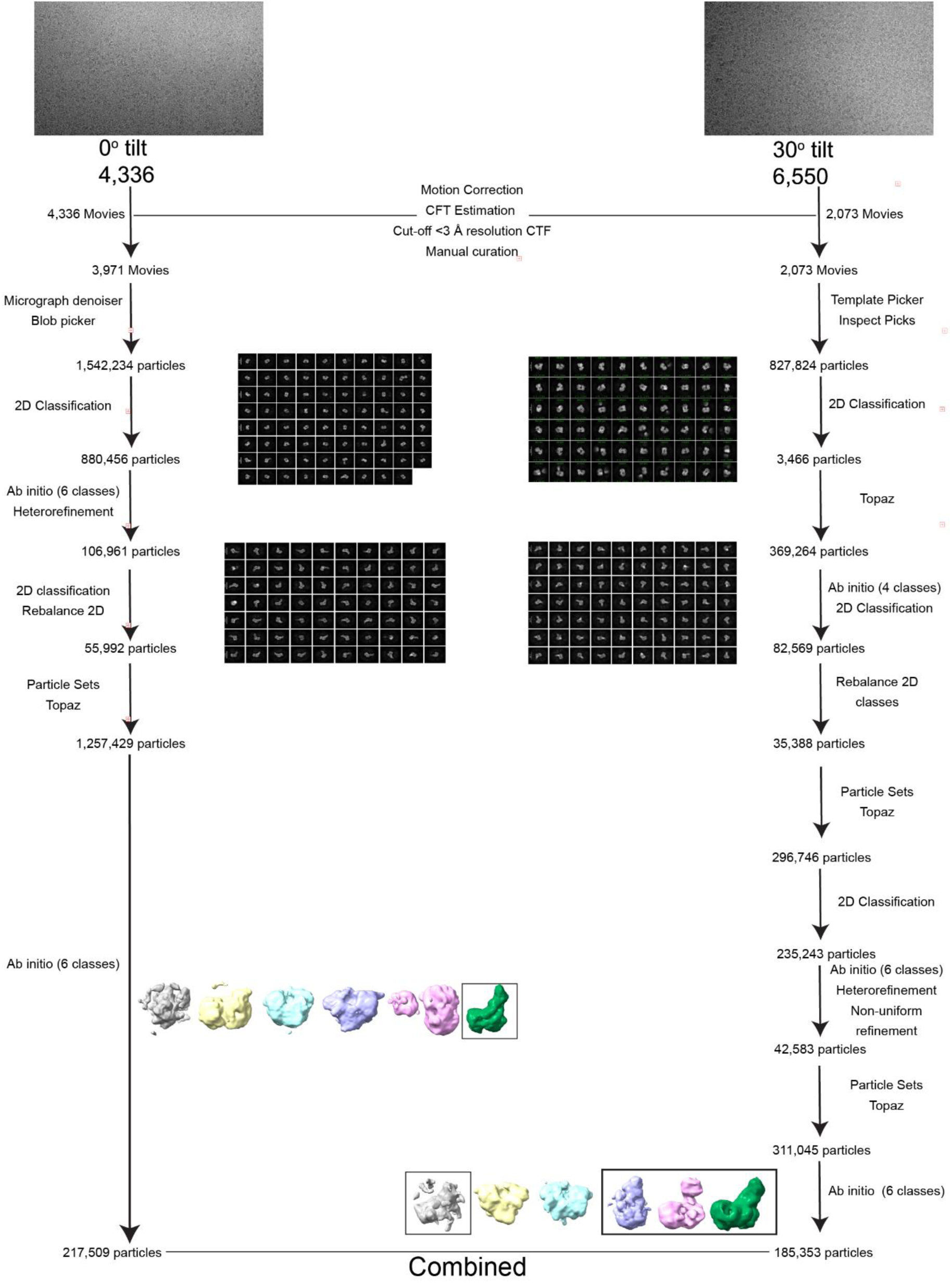
cryo-EM data processing Part 2. **a**, Workflow for Cryo-EM data processing of combined 0° and 30° datasets. **b**, **c**, Angular distribution plot (**b**) and posterior position directional distribution plot (**c**) for the final local refinement. **d**, Gold-standard Fourier shell correlation at a cut-off of 0.143. **e**, Local resolution estimation.

**Extended Data Figure 6.**
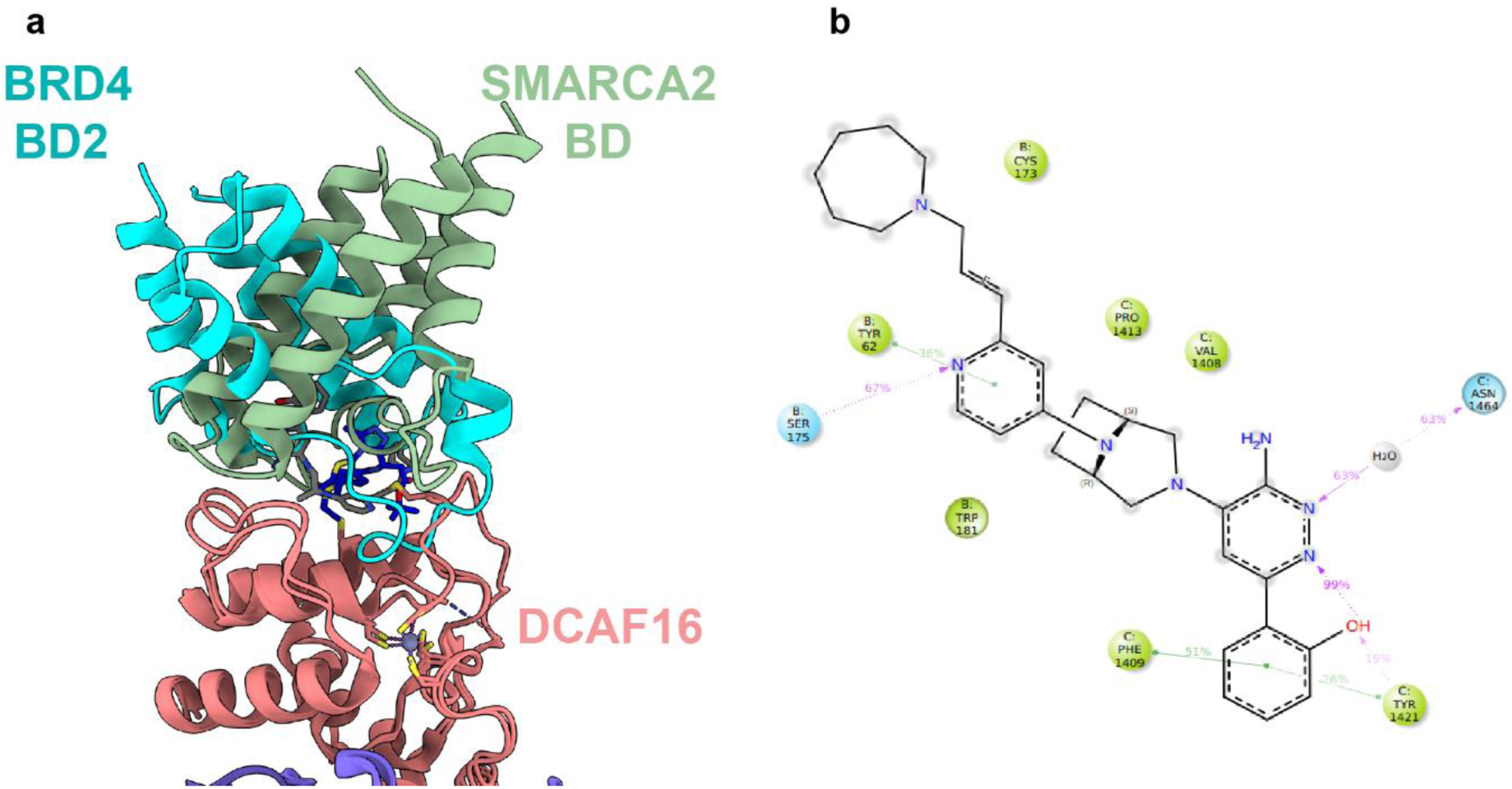
Compound 1 glues SMARCA2 to DCAF16 in a structurally distinct manner relative to BRD4. **a**, Super-imposed binding mode of BRD4 BD2 (PDB ID: 8G46) and SMARCA2 BD on DCAF16. **b**, Protein-ligand interactions map from molecular dynamics simulation of DCAF16:DDB1 (ΔBPB): DDA1, SMARCA2, **1** ternary complex.

**Extended Data Figure 7.**
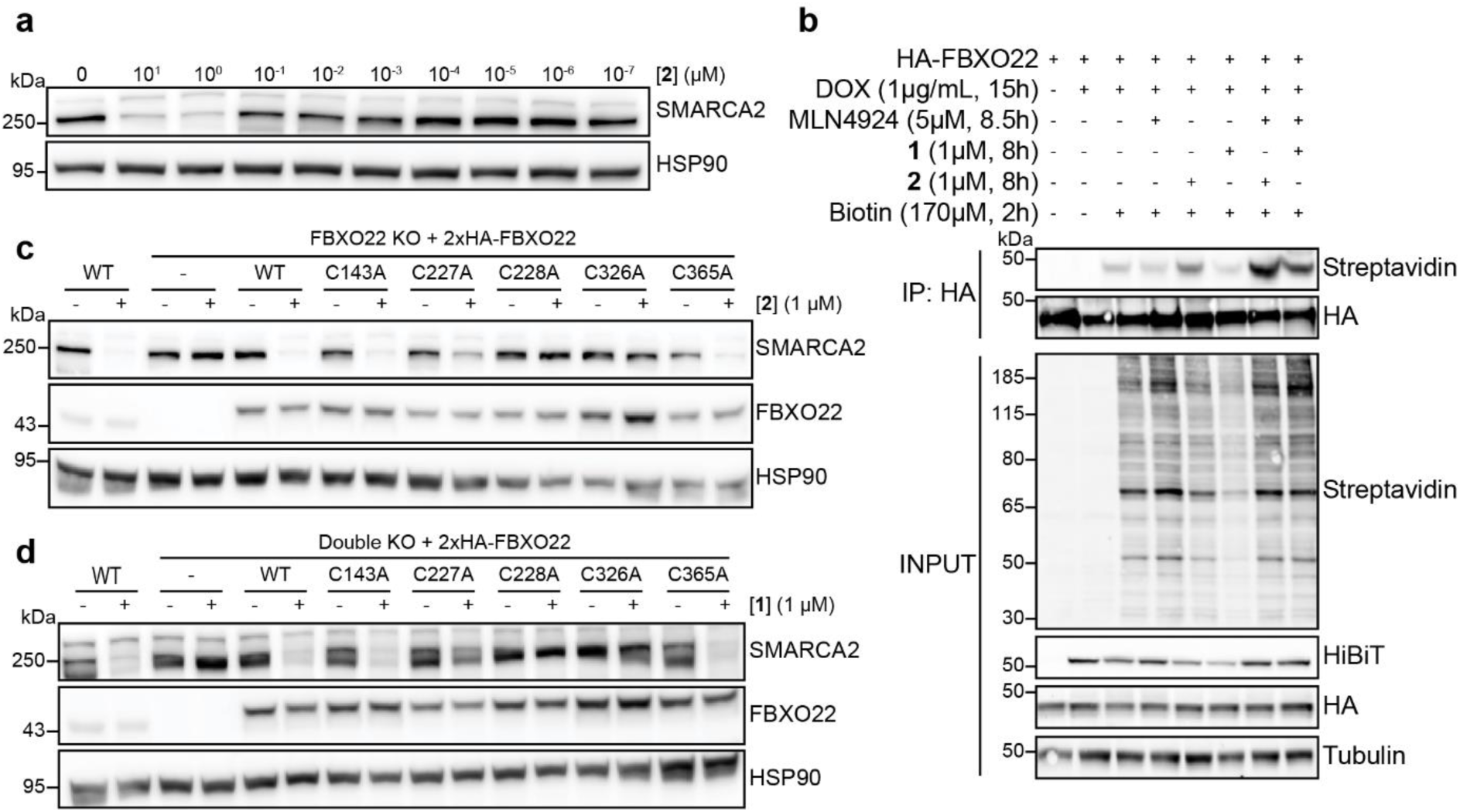
Compound 2 engages with FBXO22 via C228 and C326. **a**, Degradation of SMARCA2 by 2. HEK293T cells were treated with DMSO or a serial dilution series of **2** for 24 h. SMARCA2 was quantified by immunoblot. **b**, Biotin proximity labelling of FBXO22. Flp-In T-REx HEK293 cells transiently expressing HA-tagged FBXO22 and doxycycline-inducible miniTurboID-SMARCA2-BD were pretreated with 5 µM MLN4924 for 30 min, followed by **1** (1 µM) or **2** (1 µM) and 170 µM biotin for 8 and 2 hours, respectively. HA-tagged FBXO22 was immunoprecipitated and biotinylated proteins were detected via streptavidin immunoblotting. **c**, **d**, Immunoblots with FBXO22 Cysteine mutants. FBXO22 KO HEK293T cells stably expressing HA tagged FBXO22 variants were treated with **2** (1 µM) for 24 hours (**c**). Double KO HEK293T cells stably expressing HA tagged FBXO22 variants were treated with **1** (1 µM) for 24 hours (**d**). SMARCA2 was quantified by immunoblot.

**Extended Data Figure 8.**
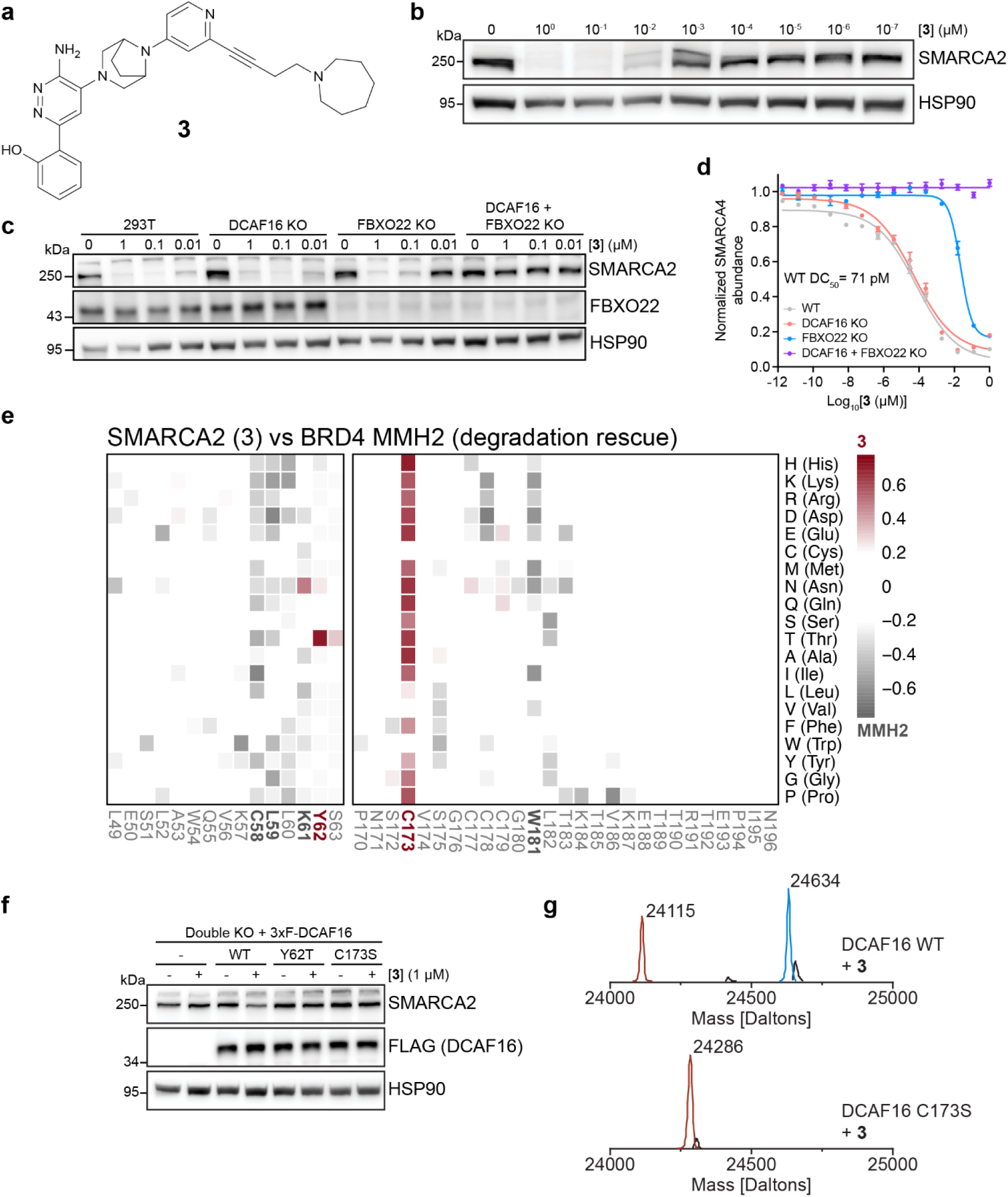
Compounds 3 also exhibits a driver ligase-switch and covalently interacts with DCAF16 via C173. **a**, structure of Compound **3**. **b**, Degradation of SMARCA2 by **3**. HEK293T cells were treated with DMSO or a serial dilution series of **3** for 24 h. SMARCA2 was quantified by immunoblot. **c**, Genetic rescue of SMARCA2 degradation. WT, DCAF16 KO, FBXO22 KO, and double KO HEK293T cells were treated with DMSO or increasing concentrations of **3** for 24 hours. SMARCA2 was quantified by immunoblot. **d**, HiBiT endpoint degradation assay. HEK293T SMARCA4-HiBit cells with single and double KOs were treated with a serial dilution series of **3** for 16 hours. SMARCA4 levels were quantified using the HiBiT lytic detection system. Data represent mean ± SEM, n = 6 biological replicates. **e**, Deep mutagenesis scanning (DMS) of DCAF16 for degradation rescue. DCAF16 and FBXO22 Double KO HEK293T cells stably expressing stability reporters for SMARCA2 BD or BRD4 tandem-BDs were transduced with a library of EGFP tagged DCAF16 variants containing single amino acid substitutions at every position. SMARCA2 and BRD4 stability reporter cells were treated with **3** (1 µM) and MMH2 (1 µM) for 24 hours, respectively. Heatmap depicting differential log_2_ fold enrichment of DCAF16 mutations normalized to maximum log_2_ fold changes versus unsorted control between SMARCA2 targeting **3** and BRD4 targeting MMH2. Data corresponds to SMARCA2^High^ and BRD4^High^ sorted populations, showing mutations that specifically prevent SMARCA2 degradation (red) and BRD4 degradation (gray) n=3 independent measurements. **f**, Validation of deep mutagenesis scanning. Double knockout HEK293T cells stably expressing 3xFlag tagged DCAF16 variants were treated with **3** (1 µM) for 24 hours. SMARCA2 was quantified by immunoblot. **g**, Intact protein mass spectra for WT (top) or C173S mutant (bottom) DCAF16 (10 µM) and **3** (20 µM).

**Extended Data Figure 9.**
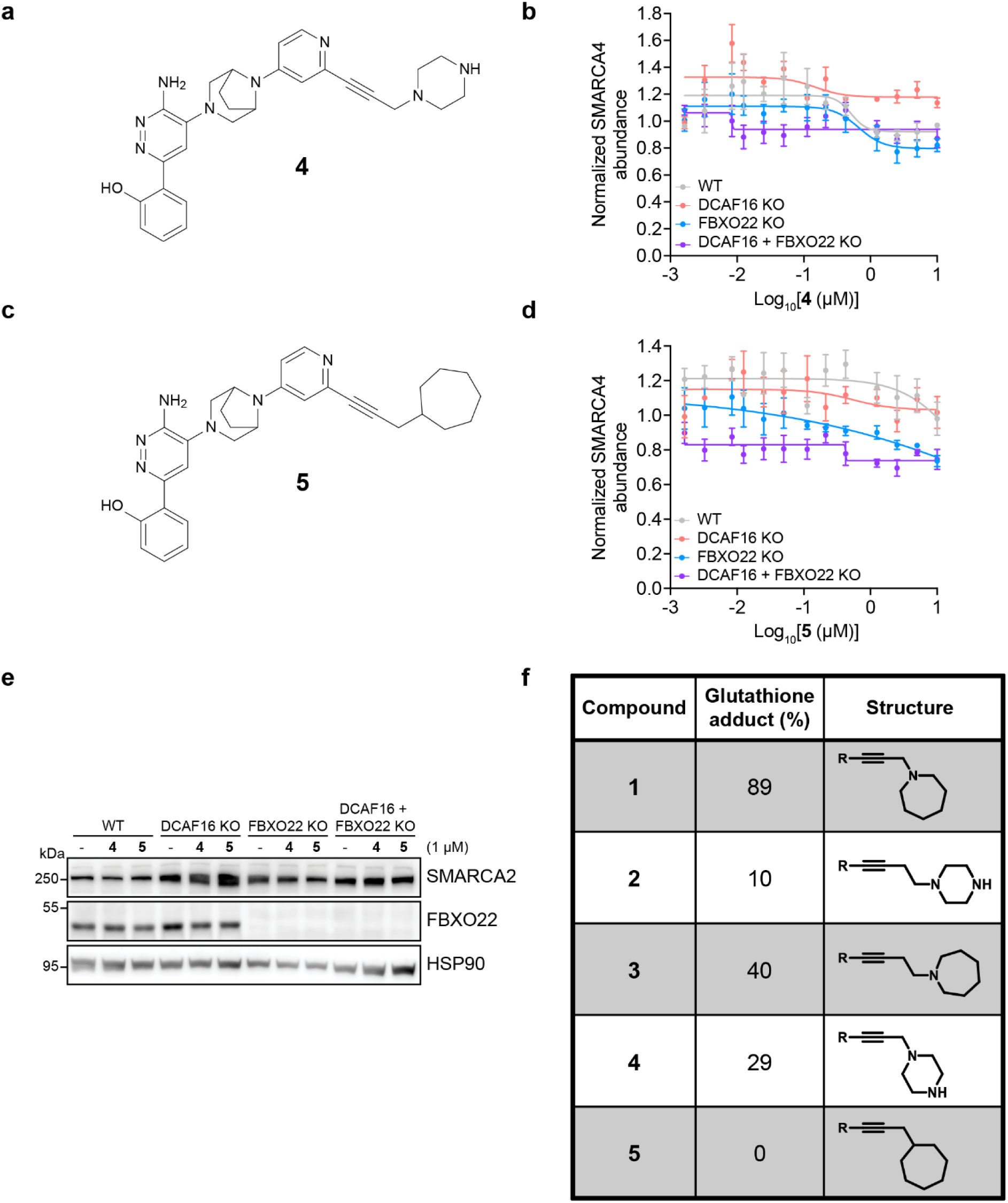
Compounds 4 and 5 do not degrade SMARCA2/4. **a**, Structure of Compound **4**. **b**, HiBiT endpoint degradation assay. SMARCA4-HiBiT HEK293T cells with single and double KOs were treated with a serial dilution series of **4** for 24 hours. SMARCA4 levels were quantified using the HiBiT lytic detection system. Data represent mean ± SEM, n = 3 biological replicates. **c**, Structure of Compound **5**. **d**, Hibit endpoint degradation assay. SMARCA4-HiBit HEK293T cells with single and double KOs were treated with a serial dilution series of **5** for 24 hours. SMARCA4 levels were quantified using the HiBiT lytic detection system. Data represent mean ± SEM, n = 3 biological replicates. **e**, Genetic rescue of SMARCA2 degradation. WT, DCAF16 KO, FBXO22 KO, and double KO HEK293T cells were treated with DMSO or increasing concentrations of **4** or **5** for 24 hours. SMARCA2 was quantified by immunoblot. **f**, Glutathione reactivity assay. The relative reactivity of SMARCA2/4 degraders was assessed using a 3-hour incubation with glutathione at 37°C. Reactivity was estimated as the UV area ratio of the glutathione adduct to the starting material (%), without external calibration

**Extended Data Figure 10.**
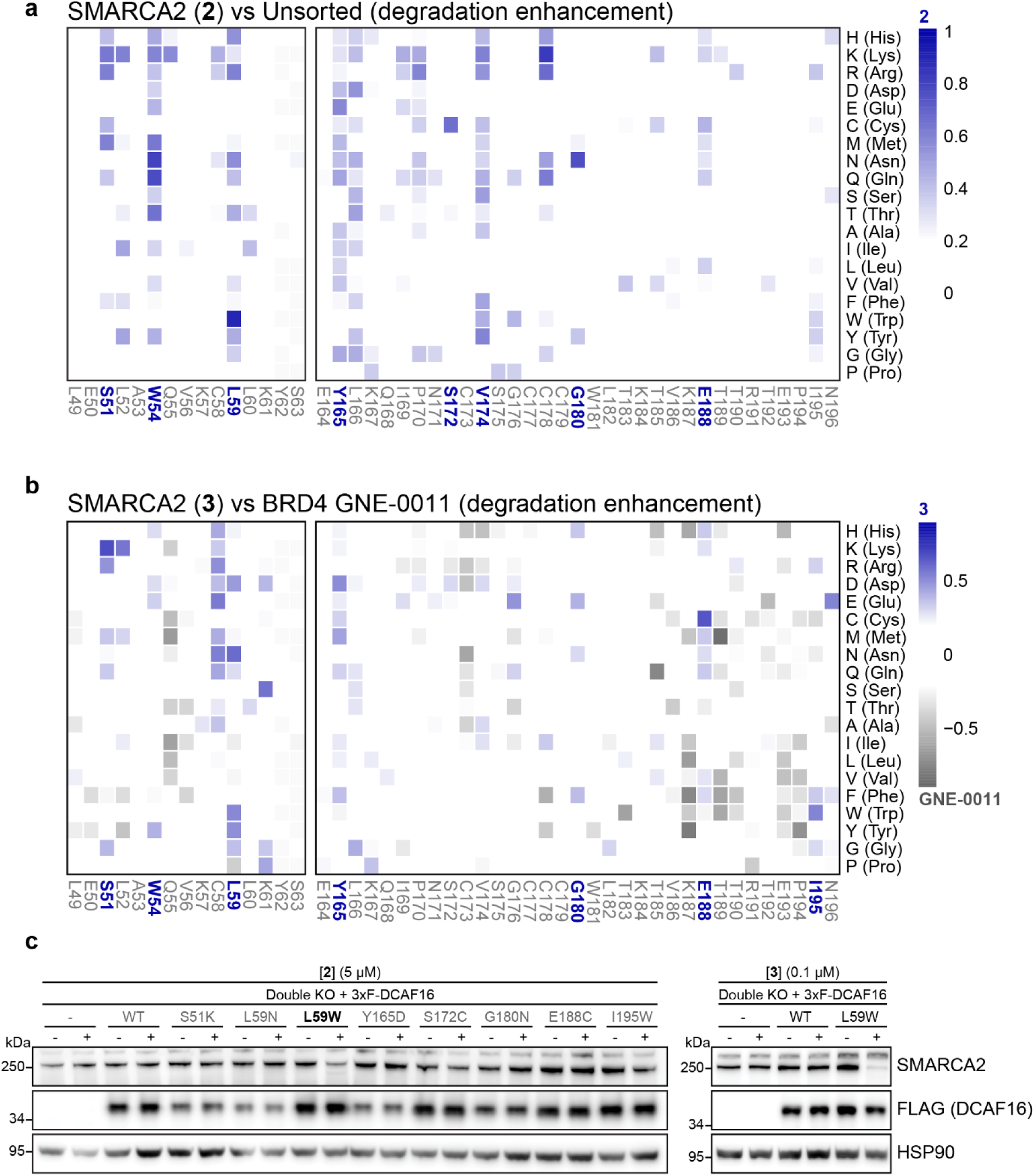
Mutations in DCAF16 that enhance degradation of SMARCA2 via compounds 2 and 3. **a**, **b**, Deep mutagenesis scanning (DMS) of DCAF16 for degradation enhancement. DCAF16 and FBXO22 Double knockout HEK293T cells stably expressing stability reporters for SMARCA2 BD or BRD4 tandem-BDs were transduced with a library of EGFP tagged DCAF16 variants containing single amino acid substitutions at every position. SMARCA2 stability reporter cells were treated with **2** (1 µM) or **3** (1µM), and BRD4 stability reporter cells were treated with GNE-0011 (0.1 µM) for 24 hours. Heatmap depicting mean log_2_ fold enrichment of DCAF16 mutations normalized to maximum log_2_ fold changes for **2** versus unsorted control (**a**). Heatmap depicting differential log_2_ fold enrichment of DCAF16 mutations normalized to maximum log_2_ fold changes versus unsorted control between SMARCA2 targeting compound **3** and BRD4 targeting GNE-0011 (**b**). Data corresponds to SMARCA2^Low^ and BRD4^Low^ sorted populations, showing mutations that specifically enhance SMARCA2 degradation (blue) and BRD4 degradation (gray), n=3 independent measurements. **c**, Validation of DMS. Double KO HEK293T cells stably expressing 3xFlag tagged DCAF16 variants were treated with **2** (5 µM) (left) or **3** (0.1 µM) (right) for 24 hours. SMARCA2 was quantified by immunoblot. Left and right represent separate Western blots.

**Extended Data Table 1:**
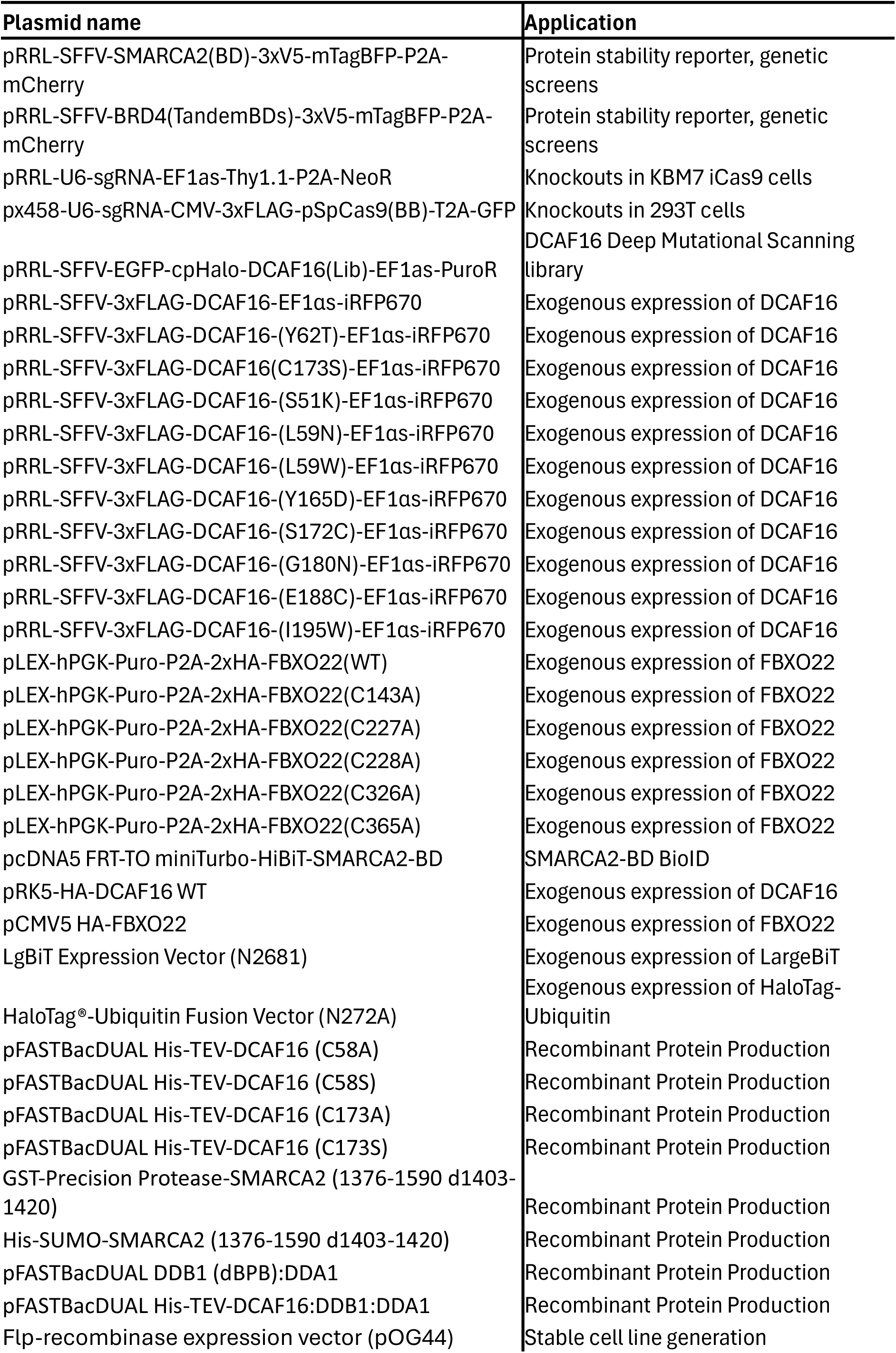
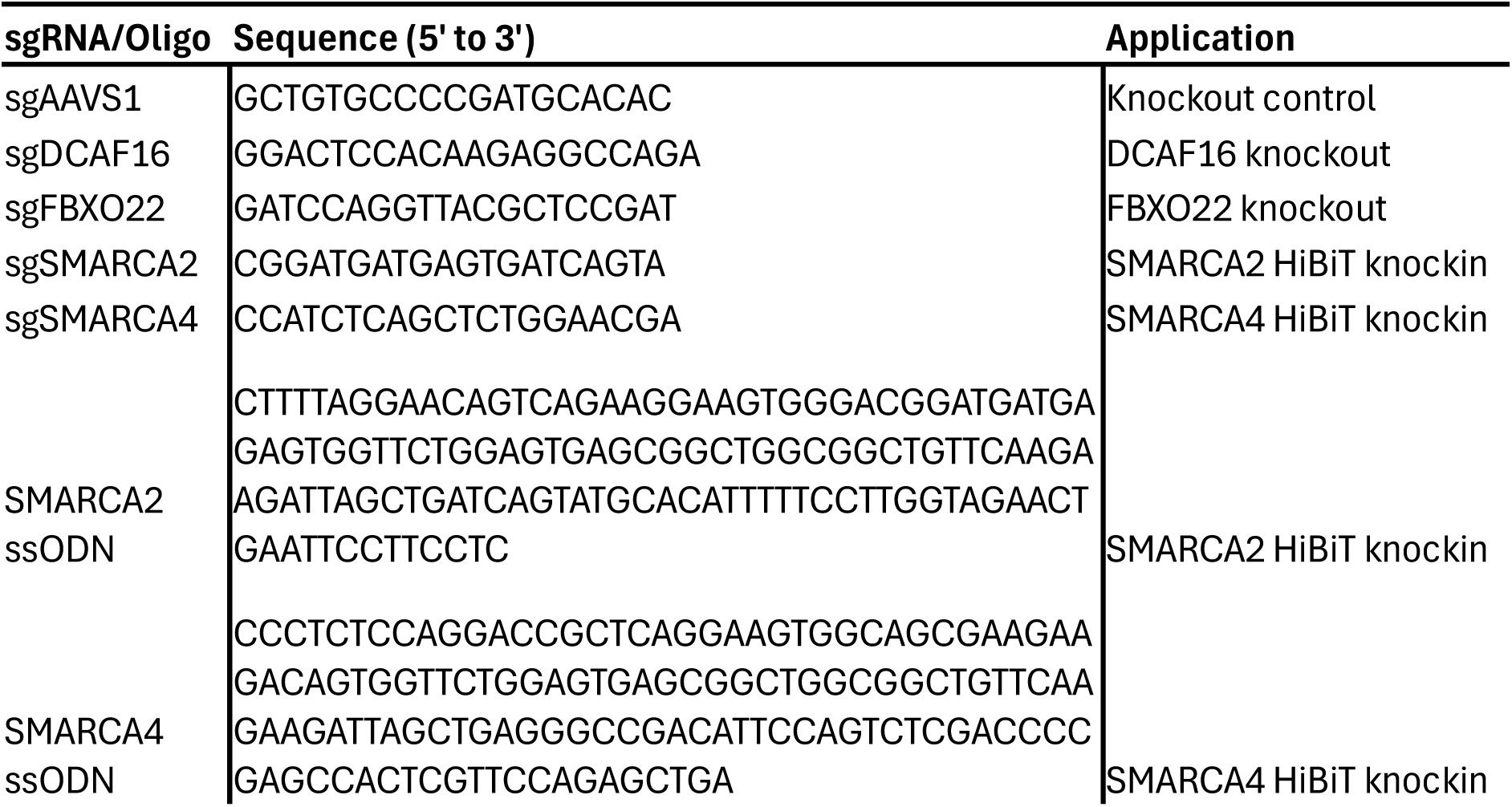
Plasmids and sgRNAs used in this study.

**Extended Data Table 2:**
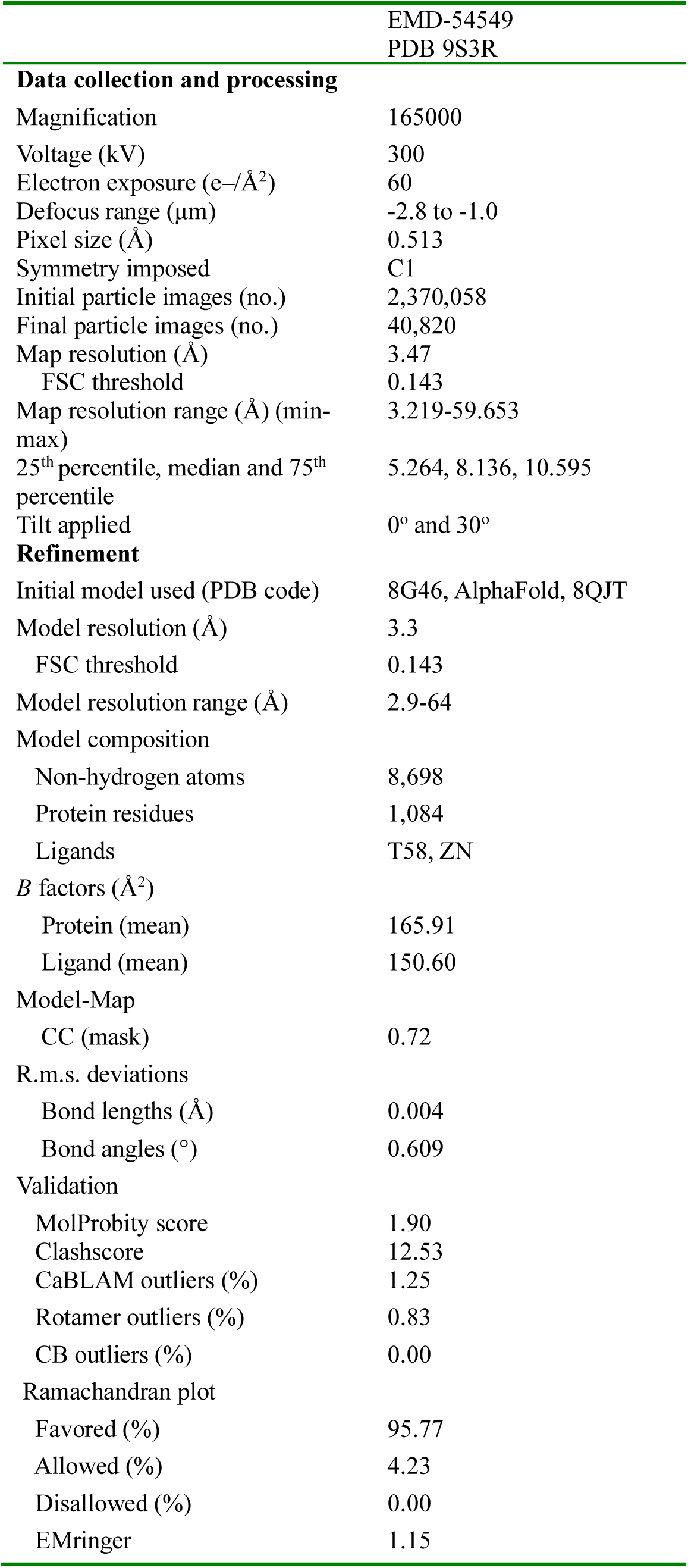
Summary of cryo-EM data collection conditions and image processing.

|  |  |
| --- | --- |
|  | EMD-54549 |
|  | PDB 9S3R |
| **Data collection and processing** |  |
| Magnification | 165000 |
| Voltage (kV) | 300 |
| Electron exposure (e-/Å<sup>2</sup>) | 60 |
| Defocus range (μm) | -2.8 to -1.0 |
| Pixel size (Å) | 0.513 |
| Symmetry imposed | C1 |
| Initial particle images (no.) | 2,370,058 |
| Final particle images (no.) | 40,820 |
| Map resolution (Å) | 3.47 |
| FSC threshold | 0.143 |
| Map resolution range (Å) (min-max) | 3.219-59.653 |
| 25<sup>th</sup> percentile, median and 75<sup>th</sup> percentile | 5.264, 8.136, 10.595 |
| Tilt applied | 0° and 30° |
| **Refinement** |  |
| Initial model used (PDB code) | 8G46, AlphaFold, 8QJT |
| Model resolution (Å) | 3.3 |
| FSC threshold | 0.143 |
| Model resolution range (Å) | 2.9-64 |
| Model composition |  |
| Non-hydrogen atoms | 8,698 |
| Protein residues | 1,084 |
| Ligands | T58, ZN |
| *B* factors (Å<sup>2</sup>) |  |
| Protein (mean) | 165.91 |
| Ligand (mean) | 150.60 |
| Model-Map |  |
| CC (mask) | 0.72 |
| R.m.s. deviations |  |
| Bond lengths (Å) | 0.004 |
| Bond angles (°) | 0.609 |
| Validation |  |
| MolProbity score | 1.90 |
| Clashscore | 12.53 |
| CaBLAM outliers (%) | 1.25 |
| Rotamer outliers (%) | 0.83 |
| CB outliers (%) | 0.00 |
| Ramachandran plot |  |
| Favored (%) | 95.77 |
| Allowed (%) | 4.23 |
| Disallowed (%) | 0.00 |
| EMringer | 1.15 |

## Experimental Methods

### Chemical Synthesis

Chemicals that are commercially available were purchased from Apollo Scientific, Sigma-Aldrich, Fluorochem, CombiBlocks, TCI, and Enamine and were used without further purification. Liquid chromatography–mass spectrometry (LC-MS) was carried out on a Shimadzu HPLC/MS 2020 equipped with a Hypersil Gold column (1.9 μm, 50 × 2.1 mm2), a photodiode array detector, and an electrospray ionization (ESI) detector. The samples were eluted with a 3 min gradient of 5–95% acetonitrile in water containing 0.1% formic acid at a flow rate of 0.8 mL/min. Flash column chromatography was performed on a Teledyne ISCO Combiflash Companion installed with disposable normal phase RediSep Rf columns (230–400 mesh, 40–63 mm; SiliCycle). Preparative HPLC purification was performed on a Gilson preparative HPLC system equipped with a Waters X-Select C18 column (100 mm × 19 mm and 5 μm particle size) using a gradient from 5 to 95% of acetonitrile in water containing 0.1% formic acid over 10 min at a flow rate of 25 mL/min. Compound characterization using NMR was performed either on a Bruker 500 Ultra shield or on a Bruker Ascend 400 spectrometer. The 1H NMR and 13C NMR reference solvents used are CDCl3 (δH = 7.26 ppm/δC = 77.16 ppm), CD_3_OD (δH = 3.31 ppm/δC = 49.86 ppm), DMSO-*d6* (δH = 2.50 ppm/δC = 39.52 ppm). Signal patterns are described as singlet (s), doublet (d), triplet (t), quartet (q), multiplet (m), broad singlet (bs), or a combination of the listed splitting patterns. The coupling constants (*J*) are measured in hertz (Hz).

### Synthesis of Compound 1*

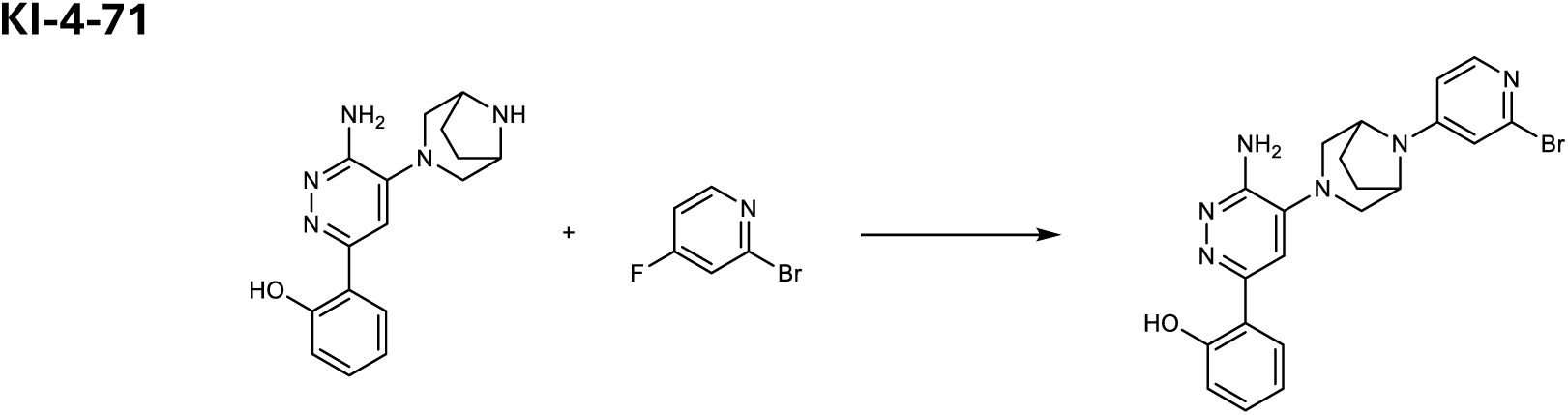

To a solution of a mixture of 2-[6-amino-5-(3,8-diazabicyclo[3.2.1]octan-3-yl)pyridazin-3-yl]phenol (1.0 g, 3.36 mmol, 1.0 eq.) in DMSO (10 mL) was added 2-bromo-4-fluoro-pyridine (651 mg, 3.7 mmol, 1.1 eq.) and triethylamine (2.3 mL, 16.8 mmol, 5.0 eq.). After being stirred at 130 °C for 6 hrs, the reaction was diluted with EtOAc and water. The mixture was extracted with EtOAc. The organic layer was dried over Na2SO4, filtered, and concentrated. The residue was suspended in CHCl_3_/EtOAc (1/10) and filtered. The solid material was collected to give Compound 1* (760 mg, 1.68 mmol) in 50% yield.

**HRMS ESI^+^**, m/z calcd for C21H21BrN6O [M+H]^+^ 453.1033; found 453.1038.

**1H NMR (500 MHz, DMSO-*d6*)** δ 14.11 (s, 1H), 7.87-7.93 (m, 2H), 7.52 (s, 1H), 7.22 (dd, *J* = 7.8, 7.8 Hz, 1H), 7.01 (s, 1H), 6.89-6.82 (m, 3H), 5.98 (s, 2H), 4.60 – 4.55 (m, 2H), 3.27 (d, *J* = 11.7 Hz, 2H), 3.00 (d, *J* = 11.7 Hz, 2H), 2.22-2.15 (m, 2H), 2.00-1.93 (m, 2H).

**13C NMR (126 MHz, DMSO-*d6*)** δ 158.51, 154.53, 153.70, 152.41, 149.99, 143.08, 140.45, 130.27, 126.43, 118.50, 117.72, 117.40, 111.60, 111.42, 109.52, 53.70, 50.92, 27.03.

### Synthesis of Compound 1

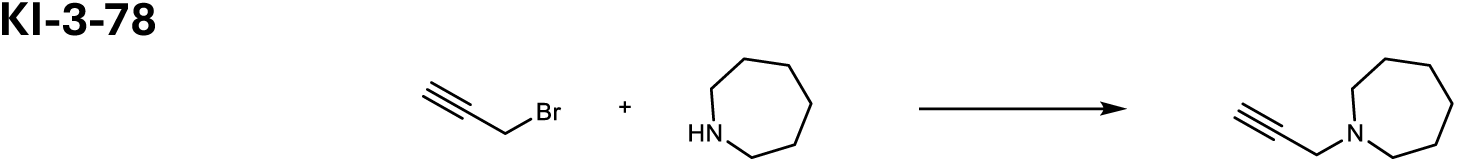

To a suspension of potassium carbonate (1.9 g, 13.4 mmol, 2.0 eq.) and azepane (3.8 mL, 33.6 mmol, 5.0 eq.) in DCM was added 3-bromoprop-1-yne (1.0 g, 6.72 mmol, 1.0 eq.) at 0 °C. The reaction was stirred at room temperature for 16 hrs. Then the mixture was filtered, and the filtrate was concentrated. The residue was suspended in heptane and then filtered. The filtrate was concentrated to give the product as a mixture of bis-propargyl azepane and azepane, which was used in the next reaction without further purifications.

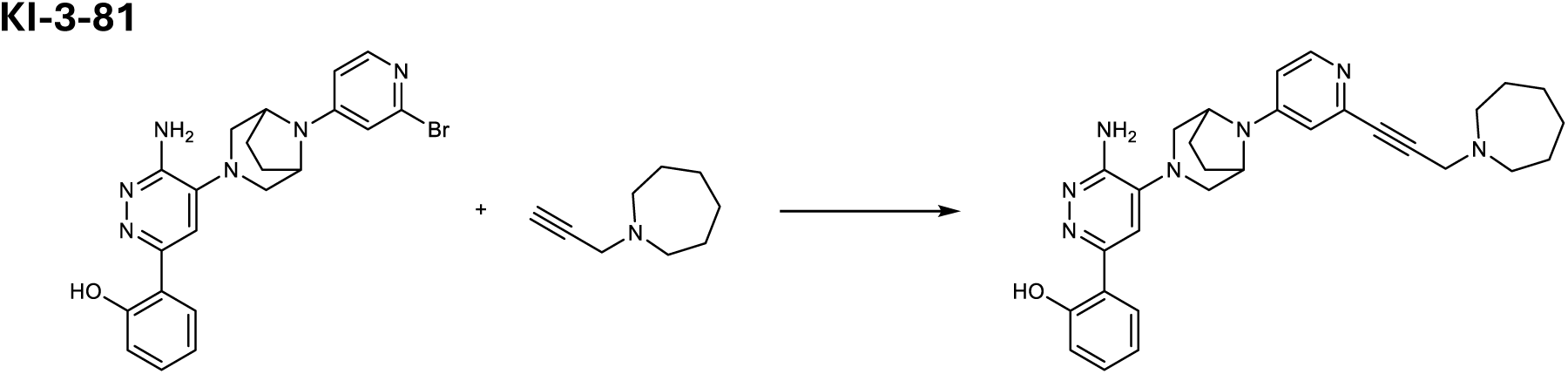

To a solution of 1-prop-2-ynylazepane (45 mg, crude mixture from the previous reaction), 2-[6-amino-5-[3-(2-bromo-4-pyridyl)-3,8-diazabicyclo[3.2.1]octan-8-yl]pyridazin-3-yl]phenol (15 mg, 0.033 mmol, 1.0 eq), Copper(I) iodide (0.63 mg, 3.3 μmol, 0.1 eq.), and N, N-diisopropylethylamine (0.06 mL, 0.33 mmol, 10.0 eq.) in THF (2.0 mL) was added [1,1′-[1,1′-Bis(diphenylphosphino)ferrocene]dichloropalladium(II) (4.9 mg, 6.6 μmol, 0.2 eq.). After being stirred at 80 °C for 4 hrs under nitrogen atmosphere, the reaction was diluted with EtOAc and water. The mixture was extracted with EtOAc, washed with saturated aqueous NaCl, dried over Na2SO4, filtered, and concentrated. Column chromatography (ODS, MeCN/H2O with 0.1% HCOOH) of the residue gave the product, which was purified with preparative HPLC (MeCN/H2O with 0.1% NH4OH) to give Compound 1 (4.8 mg, 9.3 μmol) in 28% yield.

**HRMS ESI^+^**, m/z calcd for C30H36N7O [M+H]^+^ 510.2976; found 510.2989.

**1H NMR (500 MHz, CD_3_OD)** δ 8.06 (d, *J* = 6.1 Hz, 1H), 7.73 (m, 1H), 7.47 (s, 1H), 7.22 (m, 1H), 6.99 (m, 1H), 6.92 – 6.84 (m, 2H), 6.80 (m, 1H), 4.71 (s, 1H), 4.57 (s, 2H), 3.60 (s, 2H), 3.40 – 3.32 (m, 2H), 3.12-3.05 (m, 2H), 2.84-2.79 (m, 4H), 2.29-2.21 (m, 2H), 2.17-2.11 (m, 2H), 1.78-1.69 (m, 4H), 1.69-1.61 (m, 4H).

**^13^C NMR (126 MHz, CD_3_OD)** δ 159.74, 155.98, 155.93, 153.13, 150.66, 144.17, 142.64, 131.66, 127.26, 120.17, 119.18, 118.66, 113.83, 113.27, 110.07, 85.81, 85.27, 56.52, 55.51, 52.64, 49.10, 28.49, 28.41, 27.69.

### Synthesis of Compound 2

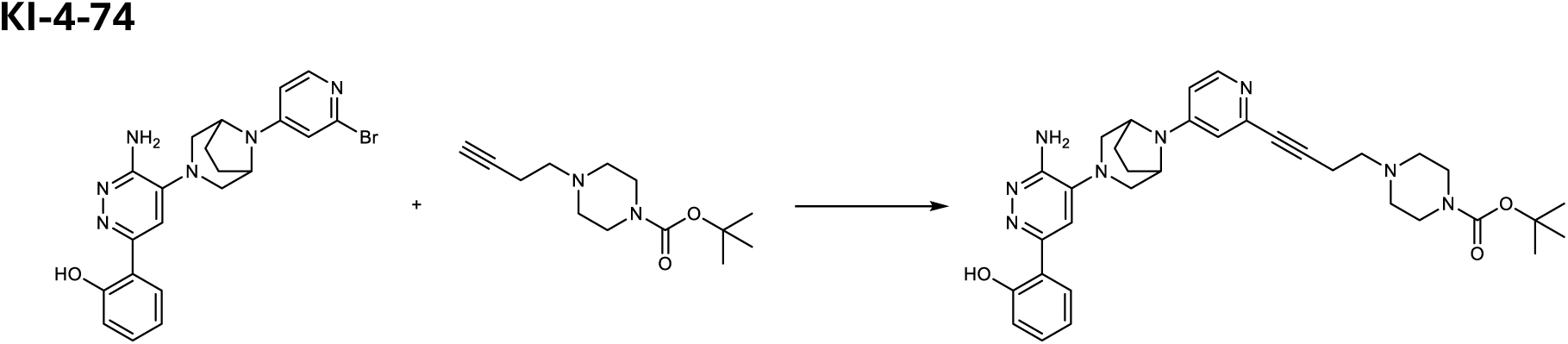

To a solution of 2-[6-amino-5-[8-(2-bromo-4-pyridyl)-3,8-diazabicyclo[3.2.1]octan-3-yl]pyridazin-3-yl]phenol (50 mg, 0.11 mmol, 1.0 eq.), tert-butyl 4-but-3-ynylpiperazine-1-carboxylate (237 mg, 0.99 mmol, 9.0 eq.), Copper(I) iodide (6.3 mg, 0.033 mmol, 0.3 eq.), and N, N-diisopropylethylamine (0.38 mL, 2.2 mmol, 20 eq.) in THF (2.0 mL) was added [1,1′-Bis(diphenylphosphino)ferrocene]dichloropalladium(II) (16 mg, 0.022 mmol, 0.20 eq.). After being stirred at 80 °C for 3 hrs under nitrogen atmosphere, the reaction was diluted with EtOAc and water. The mixture was extracted with EtOAc, washed with saturated aqueous NaCl, dried over Na2SO4, filtered, and concentrated. Column chromatography (ODS 13 g, MeCN/H2O with 0.1% HCOOH) of the residue gave the impure tert-butyl 4-[4-[4-[3-[3-amino-6-(2-hydroxyphenyl)pyridazin-4-yl]-3,8-diazabicyclo[3.2.1]octan-8-yl]-2-pyridyl]but-3-ynyl]piperazine-1-carboxylate (70 mg), which was used in the next reaction without further purifications.

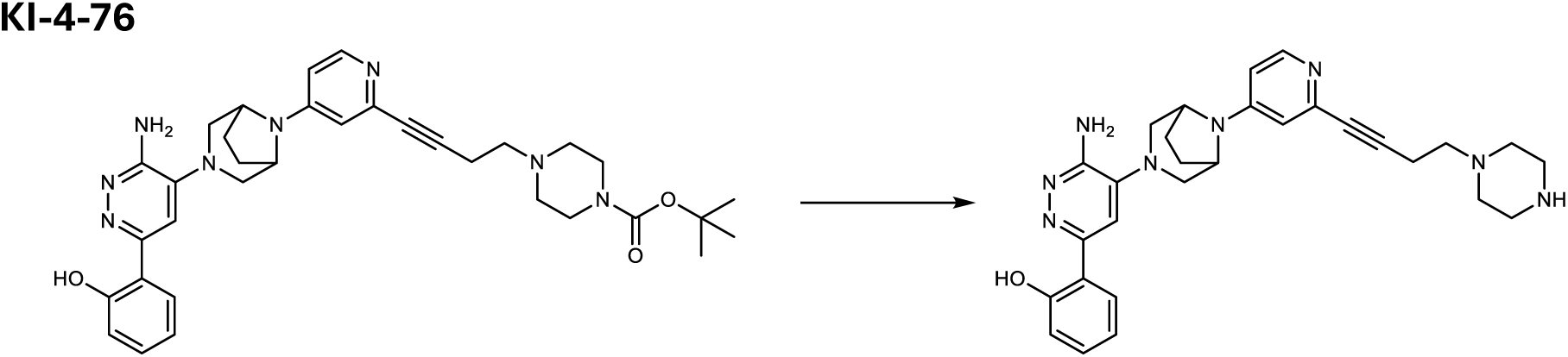

To a solution of tert-butyl 4-[4-[4-[3-[3-amino-6-(2-hydroxyphenyl)pyridazin-4-yl]-3,8-diazabicyclo[3.2.1]octan-8-yl]-2-pyridyl]but-3-ynyl]piperazine-1-carboxylate (a mixture from the previous reaction, calculated as 0.11 mmol, 1.0 eq.) in DCM (0.5 mL) was added 2,2,2-trifluoroacetic acid (0.5 mL). The reaction mixture was being stirred for 30 min and then evaporated. The residue was filtered and purified with preparative HPLC (MeCN/H2O with 0.1% NH4OH) to give 2-[6-amino-5-[8-[2-(4-piperazin-1-ylbut-1-ynyl)-4-pyridyl]-3,8-diazabicyclo[3.2.1]octan-3-yl]pyridazin-3-yl]phenol (20 mg, 0.0398 mmol) 36% yield in 2 steps.

**HRMS ESI^+^**, m/z calcd for C29H34N8O [M+H]^+^ 511.2983; found 511.2941.

**1H NMR (500 MHz, CD_3_OD)** δ 8.02 (d, *J* = 6.1 Hz, 1H), 7.71 (dd, *J* = 8.1, 1.8 Hz, 1H), 7.44 (s, 1H), 7.21 (ddd, *J* = 8.5, 7.3, 1.6 Hz, 1H), 6.94 (d, *J* = 2.5 Hz, 1H), 6.91 – 6.84 (m, 2H), 6.75 (dd, *J* = 6.2, 2.6 Hz, 1H), 4.54 (brd, *J* = 3.8 Hz, 2H), 3.36-3.32 (m, 2H), 3.06 (d, *J* = 11.6 Hz, 2H), 2.85 (m, 4H), 2.71 – 2.60 (m, 4H), 2.52 (m, 4H), 2.27-2.22 (m, 2H), 2.15-2.08 (m, 2H).

**^13^C NMR (126 MHz, CD_3_OD)** δ 159.76, 155.95, 155.90, 153.11, 150.51, 144.66, 142.60, 131.66, 127.27, 120.15, 119.20, 118.68, 113.68, 113.22, 109.87, 88.81, 82.07, 58.42, 55.48, 54.54, 52.54, 46.07, 28.41, 17.68.

### Synthesis of Compound 3

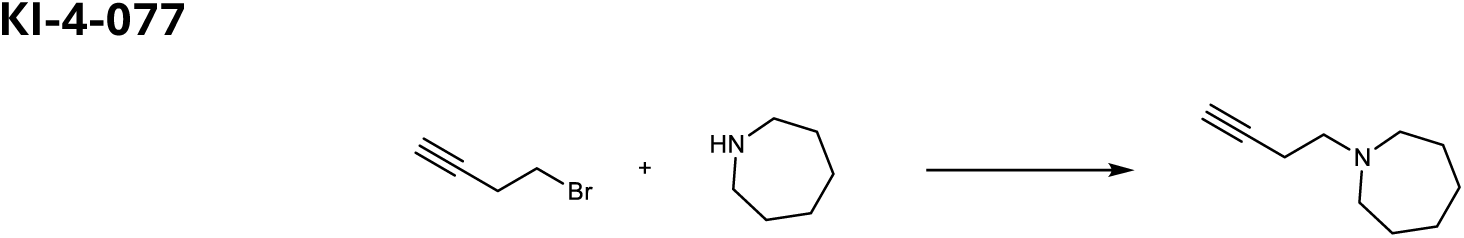

To a suspension of azepane (0.93 mL, 8.3 mmol, 1.1 eq.) and potassium carbonate (1.6 g, 11 mmol, 1.5 eq.) in MeCN (10 mL) was added 4-bromobut-1-yne (0.71 mL, 7.5 mmol, 1.0 eq.). After being stirred at 60 °C for 18 hrs, the reaction was diluted with DCM and water. The mixture was extracted with DCM, dried over Na2SO4, filtered, and concentrated to give impure 1-but-3-ynylazepane (410 mg), which was used in the next reaction without further purification.

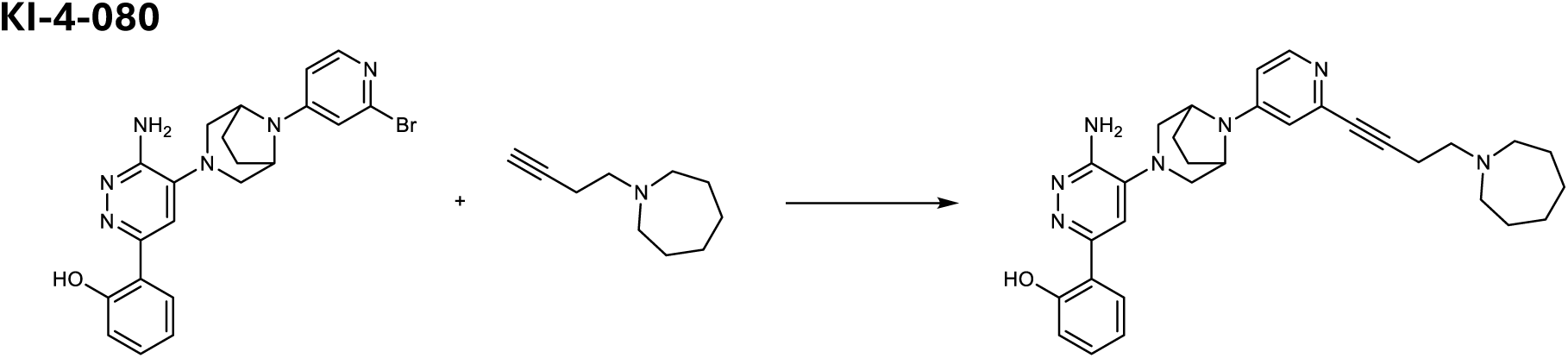

To a solution of 2-[6-amino-5-[8-(2-bromo-4-pyridyl)-3,8-diazabicyclo[3.2.1]octan-3-yl]pyridazin-3-yl]phenol (50 mg, 0.11 mmol, 1.0 eq.), a crude 1-but-3-ynylazepane (152 mg, calculated as 1.01 mmol, 9.1 eq.), Copper(I) iodide (6.3 mg, 0.033 mmol, 0.3 eq.), and N,N-Diisopropylethylamine (0.38 mL, 2.21 mmol, 20 eq.) in THF (2.0 mL) was added [1,1′-Bis(diphenylphosphino)ferrocene]dichloropalladium(II) (16 mg, 0.022 mmol, 0.2 eq.). After being stirred at 80 °C for 3 hrs under nitrogen atmosphere, the reaction was diluted with EtOAc and water. The mixture was extracted with EtOAc, washed with saturated aqueous NaCl, dried over Na2SO4, filtered, and concentrated. Column chromatography (ODS, MeCN/H2O with 0.1% HCOOH) of the residue gave the product, which was repurified with HPLC (MeCN/H2O with 0.1% NH4OH) to give Compound 3(12.1 mg, 0.023 mmol) in 21% yield.

**HRMS ESI^+^**, m/z calcd for C31H37N7O [M+H]^+^ 524.3132; found 524.3141.

**1H NMR (500 MHz, CD_3_OD)** δ 8.03 (d, *J* = 6.1 Hz, 1H), 7.72 (dd, *J* = 8.0, 1.6 Hz, 1H), 7.45 (s, 1H), 7.22 (ddd, *J* = 8.5, 7.2, 1.6 Hz, 1H), 6.94 (d, *J* = 2.4 Hz, 1H), 6.91-6.84 (m, 2H), 6.76 (dd, *J* = 6.1, 2.5 Hz, 1H), 4.54 (m, 2H), 3.33 (d, *J* = 11.6 Hz, 2H), 3.07 (d, *J* = 11.6 Hz, 2H), 2.83 (dd, *J* = 8.3, 6.8 Hz, 2H), 2.77 – 2.71 (m, 4H), 2.62 (dd, *J* = 8.4, 6.9 Hz, 2H), 2.23 (m, 2H), 2.15 (m, 2H), 1.71-1.65 (m, 4H), 1.65-1.58 (m, 4H).

**^13^C NMR (126 MHz, CD_3_OD)** δ 159.77, 155.97, 155.93, 153.12, 150.50, 144.74, 142.63, 131.65, 127.26, 120.15, 119.20, 118.67, 113.66, 113.23, 109.84, 89.07, 82.06, 57.75, 56.21, 55.48, 52.55, 28.41, 28.33, 28.00, 18.03.

### Synthesis of Compound 4

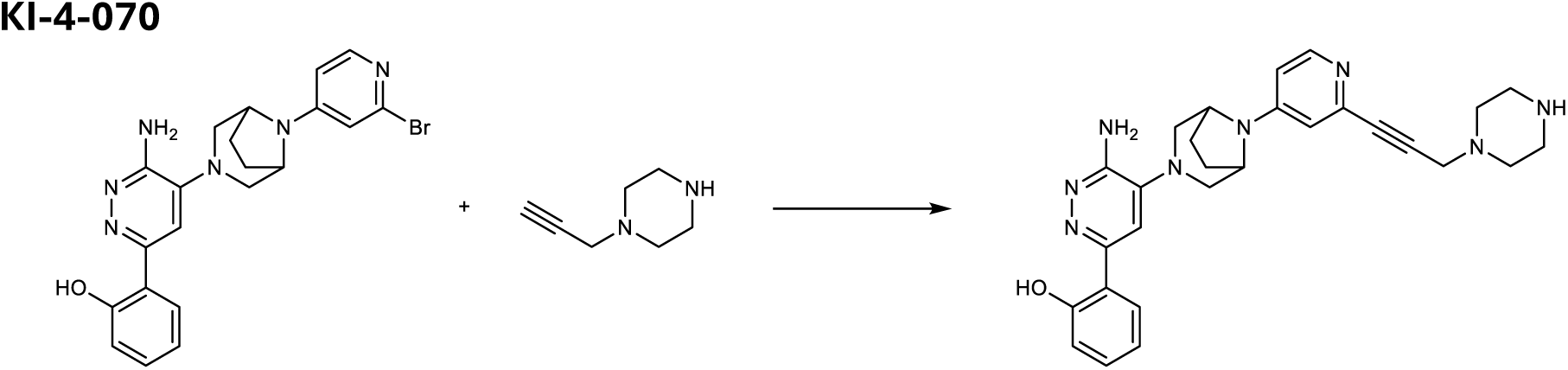

To a solution of 2-[6-amino-5-[8-(2-bromo-4-pyridyl)-3,8-diazabicyclo[3.2.1]octan-3-yl]pyridazin-3-yl]phenol (30 mg, 0.066 mmol, 1.0 eq.), 1-prop-2-ynylpiperazine (75 mg, 0.605 mmol, 9.1 eq.), Copper(I) iodide (3.8 mg, 0.020 mmol, 0.3 eq.), and N, N-diisopropylethylamine (0.23 mL, 1.32 mmol, 20 eq.) in THF (2.0 mL) was added [1,1′-Bis(diphenylphosphino)ferrocene]dichloropalladium(II) (9.7 mg, 0.013 mmol, 0.2 eq.). After being stirred at 80 °C for 3 hrs under nitrogen atmosphere, the reaction was diluted with EtOAc and was added water. The mixture was extracted with EtOAc, washed with saturated aqueous NaCl, dried over Na2SO4, filtered, and concentrated. Column chromatography (ODS, MeCN/H2O with 0.1% HCOOH) of the residue gave the product, which was re-purified with preparative HPLC (MeCN/H2O with 0.1% NH4OH) to give Compound 4(12.4 mg, 0.025 mmol) in 38% yield.

**HRMS ESI^+^**, m/z calcd for C28H32N8O [M+H]^+^ 497.2772; found 497.2783.

**1H NMR (500 MHz, CD_3_OD)** δ 8.06 (d, *J* = 6.0 Hz, 1H), 7.73 (dd, *J* = 8.0, 1.6 Hz, 1H), 7.47 (s, 1H), 7.22 (ddd, *J* = 8.5, 7.2, 1.6 Hz, 1H), 7.01 (d, *J* = 2.5 Hz, 1H), 6.91-6.85 (m, 2H), 6.80 (dd, *J* = 6.1, 2.6 Hz, 1H), 4.58 (dd, *J* = 4.7, 2.5 Hz, 2H), 3.54 (s, 2H), 3.39 – 3.32 (m, 2H), 3.09 (d, *J* = 11.5 Hz, 2H), 2.89 (m, 4H), 2.66 (m, 4H), 2.29-2.22 (m, 2H), 2.17-2.09 (m, 2H).

**13C NMR (126 MHz, CD_3_OD)** δ 159.75, 155.99, 155.93, 153.13, 150.68, 144.03, 142.63, 131.66, 127.26, 120.16, 119.19, 118.67, 113.88, 113.27, 110.12, 86.20, 84.58, 55.50, 53.51, 52.63, 48.43, 46.07, 28.41.

### Synthesis of Compound 5

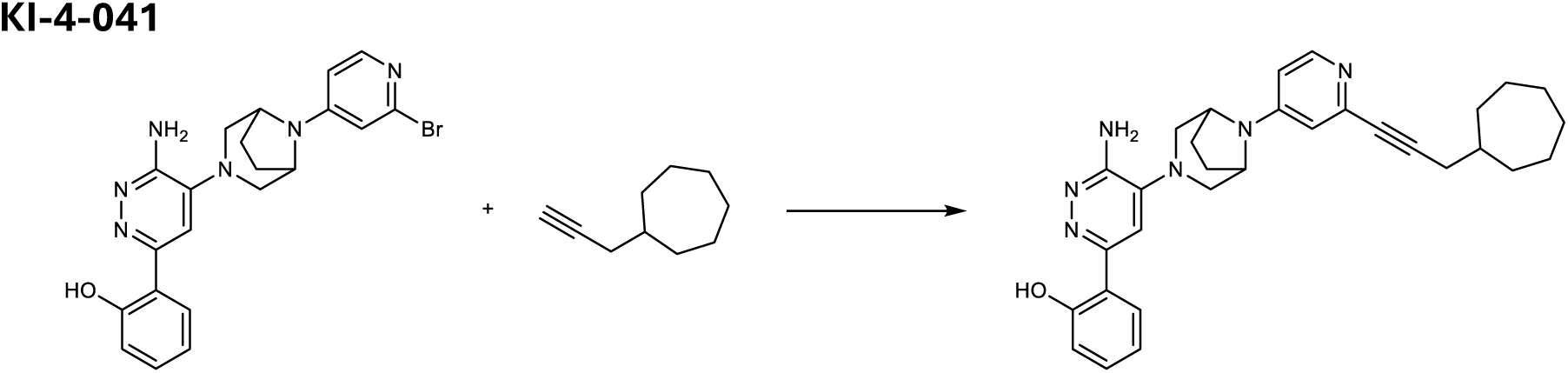

To a solution of 2-[6-amino-5-[8-(2-bromo-4-pyridyl)-3,8-diazabicyclo[3.2.1]octan-3-yl]pyridazin-3-yl]phenol (20 mg, 0.044 mmol, 1.0 eq.), prop-2-ynylcycloheptane (50 mg, 0.37 mmol, 8.3 eq.), Copper(I) iodide (2.5 mg, 0.0132 mmol, 0.3 eq.), and N, N-diisopropylethylamine (0.077 mL, 0.44 mmol, 10 eq.) in THF (2.0 mL) was added [1,1′-Bis(diphenylphosphino)ferrocene]dichloropalladium(II) (6.5 mg, 0.0088 mmol, 0.2 eq). After being stirred at 80 °C for 4 hrs under nitrogen atmosphere, the reaction was diluted with EtOAc and water. The mixture was extracted with EtOAc, washed with saturated aqueous NaCl, dried over Na2SO4, filtered, and concentrated. Column chromatography (ODS, MeCN/H2O with 0.1% HCOOH) of the residue gave the product, which was re-purified with preparative HPLC (MeCN/H2O with 0.1% NH4OH) to give COMPOUND 5(10.0 mg, 0.020 mmol) in 45% yield.

**HRMS ESI^+^**, m/z calcd for C31H36N6O [M+H]^+^ 509.3023; found 509.3030.

**1H NMR (500 MHz, CD_3_OD)** δ 8.02 (d, *J* = 6.1 Hz, 1H), 7.76 – 7.71 (m, 1H), 7.47 (s, 1H), 7.22 (ddd, *J* = 7.7, 7.7, 1.6 Hz, 1H), 6.94 – 6.85 (m, 3H), 6.76 (m, 1H), 4.56 (t, J = 3.8 Hz, 2H), 3.34 (d, *J* = 11.5 Hz, 2H), 3.08 (d, *J* = 11.5 Hz, 2H), 2.34 (d, *J* = 6.8 Hz, 2H), 2.27-2.20 (m, 2H), 2.17-2.10 (m, 2H), 1.94-1.85 (m, 2H), 1.77 (m, 1H), 1.74-1.66 (m, 2H), 1.66 – 1.57 (m, 2H), 1.57-1.42 (m, 4H), 1.41-1.31 (m, 2H).

**13C NMR (126 MHz, CD_3_OD)** δ 159.75, 155.98, 155.95, 153.13, 150.44, 145.02, 142.67, 131.65, 127.27, 120.17, 119.19, 118.66, 113.51, 113.25, 109.67, 90.57, 81.88, 55.47, 52.60, 40.54, 35.50, 29.40, 28.56, 28.42, 27.42.

### Cell culture

HEK293, HEK293T and HCT116, originally sourced from ATCC, were provided by the MRC PPU reagents facility at the University of Dundee. KBM7 iCas9 cells were a gift from Johannes Zuber. HEK293T, HCT-116, Lenti-X 293T lentiviral packaging cells (Clontech) and Flp-In T-REx-293 (Thermo Fisher Scientific) cells were maintained in high glucose Dulbecco’s Modified Eagle’s Medium (DMEM, Gibco/Sigma-Aldrich). KBM7 cells were maintained in Iscove’s Modified Dulbecco’s Medium (IMDM, Sigma-Aldrich). Both DMEM and IMDM were supplemented with 10% Fetal Bovine Serum (FBS) and 1% (v/v) penicillin/streptomycin (all supplied by Thermo Fisher Scientific/Sigma-Aldrich). Cell lines were grown in a humidified incubator at 37 °C and 5% CO_2_, routinely tested for mycoplasma contamination, and authenticated by short tandem repeat profiling.

### Plasmids and oligonucleotides

Generation of the human CRL-focused sgRNA library used for SMARCA2 stability reporter screens, lentiviral sgRNA expression vector used for DCAF16 gene knockout, and viral vectors used for the engineering of inducible Cas9 cell lines have been previously described. The Fluorescent protein stability reporter for SMARCA2 was generated by subcloning the bromodomain (residues 62-235) of SMARCA2 from SMARCA2 pDONR223 (a gift from Mikko Taipale) and inserting it into a pRRL lentiviral vector, fused to a 3xV5 tag and mTagBFP at the C-terminus, and coupled to mCherry with a P2A self-cleaving peptide for normalization. The same SMARCA2 bromodomain sequence was used to create a fusion construct with a miniTurboID enzyme (47) at the N-terminus, plus a flexible linker, and a HiBiT tag. This construct was cloned into the pcDNA5/FRT/TO vector (V652020, ThermoFischer), generously provided by Dr. Ronald Hay lab, and verified by sequencing to ensure proper insertion and orientation.

The Fluorescent protein stability reporter for BRD4 tandem bromodomains has been previously described (35). To generate knockout cell lines, DCAF16 and FBXO22 sgRNA were cloned into a pSpCas9(BB)-2A-EGFP vector from Feng Zhang (PX458, Addgene #48138). For reconstitution experiments, wild-type and cysteine mutants of FBXO22 fused to a 2xHA tag at the N-terminus in a pLEX lentiviral vector were previously described (26). Exogenous expression of DCAF16 was achieved from a pRRL lentiviral vector with a 3xFlag tag at the N-terminus, as previously described (35). Expression clones of DCAF16 mutants were generated from the wild-type 3xFlag tag vector using the Q5 site-directed mutagenesis Kit (New England Biolabs, E0552), according to the manufacturer’s instructions. For site directed mutagenesis studies, amino acids 2-216 of DCAF16 were selected for site saturation library design and cloning into a pRRL-SFFV-EGFP-cpHalo-(GSG)x9-DCAF16 backbone by GenScript. Quality control of the library by Next-Generation Sequencing was also performed by GenScript and resulted in a library coverage of 99.80% and 534x. For immunoprecipitation experiments expression plasmids pRK5-HA-DCAF16 and pCMV5 HA-FBXO22 were kindly provided by Dr. Xiaoyu Zhang (Northwestern University, USA), or acquired from the MRC PPU reagents facility at the University of Dundee (DU42092). LgBiT and HaloTag-Ubiquitin plasmids were purchased from Promega(N2681, N2721 respectively). Flp-recombinase expression vector (pOG44) was kindly provided by Dr. Ronald Hay lab. All plasmids and sgRNAs used in this study are shown in [Extended Data Table 1]. The CRL-focused sgRNA library used for FACS-based CRISPR–Cas9 screens is shown in [Supplementary Table 2].

### Lentivirus production and transduction

Semiconfluent Lenti-X cells in 10 cm dishes were co-transfected with 1 µg of the envelope plasmid pMD2.G (Addgene #12259), 2 µg of packaging plasmid psPAX2 (Addgene #12260), and 4 µg of the lentiviral plasmid using polyethylenimine (PEI MAX MW 40,000, Polysciences). 3 days after transfection, supernatant containing virus was collected and filtered through a 0.45 mm filter. Target cells were infected in the presence of 4 μg ml^−1^ polybrene (szabo scandic, SACSC-134220).

### Generation of monoclonal HEK293T knockout cell lines

Semiconfluent HEK293T cells in 6-well plates were transfected with 2.5µg of a pSpCas9(BB)-2A-EGFP plasmid containing either DCAF16 or FBXO22 sgRNA using Lipofectamine 2000 (Thermo Fisher Scientific, 11668027). 2 days after transfection, single cells expressing EGFP were sorted into 96-well plates and allowed to grow for 2 weeks. Monoclonal cell colonies were verified for FBXO22 or DCAF16 knockouts via Western Blotting and preventing BRD4 degradation in the presence of IBG3 (35). As described with single knockout cell lines, double knockout clonal cells were generated starting from a clonal population of DCAF16 knockout cells transfected with an FBXO22 sgRNA plasmid.

### Generation of SMARCA HiBiT cell lines and knockout clones

SMARCA2 and SMARCA4 HiBiT CRISPR-Cas9 knock-in cell lines were generated via ribonucleoprotein (RNP) transfection according to the manufacturer’s instructions for single-stranded DNA oligonucleotides (ssODNs, IDT). In brief, ssODNs served as donor templates, while recombinant SpCas9 Nuclease (IDT) was complexed with target-specific sgRNAs [Extended Data Table 1]. For RNP transfection, 1 x 10^6^ HEK293T cells were resuspended in MaxCyte buffer (cytiva) and mixed with SMARCA2- or SMARCA4-specific HiBiT ssODNs and RNPs. The mixture was filled into an electroporation cuvette to a final volume of 25 µl and electroporated with a MaxCyte ExPERT ATx. Following electroporation, cells were immediately transferred to a 6-well plate as droplets and allowed to recover at 37°C and 5% CO2 for 20 minutes. Following recovery, 2 ml of pre-warmed DMEM (10% FBS, 1% penicillin–streptomycin) supplemented with Alt-R HDR Enhancer V2 (IDT) was added to the cells. The following day, media was exchanged and cells were allowed to expand. Three days after electroporation, the edited cells pools were analyzed with the HiBiT lytic assay (Promega, N3030) to detect luminescence and the HiBiT blotting system (Promega, N2410) to detect proteins tagged with HiBiT at the correct weight via Western Blotting. To obtain monoclonal populations, single cells were sorted into 96-well plates and allowed to grow for 2 weeks. Clones were expanded and validated as homozygous SMARCA2/SMARCA4-HiBiT clones by genotyping and by performing a lytic HiBiT degradation assay with the SMARCA-specific PROTAC, ACBI1(48).

HiBiT-SMARCA4 HEK293T single knockout cell lines were generated via RNP electroporation using target-specific sgRNA (IDT) and spCas9 (CAS9PROT, Sigma-Aldrich) (Extended Data Table 1). After forming the RNP complex via 20 minutes incubation at room temperature, 4 x 10^5 HiBiT-SMARCA4 HEK293T cells were resuspended in Neon NxT Resuspension R Buffer (N10025, Thermo Fisher Scientific) along with the RNP complex and electroporated using a 10 µl Neon Electroporation System cuvette tip (Thermo Fisher Scientific). Immediately following electroporation, the cells were placed in pre-warmed DMEM supplemented with 10% FBS and 1% (v/v) penicillin/streptomycin. Three days after electroporation, individual cells were sorted into 96-well plates and allowed to grow for two weeks. To confirm the successful knockout of the FBXO22 or DCAF16 genes, Western Blotting was employed to verify the gene knockouts and to assess the prevention of BRD4 degradation in the presence of IBG3. Finally, double knockout clonal cells were generated starting from a clonal population of FBXO22 knockout cells that were subsequently electroporated with a DCAF16 sgRNA.

### Western Blotting

HEK293T and HCT116 cells were plated in 6 or 12-well plates depending on experimental setup at 0.5 - 0.6 x 10^6^ cells per ml. Stock solutions of compounds were prepared in DMSO at a concentration of 10 mM and stored at −80 °C. In all experiments, media from cell seeding was exchanged with fresh DMEM media containing working dilutions of experimental compounds. For the Ubiquitin-Proteasome inhibition assay, cells were co-treated for 18 hours with compound 1 and 0.5 µM of TAK243, MLN4924, or Carfilzomib.

For cell collection, cells were washed once with PBS before lysis on ice for 20 minutes with RIPA buffer (150 mM NaCl, 1% Triton X-100, 0.5% sodium deoxycholate, 0.1% SDS, 50 mM Tris-HCl pH 8) supplemented with benzonase (1:1000, Sigma-Aldrich, 70746-3), HALT EDTA-free protease inhibitor cocktail (1:100, Thermo Fisher Scientific, 78437), and 20 mM DTT. After removal of the insoluble fraction by centrifugation at 15,000 g at 4 °C for 15 min, supernatants were stored at −80°C. Protein concentration was determined by bicinchoninic acid (BCA) assay (#23225, Thermo Fisher Scientific). For HEK293, lysates were prepared with 4 x Bolt LDS sample buffer (Thermo Fisher, B0008), heated at 95°C for 5 minutes, and then run on Bolt 4–12% bis-tris gels (Thermo Fisher Scientific) with Bolt MES SDS running buffer (Thermo Fisher Scientific). For HCT116, 4× NuPAGE LDS sample buffer was added to the cell lysates of (NP0007, Thermo Fisher Scientific) supplemented with 10% DTT and heated at 95 °C for 5 min. Samples (20 to 30 µg) were analyzed via protein electrophoresis with a NuPAGE MOPS SDS running buffer (Thermo Fisher Scientific).Proteins were transferred to nitrocellulose membranes (GE Healthcare, Amersham Protran Supported 0.4 mm NC), blocked for 1 hour with 5% milk in TBS-T at room temperature, before incubation with primary antibodies at 4 °C, overnight. The following primary antibodies were used: HSP90 (1:1000, Cell signalling Technology, 4877), FBXO22 (1:200, Santa Cruz, sc-100736), SMARCA2 (1:1000, Bethyl Laboratories, A301-015A), SMARCA4 (1:1000, Bethyl Laboratories, A300-813A), HA epitope (3F10) (1:1000, Merck, 11867423001), HiBiT (1:1000, Promega, N7200), IRDye® 800CWStreptavidin (1:1000, Licor, 926-32230), hFABTM rhodamine anti-tubulin (1:5000, Bio-Rad, 12004165) and FLAG (1:1000, Sigma-Aldrich, F1804-200UG). Following overnight treatment, membranes were washed in TBS-T and incubated with horseradish peroxidase (HRP)-conjugated or fluorescence 800CW secondary antibodies for 1-2 hours at room temperature. The following secondary antibodies were used: HRP anti-rabbit IgG (1:5000, Cell Signaling Technology, 7074), HRP anti-mouse IgG (1:5000, Cell Signaling Technology, 7076), IRDye 800CW anti-rabbit (1:5000, Licor, 926-32211), IRDye 800CW anti-mouse (1:5000, Licor, 926-32210) and IRDye 800CW anti-rat (1:5000, Licor, 926-32219) and HRP anti-mouse IgG (1:5000, Cell Signaling Technology, 7076). Lastly, membranes were washed again with TBS-T and then imaged on ChemiDoc Touch imaging system (Bio-Rad) operated on Image Lab software (v2.4.0.03).

### HiBiT Degradation assays

SMARCA4 HiBiT cells were plated in 384 or 96 well plates at a density of respectively 3 x 10^3^ or 3 x10^4^ cells per well. The following day, compounds were dispensed either manually (preparing a 5x stock of compounds and dispensing then for a final concentration of 1x) or with a Labcyte Echo550/555 Liquid Handler dispenser (Beckman Coulter Life Sciences) in a final volume of 100 ul. Each treatment was performed in biological triplicates with technical replicates for each. Cells were treated for 16, 18 or 24 hours, as indicated in the respective figure legends, before lysis using the HiBiT lytic assay buffer (Promega, N3030) according to the manufacturer’s instructions. Luminescence in each plate was measured using a Perkin Elmer Victor X3 plate reader. Treated wells were normalized to a DMSO-only control and analysed using GraphPad Prism (v10.0.3) via fitting of non-linear regression curves for extraction of DC_50_ values.

### NanoBRET Ubiquitination and Kinetic degradation assay

For NanoBRET Ubiquitination assays, 0.8 x 10^6^ HiBiT-tagged HEK293T cells were seeded in 6-well plates. The following day, 1 µg of LgBiT (Promega, N2681) and 1 µg HaloTag-Ubiquitin cDNA (Promega, N2721) were transfected using FuGENE HD (Promega, E2311) at a 3:1 transfection reagent:plasmid ratio. After 8 hours, cells were trypsinized and resuspended in phenol red-free OptiMEM (Gibco) supplemented with 4% FBS and seeded in 96-well plates at a density of 0.2 x 10^6^ cells per ml in the presence or absence of 0.1 mM HaloTag NanoBRET 618 ligand (Promega, G9801). Following overnight incubation, experimental compounds were added at the indicated concentrations. After 24 hours of compound treatment, NanoBRET™ Nano-Glo Substrate (Promega, N1571) was diluted in Optimen and added to cells at a 1x final concentration. 96-well plates were analysed using a PHERAstar (BMG Labtech) plate reader operated on PHERAstar software (firmware v1.33) for NanoBRET ratio metric calculation (460 nm donor and 618 nm acceptor emissions). Data was processed by subtracting NanoBRET ligand-free controls before plotting NanoBRET signal in GraphPad Prism (v9.5.1).

Kinetic degradation assays were conducted using HiBiT-tagged SMARCA4 HEK293T cells that were transfected with exogenous LgBiT, following the same protocol as described for the NanoBRET Ubiquitination assays. The cells were incubated with Endurazine substrate (1:100, N2570, Promega) for 2.5 hours at 37 °C before the addition of 10x concentrations of experimental compounds. Luminescence measurements were taken every 10 minutes for a duration of 24 hours using a GloMAX Discover microplate reader (Promega). The data were normalized to DMSO-only controls and plotted as luminescence signal over time using GraphPad Prism (v9.5.1).

### Generation of 293 Flp-In T-REx pools

A total of 1 x 10^6 Flp-In™ T-REx™ 293 cells (R78007, ThermoFischer) were co-transfected using Lipofectamine 3000 using 3.6 µg of a Flp-recombinase expression vector (pOG44, Thermo Fisher Scientific) and 0.4 µg of a pcDNA5/FRT/TO plasmid containing a HiBiT-SMARCA2 Bromodomain with miniTurbo on the Ńterminus. 24 hours following transfection, polyclonal cell pools were washed and replaced with complete media and the following day, selected for 2 weeks with 200 µg ml−1 hygromycin B (Thermo Fisher Scientific, 10687010). Cells were expanded, then construct expression and biotinylation activity was induced with 1 µg ml−1 Doxycycline (D9891, Merck) and Biotin (B4501, Merck) to be validated by Western blotting as described above.

### BioID of SMARCA2 BD

Flp-In™ T-REx™ 293 stable cell line containing miniTurbo-SMARCA2 Bromodomain was divided into three treatment conditions, with each condition consisting of three 10-cm dishes at ∼70% confluency. Expression of the fusion protein was induced by 1 µg ml^−1^ doxycycline (Dox) treatment for a total of 20,5 hours. 12 hours after Dox induction, cells were then pre-treated with 10 µM MLN4924 Neddylation inhibitor (5054770001,Merck) for 30 mins followed by addition of 10 µM Compound 1, 10 µM Compound 2 or DMSO for 8 hours. Biotin (100 µM final concentration) was added to cells undergoing treatment with compounds or DMSO for 2 hours. At the endpoint, the culture medium was removed, and cells were collected with PBS by scraping in 15-cm tubes pelleted by centrifugation at 600g for at least 5 min and washed for a second time in PBS. BioID was performed as described previously (49, 50). Cells were lysed with RIPA lysis and extraction buffer (25 mM Tris-HCl pH 7.6, 150 mM NaCl, 1% NP-40, 1% sodium deoxycholate, 0.1% SDS) (#89900,Thermo Fisher Scientific) supplemented with complete EDTA-free protease inhibitor cocktail (11873580001,Roche) and 1:1000 benzonase nuclease (70746-3, Sigma-Aldrich). Lysates were incubated at 4 °C for 15 minutes, then centrifuged at 13,000g for 15 minutes at 4 °C. Protein concentration was determined by bicinchoninic acid (BCA) assay as described above. 300 µg of protein lysate was then incubated with 30 µl of pre-cleared High Capacity Streptavidin Agarose beads (Pierce, 20359) and the mixture was incubated on a rotating wheel for 12 hours at 4 °C. Beads were pelleted by centrifugation at 800 rpm for 5 minutes and then transferred with 1 ml of lysis buffer to a fresh Eppendorf tube. Beads were washed 5 times with washing buffer (50 mM Tris-HCl, 0.5% NP-40) and eluted with 5% SDS.

Elutes were reduced using a final concentration of 10 mM Pierce DTT (A39255, Thermo Fisher Scientific) dissolved in Triethylammonium bicarbonate buffer (TEABC) (T7408, Sigma Aldrich) at 60°C for 30 mins. Then, tubes were incubated to room temperature for 20 mins in dark and diluted to 20 mM iodoacetamide (A39271, Thermo Fisher Scientific) dissolved with TEABC. Afterwards, samples were processed with S-Trap™ mini spin column (PROTIFI, C02-mini) according to manufacteŕs instructions. Finally, tryptic digestion was performed by incubating the processed sample with 10 µg Pierce Trypsin Protease MS-grade (90058, Thermo Fisher Scientific) dissolved in 100 µl 100 mM TEABC overnight at 37 °C. Samples were then eluted from the column and transferred to a fresh Eppendorf tube. Columns were washed first with 0.15% Formic Acid and then with 50% Acetonitrile in 0.15% formic acid and pooled with the eluate. Finally, sample were lyophilized and resuspended in 1% formic acid.

The dried peptides were reconstituted in 1% formic acid and analyzed on an Orbitrap Astral mass spectrometer connected to a Thermo Fisher Scientific Vanquish Neo UHPLC. The peptides were enriched on a trap column and separated on an analytical column (Easy-Spray PepMap Neo C18, 2 μm, 75 μm x 150 mm) at 800 nl/min. Chromatographic separation was performed using a gradient elution: 5% BufferA (0,1% Formic Acid) and 22.5% buffer B [90% ACN, 0,1% Formic Acid] at 14 minutes, ramped to 35% at 21 minutes, followed by a return to 9%. The total run time was 22.6 minutes. Data acquisition was performed in Data-Independent Acquisition (DIA) mode using the Astral analyzer. The MS data acquired in Orbitrap were operated with a fixed cycle time of 5 ms and with a full scan range of 380 to 980 *m*/*z* at a resolution of 240,000. The AGC was set to 500% and ion injection time is custom. Precursor ion selection width was kept at 4-Th and peptide fragmentation was achieved by Higher-energy collisional dissociation (HCD, normalized collision energy 25%). For DIA mode, scan range used 150 to 2000, AGC target was 500%, ion injection was custom, and detector was Astral.

### Quantitative proteomics

For unbiased identification of degrader target proteins, 1 × 10^6^ HEK293 cells per condition were treated with DMSO (1:1,000), Compound 1* (1 µM) or Compound 1 (1 µM) for 12 h in biological triplicates. Cells were collected via centrifugation, washed two times in ice-cold PBS and cellular pellets were lysed in 100 µl of RIPA buffer supplemented with benzonase and cOmplete EDTA-free Protease Inhibitor Cocktail. Cell debris was removed by centrifugation at 13,000 rpm for 15 min at 4 °C. Supernatant was transferred to fresh tubes and protein concentration determined using the BCA protein assay kit. 50 µg of cell lysate was then diluted to a final 5% SDS concentration and proteins were first reduced at 55°C for 15 mins in DTT (A39255, Thermo Fisher Scientific) dissolved in Triethylammonium bicarbonate buffer (TEABC) (T7408, Sigma Aldrich) to a final concentration of 5 mM. Proteins were alkylated with to a final 20 mM iodoacetamide (A39271, Thermo Fisher Scientific) dissolved with TEABC in the dark for 10 min at room temperature. Afterwards, samples were processed with S-Trap™ micro spin column (PROTIFI, C02-micro) according to manufacturer’s instructions. Finally, tryptic digestion was performed by incubating the processed sample with 10 µg Pierce Trypsin Protease MS-grade (90058, Thermo Fisher Scientific) dissolved in 100 µl 100 mM TEABC overnight at 37 °C. Samples were then eluted from the column and transferred to a fresh Eppendorf tube. Columns were washed first with 0.15% Formic Acid and then with 50% Acetonitrile in 0.15% formic acid and pooled with the eluate. Finally, sample were lyophilized and resuspended in 1% formic acid. The sample was analyzed on an Orbitrap Ascend Tribrid mass spectrometer coupled to a Thermo Fisher Scientific Vanquish Neo UHPLC. The peptides were enriched on a trap column and resolved on an analytical column (Easy-Spray PepMap Neo 2 μm C18 75 μm × 150 mm) with 800 nl/min. The gradient for separation was used as 2-7% B at 6 min, 7-18% at 89 min, 18-27% at 114 min, 27-35% at 134 min followed by column wash. Total run time used was 155 min. The data were acquired in data-independent acquisition mode in an orbitrap analyzer. The MS data acquired in Orbitrap were operated with standard AGC and auto ion injection time and with a full scan range of 300 to 1350 *m*/*z* at a resolution of 60,000. For DIA scan precurson mass range were used as 400-1000, precursor was isolated in quadrupole with isolation window 10 Th and peptide fragmentation was achieved by HCD (normalized collision energy 30%). The resolution was set s 15,000.

### Mass spectrometry analysis

Acquired raw files were processed using DIA-NN, version 1.8 (https://github.com/vdemichev/DIaNN). Human UniProt was used as the protein sequence database. Two missed cleavage and a maximum of two variable modifications per peptide were allowed (acetylation of protein N-termini and oxidation of methionine). Carbamidomethylation of cysteines was set as fixed modification. This data analysis was carried out using library-free analysis mode in DIA-NN with “deep learning-based spectra and RTs prediction” enabled (51). MBR was enabled. The search results were further processed in Perseus (52). The data were loaded in Perseus in .txt format. Replicates were grouped based on their annotation to (Compound1, Compound2, Compound1* or DMSO). The data were log₂-transformed and missing values were imputed from a normal distribution using the “Processing → Imputation → Replace missing values from normal distribution” function in Perseus. Imputation parameters included a width of 0.3 and a downshift of 1.8 standard deviations, simulating low-abundance values to replace missing entries. The imputation was applied in ‘whole matrix’ mode. The processed data were subsequently used to generate volcano plots in GraphPad Prism (v9.5.1).

### SMARCA2-Proximity Labelling of DCAF16/FBXO22 with Immunoprecipitation

Flp-In™ T-REx™ 293 miniTurbo-SMARCA2 BD expressing cells were seeded in 10-cm dishes with 1,2 x 10^6 cells. The following day cells were transfected with 5 μg of pRK5 HA-DCAF16 or pCMV5 HA-FBXO22. 33 hours after transfection, miniTurbo-SMARCA2 expression was induced with 1 µg ml^−1^ Doxycycline treatment for a total amount of 15 hours. Afterwards, cells were pretreated for 30 mins with 5 µM MLN4924 and 1 µM Compound1/2 plus 170 µM biotin was added subsequently for 8 and 2 hours respectively. Cells were then washed in PBS and collected by centrifugation. The cell pellet was lysed in RIPA buffer supplemented with benzonase and cOmplete EDTA-free Protease Inhibitor Cocktail. 500 μg of cell lysate were incubated with 4 μg of HA antibody (51064-2-AP, Proteintech) or IgGs control (sc-2026, Santa Cruz) overnight at 4 °C. Antibodies were then purified with 25 μl of protein A/G Magnetic Beads (88802, Thermo Fisher Scientific) for 1 hour. Immunoprecipitated proteins were then washed five times in IP-washing buffer (50 mM TRIS-HCL+1% NP40+1 % SDS) and eluted in 4X LDS sample buffer. Samples were analyzed by immunoblotting.

### Flow-cytometric SMARCA2 reporter assays

KBM7 iCas9 cells were transduced with lentivirus expressing SFFV–SMARCA2-BD-mTagBFP–P2A–mCherry to generate stable reporter cell lines. Cell pools expressing both mTagBFP and mCherry were selected via sorting. To quantify the influence of genetic perturbations on compound-induced reporter degradation, stable reporter cell lines were transduced with lentiviral sgRNA (pLenti-U6-sgRNA-IT-EF1αs-THY1.1-P2A-NeoR) and selected with Neomycin. Cas9 expression was induced with doxycycline (0.4 µg ml^−1^) for 5 days, followed by 24 hours of degrader treatment before flow cytometry analysis on an LSR Fortessa (BD Biosciences) operated on BD FACSDiva software (v9.0). Data analysis was performed in FlowJo v10.10.0. BFP and mCherry mean fluorescence intensity values from Cas9 expressing cells were first normalized by background subtraction of the respective values from reporter negative cells. SMARCA2 abundance was calculated as the ratio of background subtracted BFP to mCherry mean fluorescence intensity and is displayed normalized to DMSO treated cells.

### FACS-based CRISPR–Cas9 SMARCA2 stability reporter screens

The SMARCA2 BD stability reporter FACS-based CRISPR/Cas9 screen, library preparation, next-generation sequencing and data analysis was performed as previously described (53) In brief, doxycycline-inducible Cas9 KBM7 cells stably expressing SMARCA2 BD-mTagBFP–P2A–mCherry were transduced in the presence of 8μg ml^-1^ polybrene with a ubiquitin-proteasome-system-focused (UPS) sgRNA library targeting 1301 ubiquitin-associated human genes with 6 sgRNAs per gene at a multiplicity of infection of 0.15 and over 1000x library representation (26) [Supplementary Table 2]. Cells expressing sgRNAs were selected for 14 days with G418 (1 mg ml^−1^, Sigma-Aldrich, A1720), then Cas9 expression was induced with doxycycline (0.4 μg ml^−1^, PanReac AppliChem, A2951). Three days after Cas9 induction, 50 x 10^6^ cells were treated with DMSO, 0.1 μM Compound 1 or 1 μM Compound 2 for 24 hours, in 2 biological replicates.

After treatment, cells were washed once with 1xPBS, and then incubated/stained with anti-Thy1.1-APC (1:400, BioLegend, 202526), Zombie NIR™ Fixable Viability Dye (1:1000, BioLegend, 423105), and Human TruStain FcX™ Fc Receptor Blocking Solution (1:400, BioLegend, 422301) in FACS buffer (1xPBS, 5% FBS, 1 mM EDTA) for 10 min at 4°C. Following two washes with FACS buffer, cells were fixed with BD CytoFix Fixation Buffer (BD Biosciences, 554655) for 45 minutes at 4°C, protected from light. Fixed cells were then washed again with FACS buffer and stored overnight in FACS buffer at 4°C. The following day, cells were strained trough a 35-µm nylon mesh and sorted using a BD FACSAria™ Fusion (BD Biosciences) equipped with a 70 μm nozzle. Aggregates, dead (Zombie NIR-positive), Cas9-negative (GFP-negative), and sgRNA library-negative (Thy1.1–APC-negative) cells were excluded. The remaining cells were sorted based on their SMARCA2 BD–BFP and mCherry levels into SMARCA2^high^ (∼5% of cells), SMARCA2^mid^ (∼30%) and SMARCA2^low^ (∼5%) fractions, ensuring a minimum library representation of 2000 x per replicate.

For library preparation, genomic DNA from the sorted cell fractions was isolated by cell lysis (10 mM Tris-HCl, 150 mM NaCl, 10 mM EDTA, 0.1% SDS), proteinase K treatment (New England Biolabs, P8107) and DNAse-free RNAse digestion (Thermo Fisher Scientific). DNA was then purified through two rounds of phenol extraction (Sigma Aldrich, P4557) and isopropanol precipitation (Sigma-Aldrich, I9516). Isolated genomic DNA was subjected to a two-step PCR amplification of the sgRNA cassette using AmpliTaq Gold polymerase (Thermo Fisher Scientific, 4311818), with sample-specific barcodes introduced during the first PCR and standard Illumina adapters added in the second PCR. The resulting PCR products were purified using Mag-Bind TotalPure NGS beads (Omega Bio-tek, M1378-00) and the final Illumina libraries were pooled and sequenced on a NovaSeq 6000 platform (Illumina). Screen analysis was performed using the crispr-process-nf Nextflow pipeline (https://github.com/ZuberLab/crispr-process-nf/). Briefly, raw FASTQ files were trimmed with *cutadapt* (v4.4) to remove random barcodes and spacer sequences. Demultiplexing was conducted based on sample barcodes. Reads were aligned to the custom UPS sgRNA library using *Bowtie2* (v2.4.5), and guide abundance was quantified with *featureCounts* (v2.0.1). Final read count tables were generated and subsequently analyzed using the crispr-mageck-nf workflow (https://github.com/ZuberLab/crispr-mageck-nf/) for downstream statistical analysis. Gene-level enrichment was calculated in ***MAGeCK*** (0.5.9) (54) by comparing sorted SMARCA2^high^ or SMARCA2^low^ populations against the SMARCA2^mid^ reference population, using median-normalized read counts and replicate-level variance estimation.

### Deep mutation scanning of DCAF16

To generate the lentiviral DCAF16 DMS library comprised of 4300 DCAF16 single point mutants fused to EGFP-cpHalo, Lenti-X HEK293T cells were seeded in 10cm dishes and transfected at approximately 80% confluency with 4 ug library plasmid, 2 ug psPAX2 (Addgene #12260), and 1 ug pMD2.G (Addgene #12259) using PEI (PolySciences). Viral supernatant was collected after 72h and cleared of cellular debris by filtration through a 0.45 µm polyethersulfone filter.

50 million FBXO22 and DCAF16 double knockout HEK293T cells stably expressing SMARCA2 BD-mTagBFP–P2A–mCherry or BRD4-Tandem-BDs-mTagBFP–P2A–mCherry were transduced in the presence of 8μg ml^-1^ polybrene with the DCAF16 DMS library virus at a multiplicity of infection of 0.16-0.19, yielding a calculated library representation of over 1000 cells per variant. Library-transduced cells were selected with 1μg ml^-1^ Puromycin for 7 days. Approximately 20 million cells for each stability reporter were treated with DMSO, 1 μM Compound 1, 1 μM Compound 2, 1 μM Compound 3, 1 μM MMH2, or 0.1 μM GNE-0011 for 24h, in three biological replicates. Following compound treatment, cells were trypsinized, harvested, washed with PBS, and then fixed with BD CytoFix Fixation Buffer (BD Biosciences, 554655) for 40 minutes at 4°C in the dark. Fixed cells were washed with PBS and stored in FACS buffer (1xPBS, 5% FBS, 1 mM EDTA) at 4°C overnight.

The following day, cells were strained through a 35 μm nylon mesh and sorted on a BD FACSAria™ Fusion (BD Biosciences) operated on BD FACSDiva software (v.8.0.2) and equipped with a 70μm nozzle. Aggregates, dead, reporter negative (BFP and mCherry negative), and library negative (EGFP negative) cells were excluded from the sort. The remaining cells were sorted based on their SMARCA2 BD-BFP or BRD4-Tandem-BDs-BFP and mCherry levels into low and high fractions (∼5 – 15% of cells in each fraction), ensuring a minimum library representation of 1500 x per replicate. Three replicates of 20 million unsorted cells for each stability reporter were also harvested as controls, and directly washed and frozen.

DNA libraries of sorted and unsorted cell pools for Next-generation sequencing (NGS) were prepared as previously described (53). In short, genomic DNA (gDNA) was extracted by cell lysis (10 mM Tris-HCl, 10 mM EDTA, 150 mM NaCl, 0.1% SDS), proteinase K treatment (New England Biolabs, P8107) and DNAse-free RNAse digestion (Thermo Fisher Scientific), followed by two rounds of phenol extraction and isopropanol precipitation. DCAF16 variant cDNAs were amplified via PCR from gDNA using Q5 High-Fidelity DNA Polymerase (New England Biolabs, M0491L) and the following primers: DCAF16_DMS_seq_F (TGGCACAGGAGGTTCAATG) and DCAF16_DMS_seq_R (TCATAATCCGCAGTTCCAGG). PCR reactions for each sample were pooled and purified using Mag-Bind TotalPure NGS beads (Omega Bio-tek, M1378-00). The library preparation from the amplified DNA was performed using the Tagment DNA TDE1 Enzyme and IDT for Illumina Unique Dual Indexes (Illumina). Library concentrations were quantified with the Qubit 2.0 Fluorometric Quantitation system (Life Technologies) and the size distribution was assessed using the 2100 Bioanalyzer instrument (Agilent). For sequencing, samples were diluted and pooled into NGS libraries and sequenced on NovaSeq 6000 instrument (Illumina) following a 100-base-pair, paired-end recipe.

Raw sequencing reads were converted to fastq format with samtools (v1.15.1) and bedtools (v.2.30.0) using bamtofastq function. Sequencing reads were trimmed using Trim Galore (v0.6.6) using nextera and pair modes. Short reads were aligned to the DCAF16 cassette and SAM files were generated using mem algorithm from the bwa software package (v0.7.17). SAM files were converted to BAM using samtools and mutation calling was performed using the AnalyzeSaturationMutagenesis tool from GATK (v4.1.8.1). Given the sequencing strategy, >98% of reads corresponded to wild-type sequences and were filtered out during this step. Next, relative frequencies of variants were calculated for each position and variants that were covered by less than 1 in 30,000 reads were excluded from further analysis. Read counts for each variant were then normalized to the total read count of each sample and log_2_ fold changes were calculated for SMARCA2/BRD4^high^ or SMARCA2/BRD4^low^ sorted fractions over unsorted pools. To correct for differential drug potency, each variant was then normalized to the maximum log_2_ fold change. For drug comparisons, log_2_ fold changes of SMARCA2/BRD4^high^ or SMARCA2/BRD4^low^ over unsorted pools of each drug were subtracted. Heatmaps were generated using pheatmap (v.1.0.12) package in R (v.4.1.0).

### Protein expression and purification

#### DCAF16 and mutants

The coding sequences for full-length DCAF16 with TEV-cleavable N-terminal His_6_-tags were cloned into a pFastBacDual vector under the control of the *polh* promoter. Site directed mutagenesis (SDM) PCR was used to create DCAF16 mutants (C58S, C58A, C173S and C173A). Coding sequences for full-length DDB1 or DDB1(ΔBPB) and full-length DDA1 were cloned into a pFastBacDual vector under the control of *polh* and p10 promoters, respectively. Bacmids was generated using the Bac-to-Bac baculovirus expression system (Thermo Fisher Scientific). Baculovirus was generated by adding bacmid (1 µg ml^−1^culture volume) mixed with 2 µg PEI 25 K (Polysciences) per µg bacmid in 200 µl PBS and incubated at room temperature for 30 min. The mixture was added to a suspension culture of Sf9 cells at 1 × 10^6^ cells per ml in Sf-900 II SFM (Gibco) and incubated at 27 °C with shaking at 110 rpm. Viral supernatant (P0) was collected after 7 days. For expression in *Trichoplusia ni* High Five cells were grown to densities between 1.5 to 2 × 10^6^ cells per ml in Express Five SFM (Gibco) supplemented with 18 mM L-glutamine and infected with a total virus volume of 1% per 1 × 10^6^ cells per ml, consisting of equal volumes of DCAF16 and DDB1 + DDA1 baculoviruses. For expression in SF9 cells 3.5 x 10^6^ cells per ml in SF-900 II SFM (Gibco) and infected with 1% of culture volume of each virus (DCAF16 and DDB1:DDA1). Cells were incubated at 27 °C in 2 l Erlenmeyer flasks (∼500 ml culture per flask) with shaking at 110 rpm for 72 h. Cells were spun at 1,000*g* for 20 min and supernatant was discarded. Pellets were resuspended in lysis buffer (50 mM HEPES, 500 mM NaCl, 1 mM TCEP, pH 7.5, Tween-20 (to 1% (v/v)), magnesium chloride (to 2 mM), benzonase (to 1 µg ml^−1^) and cOmplete EDTA-free Protease Inhibitor Cocktail (Roche, 2 tablets per litre initial culture volume), reuspension was frozen and stored at −80 °C. Resuspended pellets were thawed and refrozen twice to help with cell lysis. Cell suspensions were sonicated, and lysates were centrifuged at 23,000 rpm for 40 min. Clarified lysate was added to 2 ml nickel agarose resin per litre culture on a roller at 4 °C for 1 h. Mixture of resin and lysate was centrifuged at 500g for 2 minutes to separate lysate from resin taking supernatant each time and washing resin with wash buffer 3 times (50 mM HEPES, 500 mM NaCl, 1 mM TCEP, 20 mM imidazole, pH 7.5). Bound protein was eluted with elution buffer (50 mM HEPES, 500 mM NaCl, 1 mM TCEP, 500 mM imidazole, pH 7.5). Eluted protein was added to dialysis bag (Snakeskin 3.5 kDa MWCO, Thermo Fisher Scientific) and TEV protease was added to protein and incubated at 4 °C overnight. The sample was run over nickel agarose resin. Flow-through and washes with were collected and pooled. Protein was buffer exchanged into ion exchange (IEX) buffer A (50 mM HEPES, 50 mM NaCl, 1 mM TCEP, pH 7.5) by 2-hour dialysis at room temperature with a change of dialysis buffer after 1 hour. The sample was then loaded onto a HiTrap Q HP 5 ml column (Cytiva). The column was washed with IEX buffer A and bound protein was eluted with a 0–100% IEX buffer B (50 mM HEPES, 1 M NaCl, 1 mM TCEP, pH 7.5) gradient. Fractions containing protein were pooled and concentrated then run on 16/600 Superdex 200 pg column or 10/300 Superdex 200 Increase (Cytiva) equilibrated in 20 mM HEPES, 150 mM NaCl, 1 mM TCEP, pH 7.5. Fractions containing the purified protein complex were pooled and concentrated then aliquoted and flash frozen in liquid nitrogen for storage at −80 °C.

Prior to intact protein mass spectrometry experiments purified DCAF16-DDB1-DDA1 and mutants were dephosphorylated with 1:40 ratio of *E. coli* His-GST-lambda protein phosphatase (made in house) to DCAF16 protein at 4°C overnight in 20 mM HEPES, 150 mM NaCl, 1 mM TCEP pH 7.5 buffer supplemented with 1 mM MnCl_2_. The next day nickel resin (Abcam) was incubated with the mixture for 1 hour after which it was spun to sediment resin and recover the supernatant containing dephosphorylated DCAF16 and remove His-GST-lambda protein phosphatase that was immobilised on the resin.

#### SMARCA2

SMARCA2 bromodomain (residues 1376-1590 d1403-1420) was PCR amplified and sub-cloned into His-12-SUMO (pRSF-DUET1) using quick ligase with 5’-BamHI and 3’-EcoRI. Plasmid was transformed into *E. coli* BL21 (DE3). Cultures were subjected to overnight expression at 18 °C, induced with 0.25 mM IPTG at OD_600_ ∼ 0.8–1. Cells were collected by centrifugation and pellets were resuspended in lysis buffer 50 mM HEPES pH 7.5, 300 mM NaCl, 1 mM TCEP, 10% glycerol (supplemented with 2 mM magnesium chloride, Benzonase and cOmplete EDTA-free Protease Inhibitor Cocktail (Roche, 1 tablet per litre initial culture volume), resuspended pellet was frozen and stored at −20 °C. Resuspension was thawed and lysed at 30,000 psi using a CF1 Cell Disruptor (Constant Systems). The lysate was cleared by centrifugation at 20,000 rpm for 30 min at 4 °C. The lysate loaded on to a 5 ml HisTrap HP column (Cytiva) equilibrated in lysis buffer and eluted with an imidazole gradient up to 100% elution buffer (50 mM HEPES, 500 mM NaCl, 0.5 mM TCEP, 500 mM imidazole, pH 7.5). Eluted sample was placed into a dialysis bag (Snakeskin 3.5 kDa MWCO, Thermo) and ULP1 was added to sample to cleave the His_12_-SUMO tag and bag was placed into 2 L of lysis buffer to dialyse overnight at 4°C SMARCA2 bromodomain was run on a 5 ml HisTrap HP column equilibrated in lysis buffer. The flow-through and wash containing SMARCA2 bromodomain were pooled and were concentrated in 3,000 MWCO Amicon centrifugal filter units (Merck Millipore). The protein was loaded onto a HiLoad 16/600 Superdex 75 pg column (GE LifeSciences) equilibrated in 20 mM HEPES pH 7.5, 150 mM NaCl, 0.5 mM TCEP. Fractions containing either pure SMARCA2 bromodomain were confirmed by SDS–PAGE, then pooled, concentrated and aliquoted for storage at −80 °C until use. For GST-tagged SMARCA2 BD residues (1376-1590 d1403-1420) was sub-cloned into pDEST15 vectors (Invitrogen)and transformed into E.*coli* BL21 (DE3). Cultures were subjected to overnight expression at 18_o_C, induced with 0.4 mM IPTG at OD600 ∼ 0.8–1. Cells were collected by centrifugation and pellets were resuspended in lysis buffer 50 mM HEPES pH 7.5, 300 mM NaCl, 1 mM TCEP, 10% glycerol (supplemented with 2 mM magnesium chloride, Benzonase and complete EDTA-free Protease Inhibitor Cocktail (Roche, 1 tablet per litre initial culture volume), resuspended pellet was frozen and stored at −20 °C. Resuspension was thawed and lysed at 30,000 psi using a CF1 Cell Disruptor (Constant Systems). The lysate was cleared by centrifugation at 20,000 rpm for 30 min at 4 °C. The lysate loaded on to a 20 mL of Glutathione affinity resin (Abcam) equilibrated in lysis buffer and incubated on a roller for 3 hours at room temperature. Resin was washed by multiple rounds of adding fresh lysis buffer. Elution buffer (50 mM HEPES pH 7.5, 300 mM NaCl, 1 mM TCEP, 10% glycerol and 25mM L-glutathione) was added GST-SMARCA2 BD bound GST-resin and incubated for 3 hours on a roller at room temperature. Elution was collected and dialysed into 50 mM HEPES pH 7.5, 50 mM NaCl, 1 mM TCEP, 10% glycerol and the subjected to 5mL Hitrap Q and eluted on a 50 mM-1mM NaCl gradient. GST-SMARCA2 contain fractions were loaded onto a HiLoad 16/600 Superdex 75pg column (GE LifeSciences) equilibrated in 20 mM HEPES pH 7.5, 300 mM NaCl, 0.5 mM TCEP.

#### FBXO22-SKP1

His_10_-SUMO-SKP1-(GGS)x4-FBXO22 were Gibson assembled into a pRSF-DUET1 plasmid. Protein expression was performed in BL-21 Rosetta (DE3) *E.coli* and overnight expression at 18 °C was induced with 0.25 mM IPTG at OD_600_ ∼ 0.8–1. Cells were collected by centrifugation and pellets were resuspended in lysis 25 mM HEPES pH 7.5, 250 mM NaCl, 2.5 mM TCEP, 20 mM imidazole (supplemented with 2 mM magnesium chloride, Benzonase and cOmplete EDTA-free Protease Inhibitor Cocktail (Roche, 1 tablet per litre initial culture volume), resuspended pellet was frozen and stored at −20 °C. Resuspension was thawed and lysed at 30,000 psi using a CF1 Cell Disruptor (Constant Systems). The lysate was cleared by centrifugation at 20,000 rpm for 30 min at 4 °C. Clarified lysate was added to 2 ml nickel agarose resin (Abcam) per litre culture on a roller at 4 °C for 1 h. Mixture of resin and lysate was centrifuged at 500g for 2 minutes to separate lysate from resin taking supernatant each time and washing resin with wash buffer 3 times (50 mM HEPES, 500 mM NaCl, 1 mM TCEP, 20 mM imidazole, pH 7.5). An additional buffer wash was added with buffer supplemented with MgCl2 and ATP to remove heat-shock protein. Washed resin was resuspended in wash buffer and ULP1 was added to cleave His_10_-SUMO tag and left to incubate overnight at 4°C. The resin was placed into a Poly-Prep chromatography column (Biorad) and flowthrough and washes containing cleaved FBXO22-SKP1 were collected and pooled. Pooled protein was added to dialysis bag (Snakeskin 3.5 kDa MWCO, Thermo) dialysed into ion exchange (IEX) buffer A (50 mM HEPES, 100 mM NaCl, 1 mM TCEP, pH 7.5) by 2 hours dialysis at room temperature with a change of dialysis buffer after 1 hour. The sample was then loaded onto a HiTrap Q HP 5 ml column (Cytiva). The column was washed with IEX buffer A and bound protein was eluted with a 0–100% IEX buffer B (50 mM HEPES, 1 M NaCl, 1 mM TCEP, pH 7.5) gradient. Fractions containing protein were pooled and concentrated then run on 10/300 Superdex 200 Increase (Cytiva) equilibrated in 25 mM HEPES, 150 mM NaCl, 1 mM TCEP, pH 7.5. Fractions containing the purified protein complex were pooled and concentrated then aliquoted and flash frozen in liquid nitrogen for storage at −80 °C.

### BRD4 bromodomain 2

BRD4 BD2 (residues 333-460) was expressed in *E. coli* BL21(DE3) and purified as described previously (55). In brief, proteins were purified by nickel affinity chromatography and SEC. His_6_ tag cleavage and reverse nickel affinity was performed prior to SEC for some applications, for others the tag was left on. Purified proteins in 20 mM HEPES, 150 mM sodium chloride, 1 mM DTT, pH 7.5 were aliquoted and flash frozen in liquid nitrogen and stored at −80 °C.

### Intact protein mass spectrometry

Prior to analysis, 10 µM recombinant human dephosphorylated DCAF16:DDB1:DDA1 incubated with 20 µM DMSO or compounds and co-incubated with 20 µM recombinant SMARCA2 BD for 18 h at room temperature. Buffer 25 mM HEPES pH7.5, 150 mM NaCl, 1 mM TCEP. Samples were injected on a column (ZORBAX 300SB-C3), desalted for 1 minute and then eluted to an UHPLC Agilent 1290 Infinity III High-Throughput System (Agilent) using a gradient of 10% MeCN to 95% in 0.1 TFA/water, fragmentor voltage was 135 V and the Capillary voltage was 4000 V. The mass spectrometer acquired full scan mass spectra (*m*/*z) 600-3000* on InfinityLab Pro iQ Series Mass Detect. Mass spectra were deconvoluted using Agilent OpenLab CDS. Labelling efficiency for all intact mass analysis was calculated from relative abundance of the peaks.

### Cryo-EM sample preparation

Prior to plunge freezing, 20µM DDB1(ΔBPB)/DCAF16, 50µM SMARC2^BRD^, and 60 µM GNE-58 were incubated at ambient temperature for 1 and half hours. The complex diluted five-fold into 20 mM HEPES, 150 mM NaCl, 1mM TCEP, pH7.5. Prior to application on the grids, Lauryl Maltose Neopentyl Glycol (LMNG, Anatrace cat# NG310) was added to a final concentration of 0.01% (w/v). 300 mesh copper Quantifoil R1.2/1.3 grids were glow discharged for 60 sec under vacuum at 30 mA using a Quorum SC7620 5 minutes before use. A total of 3.5 μL of the complex with 0.1% LMNG was applied to the carbon side only with final concentration: 4µM DDB1(ΔBPB)/DCAF16, 10µM SMARC2^BRD^, and 12 µM GNE-58. A Vitrobot Mark IV (FEI) operating at 4°C, 100% humidity, wait time = 10 seconds, blot force = 4, blot time = 4 sec was used for blotting with grade 595 filter paper (Agar Scientific cat#AG47000) before plunging into liquid ethane.

### Cryo-EM data acquisition

All grids were clipped and screened at the University of Dundee Electron Microscopy Facility using the Glacios (Thermo Fisher Scientific) operating at 200 kV equipped with a Falcon4i direct electron detector and Selectris energy filter. The data set for DDB1(ΔBPB)/DCAF16, SMARC2^BRD^, GNE-58 was collected at eBIC (Diamond Light Source, UK) under BAG access BI-31827-25 using Krios-II (m03) operating a Schottky X-FEG at 300 kV and Ametek-Gatan BioQuantum K3 Imaging Filter with slit width of 20 eV. Data were recorded using single particle EPU v3.8 (Thermo Fisher Scientific) with aberration-free image shift (AFIS) at a nominal magnification of 165,000x, calibrated pixel size of 0.513 Å, C2 aperture 50 μm, and objective aperture of 100 μm. All movies were collected using a Gatan K3 direct electron detector (5,760×4092 pixels) in counting mode over 60 dose fractions, recorded as Tiff LZW non-gain normalized, with total dose of 60 e⁻/Å², exposure time of 1.26 seconds, and dose rate of 15.7 e/px/s. In the first instance, single particle EPU was used to collect 4,336 movies at 0-degree alpha tilt. For the high tilt data set, alpha tilt was set to 30 degrees in single particle EPU, and 6,550 movies were collected under identical settings.

### Single particle cryo-EM data processing

Full movies and corresponding meta data were separately imported to CryoSparc (56) v4.5 for the 0-degree tilted and the 30-degree tilted data set. Each data set was subjected to patch motion correction and patch CTF estimation, then manually curated with CTF estimation cut at 5 Å. For the 0-degree dataset micrographs were denoised in CryoSparc using 300 micrographs for training over 1200 epochs. Unbiased particle picking was carried out with blob picker on denoised micrographs with a circular blob (minimum diameter of 80 Å and maximum diameter of 160 Å). The blob picked particles were extracted with a box size of 512 pixels, Fourier cropped to 128 pixels (pixel size = 2.05 Å /4x bin) and subjected to 2D classification to remove obvious junk. In the first round, the blob-picked particles were input into ab-initio reconstruction with 8 classes followed by heterogeneous refinement. Many classes contained free DDB1(ΔBPB) or distorted volumes, but one class did contain the complex of DDB1(ΔBPB)/DCAF16 and SMARCA2 BD. Particles for the complex were subjected to 2D classification to remove obvious junk particles, rebalanced by orientations, and a subset of ∼8k particles were taken for training in TOPAZ (v0.2.5a)(57). The trained TOPAZ models were used to extract particles from their respective micrographs, extracted 512 pixels at a pixel size of 2.05 Å. The particles were subjected to ab initio reconstruction with 6 classes. For the 30-degree data set following manual curation a template picker using a good volume of the complex from a previous data collection was employed and picks were inspected and subjected to 2D classification, where junk and free DDB1(ΔBPB) complexes were removed. Selected particles were taken for training in TOPAZ. The trained TOPAZ models were used to extract particles from their respective micrographs, extracted 512 pixels at a pixel size of 2.05 Å. The particles were subjected to ab initio reconstruction with 4 classes. Particles for the complex were subjected to 2D classification to remove junk, rebalanced by orientations, and a subset of particles were taken for further training in TOPAZ. The particles were subjected to 2D classification, Ab initio (6 classes) and non-uniform refinement of best class. Particles sets were created from this and a further round of TOPAZ was run followed by Ab initio into 6 classes. At this point particles from Ab inito containing the complex were extracted at their full box size, 612 pixels (0.513 Å) in separate extract jobs. The extracted particles from each data set were combined for Ab initio reconstruction in 4 classes, followed by heterogeneous refinement. Some junk classes with distorted volumes or incomplete complexes were present, but a clear volume of the complex emerged and was taken for non-uniform refinement. From the best Ab inito class duplicate particles were removed. Particles were subjected to Global CTF refinement followed by a non-uniform refinement, reconstruction and 3D classification into 5 classes. The best class was reconstructed and subjected to further classification (4 classes). The best volume from the 3D classification was imported to UCSF ChimeraX (version 1.8), gaussian blurred, segmented, resampled, and an mrc volume encompassing SMARCA2 and part of DCAF16 was exported to CryoSparc. The mask was binarized, soft padded, and dilated in CryoSparc prior to local refinement. Final refinement contained 30,549 particles from the 0-degree data set and 10,271 particles from the 30-degree data set. The full processing workflow, GSFSC curve, local resolution estimation, angular distribution plot, and posterior position directional distribution plot are presented in Extended Data Figures 4 and 5

### Cryo-EM model building and refinement

DCAF16:DDB1(ΔBPB):DDA1 was extracted from PDB entry 8G46 (34) and SMARCA2-BD was derived using sequence in Alphafold2 (58) server from ChimeraX. They were manually placed into the map in ChimeraX using Fit in Map tool, followed by visual inspection of residues in COOT (version 0.9.8.93). Initial restraints for Compound1 were generated using a SMILES string with eLBOW (in Phenix 1.21-5207) (59).To help guide proper placement of **1** into density, coordinates from PDB ID 8QJT (60) were used as it contained a similar SMARCA^BD^ binder. To create covalent bond, a bespoke restraints file needed to be created and refined in Phenix (linux command line version 1.21.1-5286). After initial fitting it was apparent that there was density visible at the back of DCAF16 that has been previously unresolved in PDB ID 8G46 (34). Therefore, the chain was built using alanine residues and then replaced with correct side chains. The structure was refined with rounds of model building in Coot and refinement with Phenix (v1.20.1-4487) real-space refinement. Validation was done in Phenix. Figures were generated in ChimeraX and PyMOL (version 3.0.2). Electrostatic potential map was generated by using PDB2PQR and APBS server and visualising data in ChimeraX (61).

### Cy5 NHS ester labelling

For DCAF16-DDB1-DDA1 and FBXO22-SKP1 labelling, Alexa Fluor 647 Cy5 NHS ester (Thermo Fisher Scientific) in DMF was prepared to a final concentration of 1 mg/ml with a final concentration of DCAF16–DDB1–DDA1/ FBXO22-SKP1 at 10 mg/ml and sodium bicarbonate 100 mM, diluted in buffer 25 mM HEPES pH 8, 150 mM NaCl, 1 mM TCEP. The solutions were protected from light and incubated for 1 h at room temperature. The solutions run on a Superdex 200 10/300 GL column (Cytiva) to remove free dye and aggregated protein. Fractions containing the Cy5-labelled protein were pooled and concentrated, the degree of labelling was calculated to be greater than 100% for each batch of labelled protein. Labelled protein was aliquoted then flash frozen in liquid nitrogen and stored at −80 °C.

### TR-FRET proximity assay

Stock solutions of reaction components including Cy5-labelled DCAF16–DDB1–DDA1, Cy5-labelled FBXO22, GST-SMARCA2 BD, compounds and LanthaScreen^TM^ Eu-Anti-GST donor (Thermo Fisher Scientific) were prepared in TR-FRET assay buffer (50 mM HEPES pH 7.5, 150 mM NaCl, 1 mM TCEP, 0.05% Tween-20). In this assay the compound was titrated into protein to determine ternary complex-formation. Compounds were titrated 1 in 4 into 125 nM GST-SMARCA2 BD and 500 nM Cy5-DCAF16 or Cy5-FBXO22 to a Corning 3820 low volume 384-well flat bottom plate (black) to a final well volume of 20 μl. Anti-GST donor and DMSO concentrations were kept constant across the plate for both assay formats at 2 nM and 1%, respectively. Background subtraction was performed with using the condition containing Cy5-labelled DCAF16 or FBXO22 and GST-SMARCA2 BD but no compound. Components were mixed by spinning down plates at 50 *g* for 1 min and plates were covered and incubated at room temperature for 30 minutes. Plates were read on a PHERAstar FS (BMG LABTECH) with fluorescence excitation and dual emission wavelengths of 337 and 620/665 nm, respectively, with an integration time between 70 and 400 μs. Data were processed in GraphPad Prism (v10.4.1) curve was fit to a non-linear fit (log (agonist) vs response) with variable slope (four parameters).

### Glutathione reactivity assay

A 1 mM solution of L-glutathione (Sigma-Aldrich, G4251) was prepared in DPBS (Gibco™, 14287080). Each test compound (2 μL, 10 mM in DMSO) was mixed with 98 μL of the L-glutathione solution and incubated at 37 °C for 3 hours. The UV ratio of the test compound to the glutathione adduct was then measured using LC-MS.

### Molecular dynamics (MD)

The cryo-EM structure of the ternary complex between compound **1**, SMARCA2, and DCAF16 was pre-treated using the Protein Preparation Workflow (PPW) available in Schrödinger Suite (Release 2024-3) and Maestro before initiating the MD simulation experiments. The PPW treatment involved preparing the structure at pH 7.5, by adding hydrogen atoms, and building the missing side chain atoms. The hydrogen bond network was optimized by reorienting hydroxyl and thiol groups, water molecules, amide groups of asparagine (Asn) and glutamine (Gln), and the imidazole ring in histidine (His); and predicting protonation states of histidine, aspartic acid (Asp) and glutamic acid (Glu) and tautomeric states of histidine.

Simulations were carried out using the Desmond Molecular Dynamics System implemented within the Schrödinger Suite Release 2024-3 (Schrödinger, LLC). The systems for MD simulations were built being placed in an orthorhombic box (with buffer distance of 12 Å) of explicit water molecules (SPC water model). A salt concentration of 0.15 M was maintained and ions (Na+, Cl-) were placed to neutralize the systems. The built systems were relaxed before the production run in three sequential steps to relieve strain and optimize geometry. In the first step, all atoms in the proteins and the ligand were constrained. In the second step, only the Cα atoms of the proteins were constrained. In the final step, no positional constraints were applied. This stepwise approach allowed gradual relaxation of the system while maintaining the overall structural integrity.

Production MD simulations were carried out in the NPT ensemble (300 K, 1 bar) using Desmond default settings, Nosé-Hoover chain thermostat (relaxation time 1 ps), Martyna-Tobias-Klein barostat (2 ps), and a multi-time-step RESPA integrator (2 fs for bonded and short-range, 6 fs for long-range interactions). Long-range electrostatics employed the u-series decomposition with a 9 Å real-space cut-off. Two independent 500 ns production runs were started from randomised velocities. Coordinates and energies were saved every 500 ps (1 000 frames per run). All analyses reported in this study are based on these production trajectories.

